# Lower promoter activity of the *ST8SIA2* gene has been favored in evolving human collective brains

**DOI:** 10.1101/2021.04.11.439101

**Authors:** Toshiyuki Hayakawa, Masahiro Terahara, Naoko T. Fujito, Takumi Matsunaga, Kosuke Teshima, Masaya Hane, Ken Kitajima, Chihiro Sato, Naoyuki Takahata, Yoko Satta

**Author notes:** Address correspondence to, Toshiyuki Hayakawa, Kyushu University, 744 Motooka, Nishi-ku, Fukuoka 819-0395, Japan.

## Abstract

ST8SIA2 is the main factor regulating expression of the phenotype involved in schizophrenia. Lowered promoter activity of the *ST8SIA2* gene is considered to be protective against schizophrenia by conferring tolerance to psychosocial stress. Here, we examined the promoter type composition of anatomically modern humans (AMHs) and archaic humans (AHs; Neanderthals and Denisovans), and compared the promoter activity at the population level (population promoter activity; PPA) between them. In AMHs, the TCT-type, showing the second lowest promoter activity, was most prevalent in the ancestral population of non-Africans. However, the detection of only the CGT-type from AH samples and recombination tracts in AH sequences showed that the CGT- and TGT-types, exhibiting the two highest promoter activities, were common in AH populations. Furthermore, interspecies gene flow occurred into AMHs from AHs and into Denisovans from Neanderthals, influencing promoter-type compositions independently in both AMHs and AHs. The difference of promoter-type composition makes PPA unique in each population. East and Southeast Asian populations show the lowest PPA. This results from the selective increase of the CGC-type, showing the lowest promoter activity, in these populations. Every non-African population shows significantly lower PPA than African populations, resulting from the TCT-type having the highest prevalence in the ancestral population of non-Africans. In addition, PPA reduction is also found among subpopulations within Africa via a slight increase of the TCT-type. These findings indicate a trend toward lower PPA in the spread of AMHs, interpreted as a continuous adaptation to psychosocial stress arising in migration. This trend is considered as genetic tuning for the evolution of collective brains. The inferred promoter-type composition of AHs differed markedly from that of AMHs, resulting in higher PPA in AHs than in AMHs. This suggests that the trend toward lower PPA is a unique feature in AMH spread.

## Introduction

*Homo sapiens*, anatomically modern humans (AMHs), emerged in Africa and expanded its habitat to almost every corner of the globe during the past 100 thousand years (kyr) [1]. During this spread across the globe, AMHs generated greater technological sophistication and larger bodies of adaptive knowhow to survive efficiently in new environments [2]. Once individuals evolve to learn from one other with sufficient accuracy (fidelity), social groups of individuals develop what might be called *collective brains* [2, 3]. Collective brains, in which individuals are regarded as neurons, generate technological sophistication and adaptive knowhow, and drive cumulative cultural evolution [2, 3].

The *ST8SIA2* gene encodes a sialyltransferase that synthesizes a linear homopolymer of sialic acid (polysialic acid; PSA) in the brain [4]. Sialic acids are a family of nine-carbon monosaccharides that are found at the outer end of glycan chains on the cell surface and are secreted molecules in the deuterostome lineage; they play important roles as ligands in cell–cell communication [5]. The major carrier of PSA is neural cell adhesion molecule (NCAM), and polysialylated NCAM (PSA-NCAM) plays an important role in neurite outgrowth, synapse formation and plasticity [4]. Thus, ST8SIA2 contributes to these neuronal events in the brain by producing PSA.

Three SNPs (rs3759916, rs3759915, and rs3759914) in the promoter region of the *ST8SIA2* gene are associated with schizophrenia risk and involved in its promoter activity [6–9]. Since this association was detected in multiple populations including Asians and Europeans [6–9], these three SNPs can be regarded as a general regulator of the expression of the phenotype involved in schizophrenia. We previously reported that a non-risk haplotype of these three promoter SNPs [CGC-type (C in rs3759916, G in rs3759915, C in rs3759914)] has been under positive selection mainly in Asia during the past ~20–30 kyr [9]. Schizophrenia is an important mental disease because of the serious social impairment that it produces. Since environmental risk factors interact with genetic risk factors toward the onset of schizophrenia [10], the selective pressure in positive selection on the non-risk haplotype (i.e., CGC-type) should be one of the environmental risk factors. Considering that the *ST8SIA2* gene participates in brain functions, psychosocial stress, the sole major environmental risk factor that brain functions are responsible for dealing with [11–15], can be regarded as the selective pressure. Ultimately, positive selection on the CGC-type is considered as an adaptation to the environment exerting psychosocial stress [9]. Therefore, ST8SIA2 is also a master regulator in the expression of the phenotype involved in psychosocial stress.

The CGC-type shows the lowest promoter activity among the four major promoter types (the rest are TGT-, TCT-, and CGT-types), suggesting that the lowered promoter activity of the *ST8SIA2* gene has been favored in adaptation to the environment causing psychosocial stress [9]. This is also supported by the finding that the TCT-type showing the second lowest promoter activity is identified as a haplotype not conferring a risk of schizophrenia in the population in which the frequency of the CGC-type is very low [8]. In addition to the CGC-type, the TCT-type can therefore be regarded as conferring tolerance of psychosocial stress. Overcoming psychosocial stress by increasing the frequency of *ST8SIA2* promoter haplotypes conferring tolerance to such stress (i.e., tolerant types) at the population level would make collective brains bigger, associated with increases in the degree of social interaction and the population size. To elucidate the involvement of the *ST8SIA2* gene in the evolution of collective brains, we examined the promoter-type composition not only in AMHs but also in archaic humans (AHs), and compared its functional consequence between these humans.

## Materials and Methods

### DNA sequence data

Approximately 1 Mb of DNA sequences spanning the three promoter SNPs were retrieved from phase 3 of the 1000 Genomes Project database [16], covering 2,504 individuals (5,008 sequences). We classified them into five meta-populations: Africa (AFR), Europe (EUR), South Asia (SAS), East and Southeast Asia (EAS), and America (AMR) [216 for YRI (Yoruba in Ibadan, Nigeria), 198 for LWK (Luhya in Webuye, Kenya), 226 for GWD (Gambian in Western divisions in the Gambia), 170 for MSL (Mende in Sierra Leone), 198 for ESN (Esan in Nigeria), 122 for ASW (Americans of African ancestry in Southwest USA), and 192 for ACB (African Caribbeans in Barbados) in AFR (in total 1322); 198 for CEU (Utah residents with Northern and Western European ancestry), 214 for TSI (Toscani in Italy), 198 for FIN (Finnish in Finland), 182 for GBR (British in England and Scotland), and 214 for IBS (Iberian population in Spain) in EUR (in total 1006); 206 for GIH (Gujarati Indian from Houston, Texas, USA), 192 for PJL (Punjabi from Lahore, Pakistan), 172 for BEB (Bengali from Bangladesh), 204 for STU (Sri Lankan Tamil from the UK), and 204 for ITU (Indian Telugu from the UK) in SAS (in total 978); 206 for CHB (Han Chinese in Beijing, China), 208 for JPT (Japanese in Tokyo, Japan), 210 for CHS (Southern Han Chinese, China), 186 for CDX (Chinese Dai in Xishuangbanna, China), and 198 for KHV (Kinh in Ho Chi Minh City, Vietnam) in EAS (in total 1008); and 128 for MXL (Mexican ancestry in Los Angeles, California), 208 for PUR (Puerto Rican in Puerto Rico), 188 for CLM (Colombian in Medellin, Colombia), and 170 for PEL (Peruvian in Lima, Peru) in AMR (in total 694)] [9]. This sequence dataset is hereafter abbreviated as *D*_1000_.

DNA sequences of AHs [three Neanderthals (Vindija, Altai, and Chagyrskaya) and one Denisovan] were obtained from the public database (http://cdna.eva.mpg.de/neandertal/Vindija/; http://cdna.eva.mpg.de/neandertal/altai/; http://ftp.eva.mpg.de/neandertal/Chagyrskaya/VCF/; http://cdna.eva.mpg.de/neandertal/denisova/). The chimpanzee sequence was obtained from the chimpanzee genome database (Clint_PTRv2/panTro6; https://www.ncbi.nlm.nih.gov/genome), and used as an out-group in tree constructions.

### Schizophrenia-associated genetic loci

To examine the uniqueness of the promoter-type distribution of the *ST8SIA2* gene, we compared it with the allele frequencies of other schizophrenia-associated genetic loci identified from European and East Asian populations [17, 18]. A total of 308 index SNPs associated with schizophrenia risk were chosen because these are expected to link strongly to functional SNPs directly involved in schizophrenia risk.

### Construction of haplotype sequences of archaic humans

The three promoter SNPs are located in an 18-kb region that is sandwiched in between recombination hot spots [9]. We therefore focused on this 18-kb region because the use of a longer region would cause ambiguity in the construction of archaic haplotypes by breaking the linkage of SNPs. The 18-kb region is homozygous in Altai Neanderthal (Supplementary Fig. 1). Based on the linkage in the Altai Neanderthal sequence, we decided on the linkage of SNP sites in other AH sequences (Supplementary Fig. 1).

### Haplotype analysis

A phylogenetic tree of haplotype sequences was constructed using the Neighbor-Joining method [19]. Genetic distances were calculated with multiple hit corrections with MEGAX software [20]. By following the work of Fujito et al. (2018) [9], homozygote tract lengths (HTLs) were obtained.

### Detection of introgressed fragments

To detect introgressed fragments, we applied the Sprime method [21].

### Ages of variants

Ages of variants were obtained from a website (https://human.genome.dating/; [22]).

### Examination of archaic origin of promoter type not found in the AH database

To examine the archaic introgression of promoter types not found in the AH sequence database, we focused on the population that is considered as a direct descendant of the population in which the archaic introgression occurred, and used the information of the site differences between AMHs and AHs, the number of private haplotypes, recombination with other promoter types, and haplotypes shared with Africans. Based on these, we extracted candidates of descendant haplotypes of the archaic ancestry by following five steps.

Step 1. The number of site differences between AMHs and AHs (AMH–AH site differences) is obtained from comparison of the sequence between AMHs and AHs by ignoring recombinants. The pairwise comparison of site differences among haplotypes is performed to make a site difference matrix.
Step 2: Haplotypes showing the AMH–AH site differences in the matrix are selected for step 3 because these divergences correspond to the AMH–AH divergence.
Step 3: Phylogenetic trees of selected haplotypes are constructed. In the trees, it is expected that haplotypes can be divided into two groups corresponding to AMHs and AHs, if archaic introgression occurred.
Step 4: If haplotypes are distributed widely in AFR, such haplotypes and close relatives whose site differences with them are less than the AMH–AH site differences are rejected.
Step 5: If there are alleles shared only by the remaining haplotypes, the ages of such variants are examined. If not, the ages of variants uniquely found in the remaining haplotypes are examined. If the ages are not close to the time of archaic introgression, such haplotypes are rejected.

### Promoter activity at the population level

Since the *ST8SIA2* gene is not an imprinting gene, it is assumed that the two promoter-type alleles contribute equally to the gene expression in individuals. Each promoter type has unique promoter activity [9, 23], which was measured using the region whose sequence is the same as that from the most abundant haplotype of each promoter type (see Supplementary Tables 1–3; [9]). Based on the relative difference in the promoter activity compared with that of the CGC-type, we calculated the average of the relative differences (*a*_1_ and *a*_2_) of two promoter-type alleles as the individual promoter activity [(*a*_1_ + *a*_2_)/2], and then obtained the promoter activity at the population level (population promoter activity; PPA) as the average of individual promoter activities in the population by

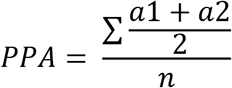

where *n* is the number of individuals in the population.

## Results

### Distribution of promoter types

In the present-day human population, over 99% of promoter types are the TGT-, TCT-, CGT-, and CGC-types ([9]; Table 1). The TGT-type is the ancestral type [9] and is distributed globally. Its frequency is lower in non-AFR than in AFR [20.2% (1013/5008) in total, 46.3% (612/1322) in AFR, 3.3% (33/1006) in EUR, 11.1% (109/978) in SAS, 13.2% (133/1008) in EAS, and 18.2% (126/694) in AMR] (Table 1; Supplementary Fig. 2A). In contrast to the TGT-type, the TCT-type is most prevalent in non-AFR, especially in EUR [67.3% (3369/5008) in total, 47.3% (625/1322) in AFR, 96.3% (969/1006) in EUR, 80.0% (782/978) in SAS, 51.1% (515/1008) in EAS, and 68.9% (478/694) in AMR] (Table 1; Supplementary Fig. 2B). The CGT-type is the rarest among the four promoter types [1.5% (74/5008) in total, 3.5% (46/1322) in AFR, 0.1% (1/1006) in EUR, 1.1% (11/978) in SAS, 0.8% (8/1008) in EAS, and 1.2% (8/694) in AMR], and its frequency is lower outside of Africa (Table 1; Supplementary Fig. 2C). The CGC-type is relatively common in SAS, EAS, and AMR [10.4% (520/5008) in total, 0.8% (11/1322) in AFR, 0.2% (2/1006) in EUR, 7.8% (76/978) in SAS, 34.6% (349/1008) in EAS, and 11.8% (82/694) in AMR] (Table 1; Supplementary Fig. 2D). We previously reported that this distribution of the CGC-type has been influenced by positive selection [9].

**Table 1.** Distribution of promoter types

| Promoter type | AFR | EUR | SAS | EAS | AMR | Total |
| --- | --- | --- | --- | --- | --- | --- |
| TGT | 612 | 33 | 109 | 133 | 126 | 1013 |
| TCT | 625 | 969 | 782 | 515 | 478 | 3369 |
| CGT | 46 | 1 | 11 | 8 | 8 | 74 |
| CGC | 11 | 2 | 76 | 349 | 82 | 520 |
| Others | 28 | 1 | 0 | 3 | 0 | 32 |
| Total | 1322 | 1006 | 978 | 1008 | 694 | 5008 |

### Composition of haplotypes in meta-populations

Since the 18-kb region containing the three promoter SNPs is sandwiched between recombination hot spots [9], we focused on it to examine the haplotype composition using SNPs whose minor allele frequency (MAF) is >1%. Eighty-one TGT haplotypes were identified from *D*_1000_ (Supplementary Table 1). The number of haplotypes is 58, 7, 18, 7, and 25, for AFR, EUR, SAS, EAS, and AMR, respectively. In AFR, over half of TGT sequences are occupied by six haplotypes (HG01271.0, HG00728.0, HG00553.0, HG01886.1, HG01241.0, and HG01110.0) (Table 2). In contrast, HG00132.1, which is minor in AFR (2%), occupies over half of haplotypes in every non-AFR meta-population (Table 2).

**Table 2.**
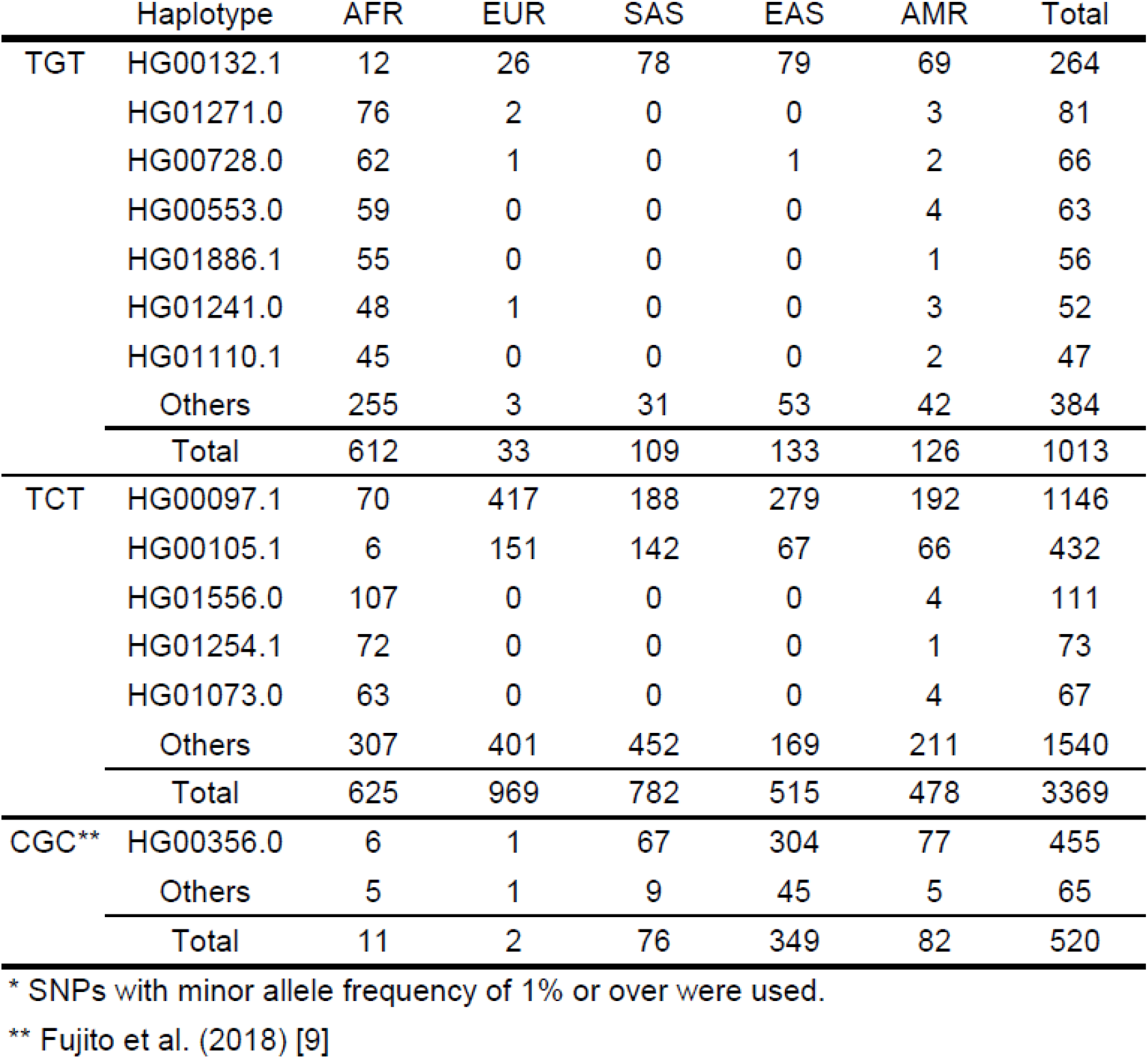
Haplotypes* of the TGT-, TCT-, and CGC-types

|  | Haplotype | AFR | EUR | SAS | EAS | AMR | Total |
| --- | --- | --- | --- | --- | --- | --- | --- |
| TGT | HG00132.1 | 12 | 26 | 78 | 79 | 69 | 264 |
|  | HG01271.0 | 76 | 2 | 0 | 0 | 3 | 81 |
|  | HG00728.0 | 62 | 1 | 0 | 1 | 2 | 66 |
|  | HG00553.0 | 59 | 0 | 0 | 0 | 4 | 63 |
|  | HG01886.1 | 55 | 0 | 0 | 0 | 1 | 56 |
|  | HG01241.0 | 48 | 1 | 0 | 0 | 3 | 52 |
|  | HG01110.1 | 45 | 0 | 0 | 0 | 2 | 47 |
|  | Others | 255 | 3 | 31 | 53 | 42 | 384 |
|  | Total | 612 | 33 | 109 | 133 | 126 | 1013 |
| TCT | HG00097.1 | 70 | 417 | 188 | 279 | 192 | 1146 |
|  | HG00105.1 | 6 | 151 | 142 | 67 | 66 | 432 |
|  | HG01556.0 | 107 | 0 | 0 | 0 | 4 | 111 |
|  | HG01254.1 | 72 | 0 | 0 | 0 | 1 | 73 |
|  | HG01073.0 | 63 | 0 | 0 | 0 | 4 | 67 |
|  | Others | 307 | 401 | 452 | 169 | 211 | 1540 |
|  | Total | 625 | 969 | 782 | 515 | 478 | 3369 |
| CGC** | HG00356.0 | 6 | 1 | 67 | 304 | 77 | 455 |
|  | Others | 5 | 1 | 9 | 45 | 5 | 65 |
|  | Total | 11 | 2 | 76 | 349 | 82 | 520 |

\* SNPs with minor allele frequency of 1% or over were used.
\*\* Fujito et al. (2018) [9]

The TCT-type sequences in *D*_1000_ are composed of 163 haplotypes (Supplementary Table 2). AFR, EUR, SAS, EAS, and AMR contain 62, 62, 58, 31, and 51 haplotypes, respectively. In AFR, HG01556.0 is most abundant and occupied around half of sequences with three haplotypes (HG00097.1, HG01254.1, and HG01073.0) (Table 2). In contrast, HG00097.1 is the most prevalent type in every non-AFR meta-population and occupies around half of sequences with or without HG00105.1, which are rare in AFR (<1%), in each non-AFR meta-population (Table 2).

Compared with the TGT- and TCT-types, the number of haplotypes is smaller in both CGC- and CGT-types (31 CGC haplotypes and 20 CGT haplotypes; [9]; Table 3; Supplementary Table 3). As for the CGC-type, AFR, EUR, SAS, EAS, and AMR contain 3, 2, 7, 21, and 5 haplotypes, respectively [9]. The excess in EAS can be explained by the increase of CGC sequences due to positive selection [9]. HG00356.0 is most prevalent in every meta-population [9]. As for the CGT-type, the number of haplotypes is 15, 1, 4, 1, and 5 in AFR, EUR, SAS, EAS, and AMR, respectively (Supplementary Table 3). Two haplotypes (HG01073.1 and HG03667.1) are shared by AFR and non-AFR. However, almost all HG01073.1 sequences are found in AFR (12/13), and the distribution of HG03667.1 is biased to non-AFR (14/16) (Table 3; Supplementary Table 3). In the remaining 18 CGT haplotypes, 13 haplotypes are found only in AFR and the others are identified only in non-AFR (Table 3; Supplementary Table 3). Based on these distributions, the CGT haplotypes can be divided into two groups: CGT1 (AFR unique haplotypes and HG01073.1) and CGT2 (non-AFR unique haplotypes and HG03667.1) (Table 3; Supplementary Fig. 3; Supplementary Table 3).

**Table 3.** Distribution of CGT haplotypes

| Lineage | Haplotype | AFR | EUR | SAS | EAS | AMR | Total |
| --- | --- | --- | --- | --- | --- | --- | --- |
| CGT1 | HG03105.0 | 1 |  |  |  |  | 1 |
|  | HG02095.1 | 1 |  |  |  |  | 1 |
|  | NA19908.0 | 4 |  |  |  |  | 4 |
|  | HG03572.1 | 1 |  |  |  |  | 1 |
|  | HG03311.0 | 8 |  |  |  |  | 8 |
|  | NA19116.1 | 1 |  |  |  |  | 1 |
|  | HG02433.0 | 8 |  |  |  |  | 8 |
|  | NA19468.0 | 1 |  |  |  |  | 1 |
|  | HG01073.1 | 12 |  |  |  | 1 | 13 |
|  | HG03100.1 | 3 |  |  |  |  | 3 |
|  | HG02095.0 | 1 |  |  |  |  | 1 |
|  | NA19472.1 | 1 |  |  |  |  | 1 |
|  | NA18909.0 | 1 |  |  |  |  | 1 |
|  | HG03437.1 | 1 |  |  |  |  | 1 |
| CGT2 | NA19664.0 |  |  |  |  | 2 | 2 |
|  | HG03667.1 | 2 | 1 | 5 | 8 |  | 16 |
|  | HG03772.1 |  |  | 1 |  |  | 1 |
|  | HG04042.1 |  |  | 4 |  | 3 | 7 |
|  | HG02494.1 |  |  | 1 |  | 1 | 2 |
|  | HG01455.0 |  |  |  |  | 1 | 1 |
| Total |  | 46 | 1 | 11 | 8 | 8 | 74 |

### High prevalence of the TCT-type in the ancestral population of non-AFR

The TCT-type is most prevalent in every non-AFR meta-population, and has an impact on the promoter composition of non-AFR. The positive selection on the CGC-type has influenced the promoter type composition in Asian and American populations [9]. However, the very low frequency of the CGC-type (0.2%) in EUR indicates that the impact of such positive selection on the promoter-type composition can be ignored in EUR. Thus, EUR is regarded as a meta-population in which promoter-type composition has not been influenced by local events. Since the TCT-type can be divided from the others by rs3759915, the TCT-type and the others can be treated as biallelic. To determine the uniqueness of the high prevalence of the TCT-type outside of Africa, we therefore examined the change of allele frequency of 308 schizophrenia-associated SNPs between AFR and EUR by dividing the SNPs into frequency bins.

Based on the TCT-type frequency in AFR (47.3%), 45 schizophrenia-associated SNPs showing 40.1%–49.2% frequency were selected. By comparison with the frequencies of these SNPs in EUR, the average value of fold difference was obtained as 1.20 (1.02–1.38; 99% confidence interval of the normal distribution accepted by Kolmogorov–Smirnov test). Since the fold difference of the TCT-type frequency from AFR is 2.04 in EUR (Fig. 1), the TCT-type frequency in EUR (96.3%) is much higher as expected from the frequency in AFR (48.3%–65.3%). Even though the TCT-type frequency is very high in EUR, we did not detect any signal of positive selection on the TCT-type using *F_st_*, extended haplotype homozygosity, or *F*_c_ ([9]; data not shown). Furthermore, the nucleotide diversity (*π*) of the TCT-type in EUR (0.07%) is similar to that in AFR (0.09%). These findings indicate that the very high frequency of the TCT-type in EUR does not result from the recent local selection, consistent with the high prevalence of the TCT-type in the ancestral population of non-AFR. The median value of fold difference from AFR in EUR is 0.97 using 41 schizophrenia-associated SNPs showing 90.1%–100.0% frequency in EUR. Using this median value, the TCT-type frequency in the ancestral population of non-AFR can be inferred to be around 90%.

**Fig. 1.**
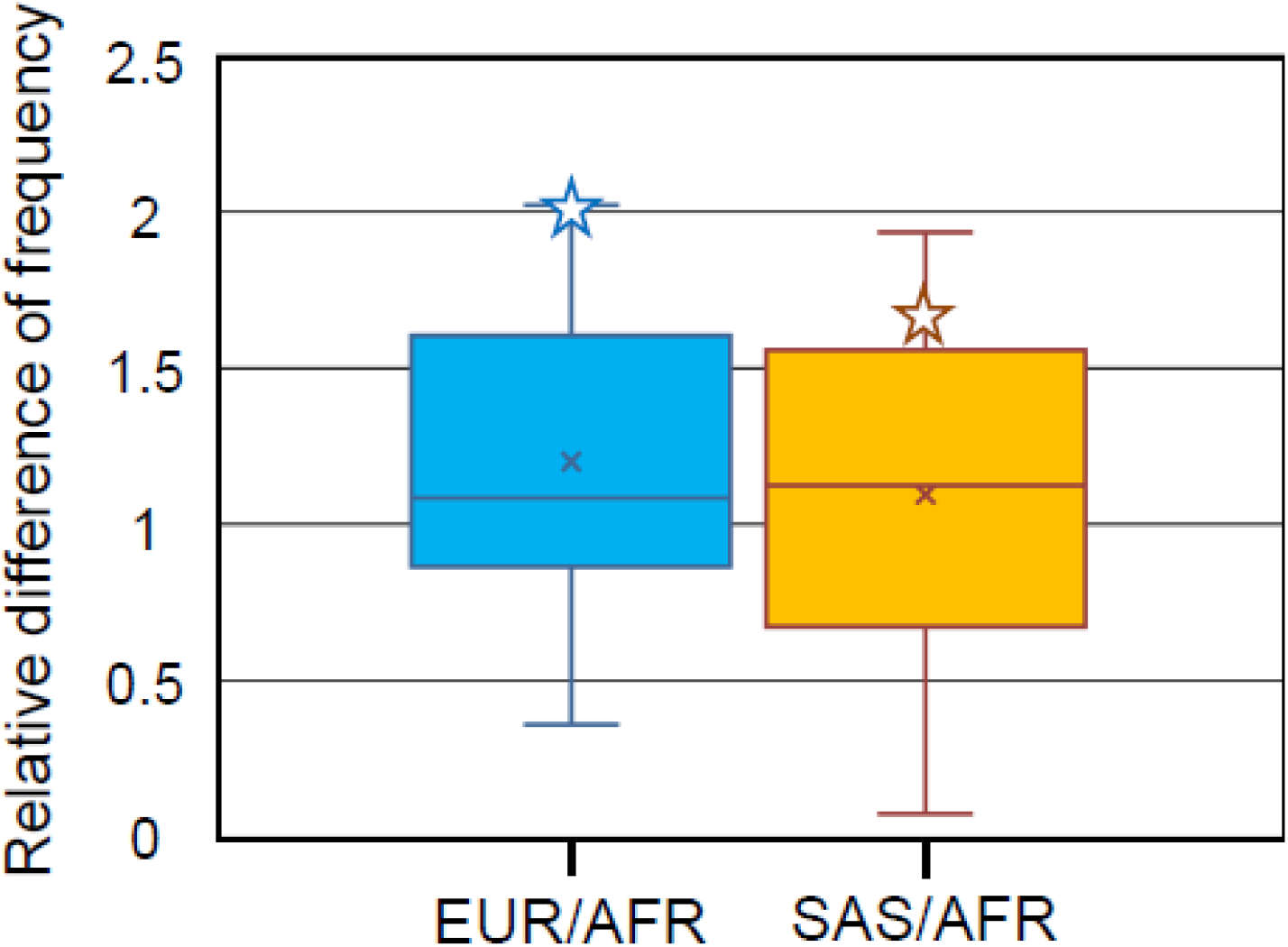
Change of frequencies in schizophrenia-associated genetic loci. Change of frequency is shown as difference of frequency in EUR and SAS relative to AFR. Based on the TCT-type frequency in AFR (47.3%), relative differences were calculated using the schizophrenia-associated genetic loci that show frequencies of 40.1%–49.2% in AFR. Relative difference of TCT-type is represented by an open star.

We also found similar results even if focus was placed on SAS. The average value of fold difference (SAS/AFR) is 1.10 (0.91–1.28; 99% confidence interval of the normal distribution accepted by Kolmogorov–Smirnov test) (Fig. 1). The TCT-type frequency in SAS (80.0%) is much higher than expected from the frequency in AFR (43.0%–60.5%) because the fold difference of the TCT-type frequency from AFR is 1.69 in SAS. The frequency of the TGT-type is higher in SAS than in EUR. According to the phylogenetic tree of the TGT-type haplotypes from SAS, this higher frequency results from the recent diversification of one TGT-type lineage (Supplementary Fig. 4). The timing of this diversification was obtained as 31 kyr [from ~11 kb of mean upstream homozygosity tract length (HTL)] using a published equation [9]. It is therefore concluded that the increase of the TGT-type frequency occurred after the migration out of Africa, compatible with the high prevalence of the TCT-type in the ancestral population of non-AFR.

### Unique TCT haplotype composition in LWK

In addition to the high prevalence of the TCT-type, the TCT haplotype composition in non-AFR is different from that in AFR (Table 2; Supplementary Table 2) as mentioned above. HG00097.1 is just one of the common haplotypes in AFR, but is markedly prevalent in non-AFR. In contrast, HG01556.0 is the most prevalent type in AFR, but very rare in non-AFR. In AFR, the frequencies of HG00097.1 in ACB (12.6%), ASW (23.4%), and LWK (18.3%) are higher than the average value of AFR (11.2%). Among these three subpopulations, ACB can be grouped with others (ESN, GWD, MSL, and YRI) because of the lack of significance of the difference in frequency between them (P > 0.01). As for ASW, admixture with Europeans occurred during its history [24], and the high frequency of HG00097.1 would result from the introgression by this admixture. By excluding ASW, AFR sub-populations can thus be divided into two groups: LWK and the others. LWK shows that the HG00097.1 frequency (18.3%) is not significantly different from that (24.0%) in SAS, the non-AFR meta-population showing the lowest TCT-type frequency (P = 0.258). In addition to this high frequency of HG00097.1, the frequency of HG00105.1, the second major haplotype in non-AFR, is highest in LWK (4.2%). Therefore, it is concluded that LWK is most closely related to non-AFR in AFR subpopulations.

To examine the effect of recent admixture on the close relationship between LWK and non-AFR, we focused on HG01556.0, the most prevalent haplotype in AFR. ASW shows the lowest frequency (7.8%) in AFR subpopulations. Since ASW has admixed with Europeans, this lowest frequency would result from the reduction of HG01556.0 by the admixture because of very low frequency of HG01556.0 in non-AFR (0.8% in AMR). In contrast to ASW, LWK shows the highest frequency (22.5%) in AFR subpopulations (Supplementary Table 2). This significant difference between ASW and LWK (P < 0.01) indicates that LWK has not undergone significant admixture with non-AFR.

In AFR, LWK shows the highest frequencies of both HG00097.1, the most prevalent type in non-AFR, and HG01556.0, the most prevalent type in AFR. In line with this, this hybrid composition in LWK reflects the TCT haplotype composition of the common ancestor of non-AFR and AFR. Interestingly, the TGT-type, the ancestral promoter type, has the highest frequency in LWK (58.1%; see Supplementary Table 1). Taking these findings together, it is likely that the promoter-type composition of LWK is close to those of AMH ancestors more than other subpopulations.

### Promoter types in AHs

We extracted genomic sequences from three Neanderthals (Vindija, Altai, and Chagyrskaya) and one Denisovan, and found that they possess only CGT at the three promoter SNPs (Supplementary Figs. 1 and 5). These AH samples came from Croatia (Vindija Neanderthal) and Siberia (Altai Neanderthal, Chagyrskaya Neanderthal, and Denisovan), and thus are from locations distant from each another [25, 26]. This shows that the CGT haplotype was widely dispersed in AH species. Only the CGT-type was identified from the AH samples, indicating that the CGT/CGT homozygote was common in AH population. Based on the identification of only CGT/CGT homozygosity in all four AH individuals, the CGT-type frequency in AH populations was estimated as over 71.7 % (95% credible interval from the β-distribution). To examine the existence of non-CGT-types in AH populations, we detected recombination tracts because their presence reflects the combination of promoter types. In this analysis, we used SNPs with MAF of 0.5% or more because some AH SNPs have an MAF in AMHs of less than 1%.

Among AH SNPs, eight SNP sites [rs145606077 (AH-SNP1), rs13379489 (AH-SNP2), rs4777968 (AH-SNP3), rs4570781 (AH-SNP4), rs144258052 (AH-SNP5), rs150882587 (AH-SNP6), rs192224395 (AH-SNP7), and rs373352748 (AH-SNP8)] are also polymorphic in AMHs (see Fig. 2 and Supplementary Fig. 5). The Altai haplotype of these SNPs, T-T-G-C-G-A-T-G, is shared with Denisovan-2, Vindija-2, and Chagyrskaya-2. Denisovan-1, Vindija-1, and Chagyrskaya-1 are different from Altai haplotype at five, three, and two AH-SNP sites, respectively. Chagyrskaya-1 and Vindija-1 have one unique allele, “C” at AH-SNP1 and “A” at AH-SNP3, respectively. In contrast, “A,” “C,” “C,” and “C” at AH-SNP5–8 are unique to Denisovan-1. To examine the recombination with non-CGT-type in the emergence of these unique alleles, we compared the lengths of tracts identical to Denisovan-1, Vindija-1, and Chagurskaya-1 among non-CGT haplotypes (Fig. 2; Supplementary Fig. 5). Uncertain sites (“N”) in AH sequences were ignored in this comparison.

**Fig. 2.**
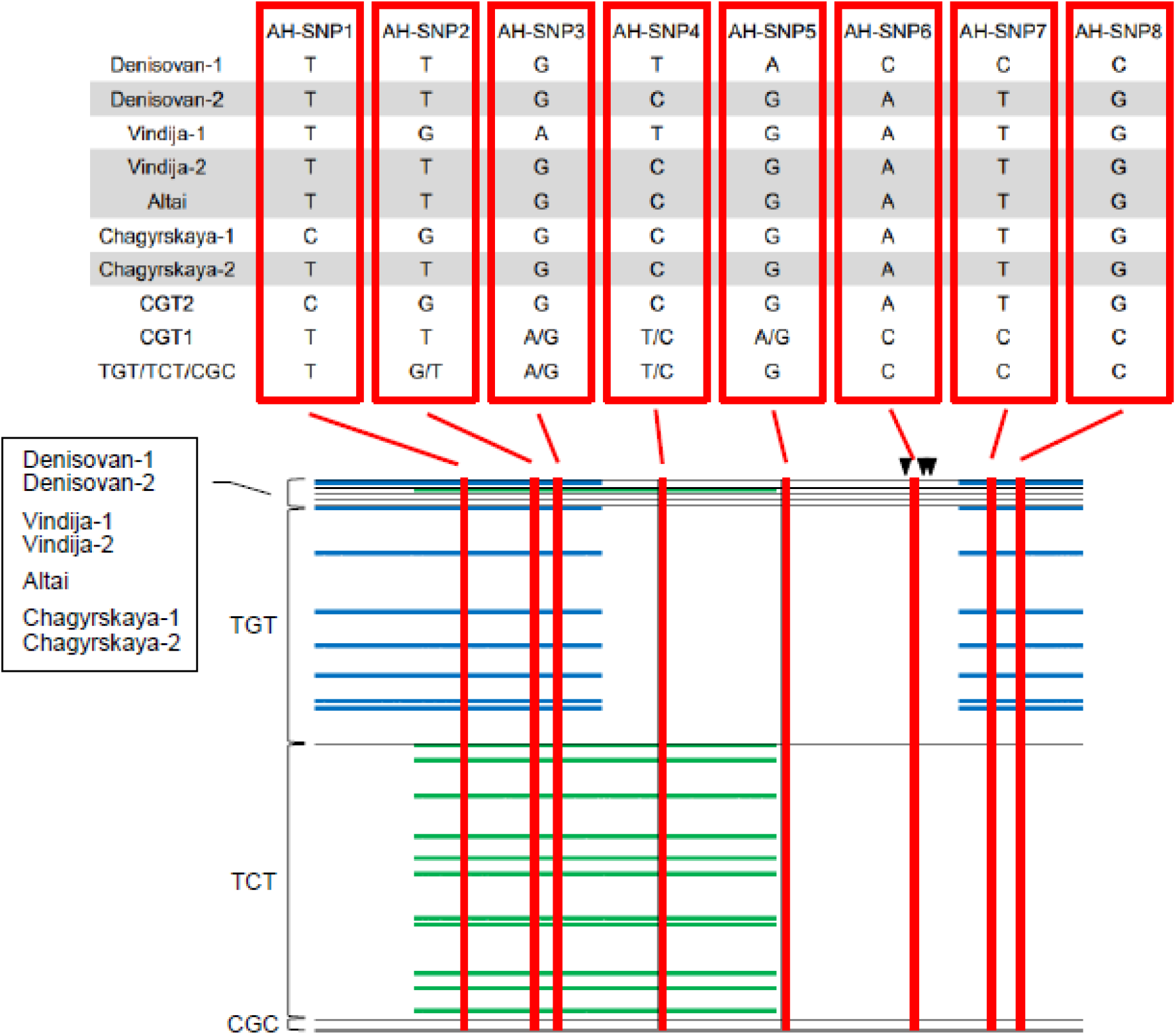
Comparison between AH haplotypes and AFR haplotypes. SNPs with minor allele frequency of 0.5% or over in the total AMH population were used, with the exception of AH-SNP5 (rs144258052; 0.34% in total). As for non-CGT-types, all AFR haplotypes were used. AH haplotypes of eight AH-SNP sites are represented in the upper panel. In the lower panel, the longest identical tracts detected between AH sequences and non-CGT haplotypes are represented by blue bars (Denisovan-1) and green bars (Vindija-1). Arrowheads represent positions of three promoter SNPs. Uncertain sites (“N”) in AH sequences were ignored in the comparison.

The tract identical to Denisovan-1 in the 3’ end region containing AH-SNP7 and AH-SNP8 was longest in some TGT haplotypes (Fig. 2; Supplementary Fig. 5), which suggests that “C”s in AH-SNP7 and AH-SNP8 were derived from a TGT-type by recombination. In addition, some of these TGT haplotypes show a tract identical to Denisovan-1 in the 5’ end region containing AH-SNP1–3. These findings indicate that the ~8-kb part containing the three promoter SNPs is sandwiched between the TGT-type sequences in the 18-kb region of Denisovan-1. It is therefore concluded that Densisovan-1 was established by recombination in which the ~8-kb middle part of the TGT-type was exchanged for CGT-type. Using a published equation [27] with the recombination rate of the 18-kb region (3.3 cM/Mb; [9]), the Neanderthal–Denisovan split time of 450 kyr [25], introgression time of 50 kyr, and generation time of 29 years, the length of the part containing the three promoter SNPs rules out that the Denisovan-1 sequence was derived from the AH common ancestor (P = 0.006). In other words, the recombination between the CGT- and TGT-types occurred in the AH population.

The tract identical to Vindija-1 in the region surrounding AH-SNP3 was longest in some TCT haplotypes (Fig. 2; Supplementary Fig. 5), which suggests that “A” at AH-SNP3 was derived from a TCT-type by recombination. The length of the identical tract is ~9 kb, which refutes that the Vindija-1 sequence is derived from the AH common ancestor (P = 0.002). In addition to the TGT-type, the TCT-type anticipated the recombination with the CGT-type in the AH population.

According to the position of the recombination tracts, Denisovan-1 has the TGT part at both ends of the 18-kb region (i.e., TGT-CGT-TGT), and Vindija-1 contains the TCT part in the middle (i.e., CGT-TCT-CGT). Since three CGT-types, two TGT-types (or recombinants with the TGT-type), and one TCT-type (or recombinant with the TCT-type) are needed to make these haplotype structures (Supplementary Fig. 6), the detected structural difference suggests that the frequency of the TGT-type was higher than that of the TCT-type in the AH population.

### Archaic introgression of the CGT-type from AHs into AMHs

To examine the phylogenetic relationship of the CGT haplotypes of AMHs and AHs, we constructed a phylogenetic tree. Interestingly, the obtained tree shows that all CGT2 haplotypes are clustered with AH sequences after deep divergence from the CGT1 haplotypes (Fig. 3). To explain the topology of this tree, two scenarios are proposed. One is ancestral polymorphism, meaning that the topology of the obtained phylogenetic tree already existed in the common ancestor of AMHs and AHs (incomplete lineage sorting). The other is archaic introgression after the split of AMHs and AHs [25, 28, 29]. The mean value of HTLs was 28,042 bp for the CGT2 sequences. Using a published equation [27] and conditions [30] with the exception of the recombination rate of the 18-kb region (3.3 cM/Mb; [9]), this mean HTL rules out the former scenario (P = 9.7 × 10^-14^). Unlike AH sequences, all of the CGT2 sequences form a single cluster in the phylogenetic tree after the divergence from AH sequences (Fig. 3), which indicates the archaic introgression into AMHs from AHs.

**Fig. 3.**
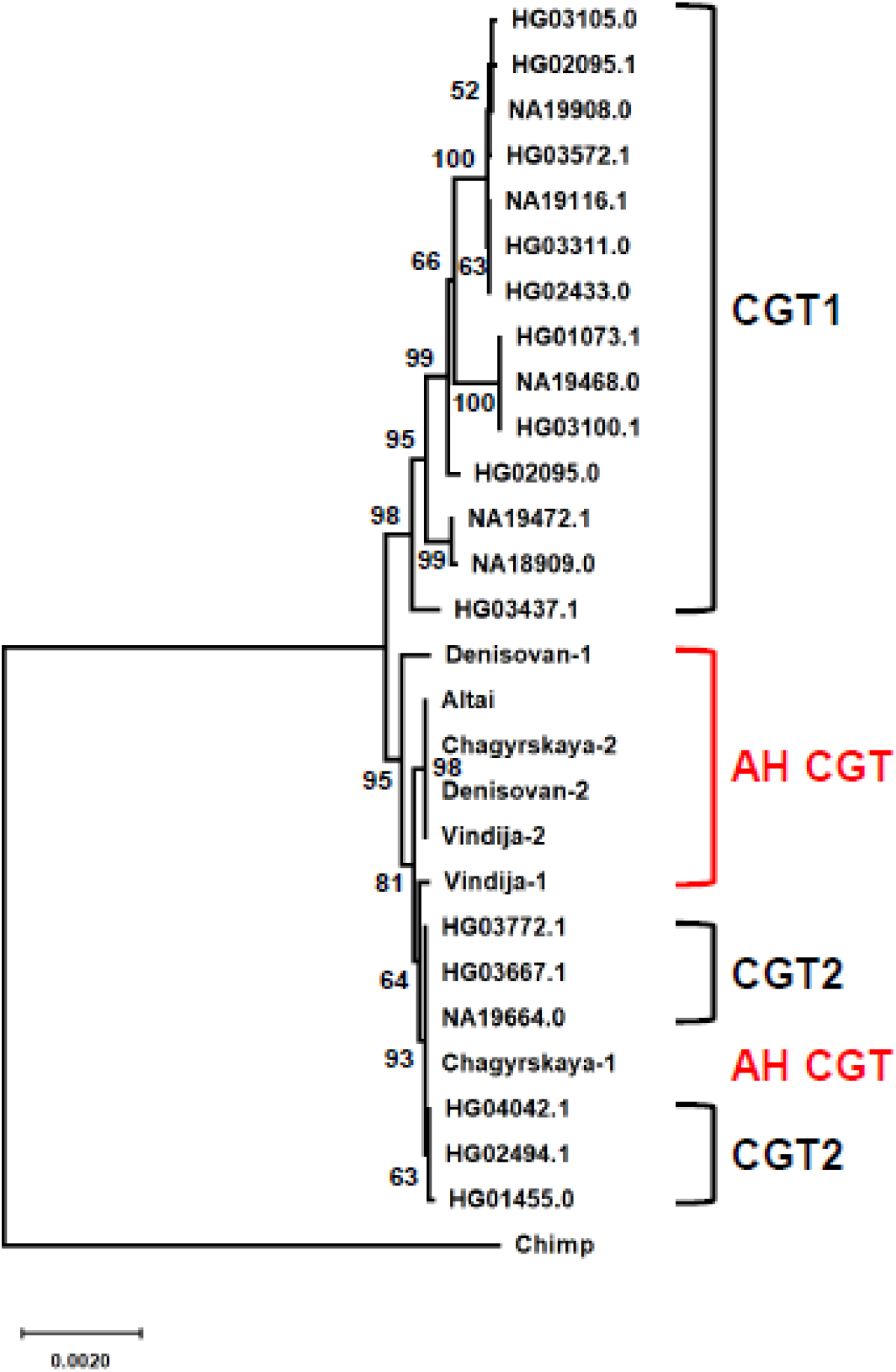
Phylogenetic relationship of the CGT-type sequences from AMHs and AHs. Phylogenetic tree is constructed using the Neighbor-Joining method with Jukes–Cantor correction. Bootstrap values of more than 50% from 1,000 replications are shown on the tree branches.

Based on the number of accumulated mutations within the CGT2 cluster, the maximum likelihood estimation [31] shows that the CGT2 lineage began to diversify into sub-lineages 46 kyr ago (33–59 kyr ago; mutation rate of 0.5 × 10^-9^ per site per year; [32]). Furthermore, derived alleles of six SNP sites (rs139630787, rs186699149, rs150882587, rs138393846, rs192224395, and rs373352748) are shared only between AH and CGT2 sequences (see Supplementary Figs. 1 and 5), and the ages of these variants were obtained as 71, 69, 66, 63, 63, and 82 kyr, respectively (joint clock; combined dataset; assuming 29 years per generation; [22]). These estimated times are much younger than the split time of AMHs and AHs (550–765 kyr) [25] and consistent with archaic introgression.

We detected 29 CGT2 sequences in *D*_1000_ (Table 3). Sixteen sequences are classified into the HG03667.1 haplotype found widely in AFR, EUR, EAS, and SAS (Table 3). The HG03667.1 haplotype is also an ancestral type because it is most closely related to the AH sequences (Fig. 3). The second major haplotype to which seven sequences belong is HG04042.1 found in SAS and AMR, and other haplotypes contain only one or two sequences from SAS and/or AMR (Table 3). SAS contains nearly half of the sequences (11 sequences) that belong to four haplotypes, including the ancestral type. It is therefore considered that the archaic introgression of the CGT-type occurred in an ancestral population of SAS.

### Archaic introgression from Neanderthals to Denisovans

We also examined the archaic introgression without determining the allelic linkage of AH sequences. To detect introgressed segments in the 978 SAS genomes, we applied the Sprime method [21]. Similar to the S* approach [33], it is reference-free and therefore advantageous when introgressed archaics are unknown. Chen et al. (2020) [34] pointed out false-negatives of these approaches where a modern reference (African) population is not completely free from archaic segments by introgression and/or gene flow. Among 116 possible introgressed segments identified in chromosome 15, one is about 250 kb long and spans the *ST8SIA2* gene. Curiously, however, the match proportion in this particular segment is ~83% both to the Altai Neanderthal genome [25] and to the Denisovan genome [28], suggesting introgression not only between modern humans and an archaic but also between archaics themselves. Examining the unphased genotype data of the Denisovan and Neanderthal genomes [26, 35], we found heterogeneous allele sharing and/or introgression in the 250 kb segment spanning the *ST8SIA2* gene. Focusing on one subsegment (22 kb long) ranging from 929133845 to 92936297 that includes the promoter SNPs, we computed the number of nucleotide differences (distance) in all pairwise comparisons of the four archaic diploid and one CGT2 (HG03667.1, Supplementary Table 3) haploid sequences. The Altai sequence turned out to be homozygous for the entire subsegment, so that the distance at a nucleotide site was defined as 1 and 1/2 where an Altai Neanderthal site differs from a homozygous and heterozygous site, respectively, of a diploid in comparison. Similarly, we computed the distance between two unphased diploid sequences. The distance matrix and the resulting model of the relatedness of the segments are given in Fig. 4. They suggest that one of the Denisovan segments was replaced by a Neanderthal one. Taken together with widespread admixture between Altai Neanderthals and Denisovans [36], multiple interspecies gene flows in the *ST8SIA2* gene make it difficult to know which AH provided archaic ancestry of CGT2 in the admixture with AMHs. It is noteworthy that the topology shown in Fig. 4 is compatible with that in Fig. 3. This suggests that our linkage determination in constructing the AH haplotype is reasonable.

**Fig. 4.**
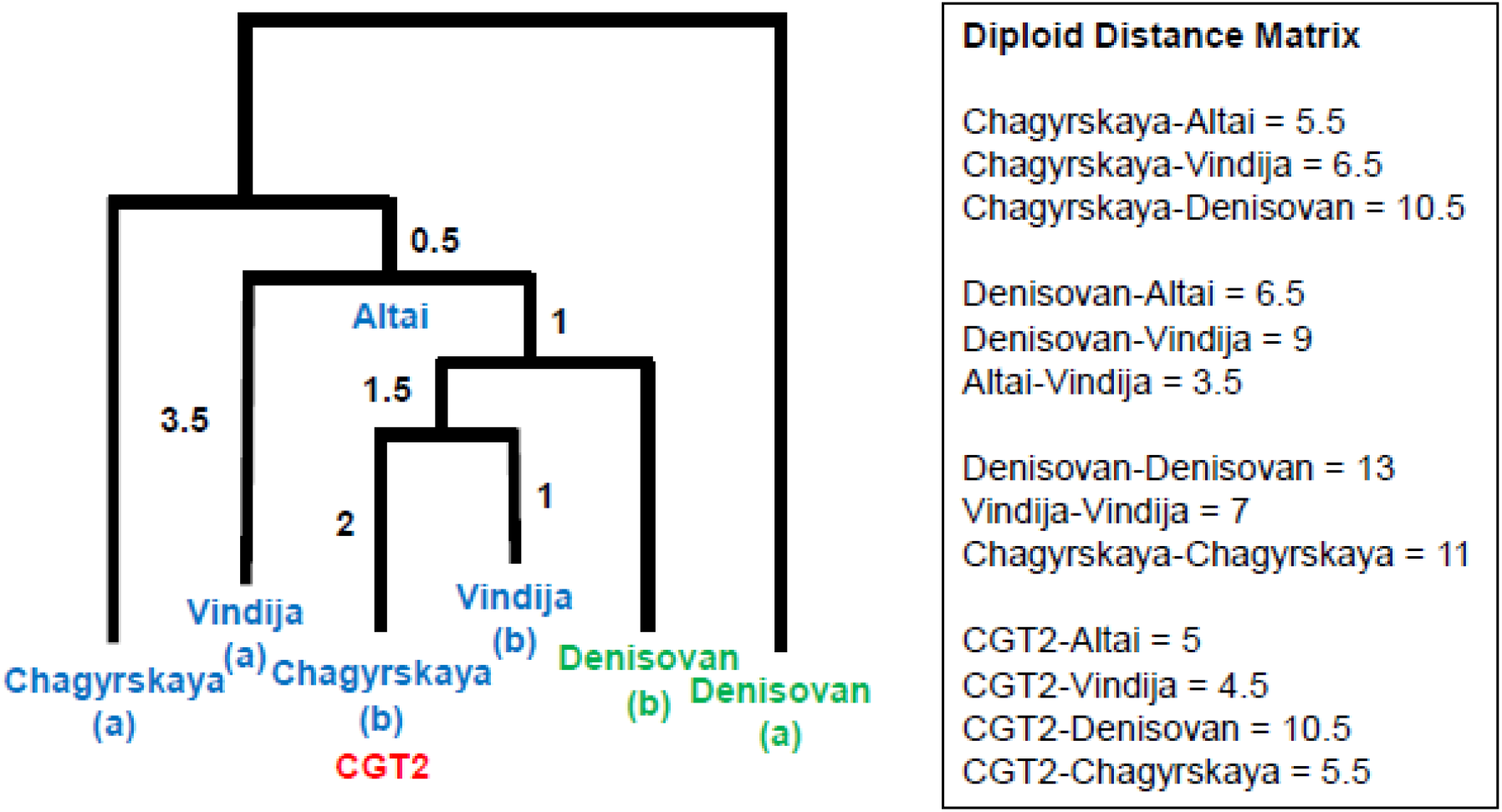
Diploid distance matrix and resulting tree of 22-kb region encompassing the three promoter SNPs. The tree topology was obtained by realizing compatibility of diploid distances among sequences.

### No detection of the archaic ancestry of non-CGT-type

We also examined the possibility of archaic introgression of non-CGT-types by focusing on SAS sequences. According to the phylogenetic tree of the CGT-type (Fig. 3), the CGT1 and CGT2 lineages correspond to the AMH and AH lineages. The numbers of different sites between the CGT1 and CGT2 haplotypes can be regarded as those expected in the AMH-AH divergence. Since parts of the HG03437.1, NA18909.0, and NA19472.1 sequences were influenced by recombination with the TGT- and TCT-types (Supplementary Fig. 5), we counted the number of different sites between the CGT1 and CGT2 haplotypes by eliminating these three haplotypes. The obtained numbers of different sites were 12–15 (Supplementary Table 4), which can be regarded as the AMH–AH site differences. This is consistent with the site differences (10–14 site differences) expected from the divergence time between AMHs and AHs (550–765 kyr; [25]).

Among the TGT haplotypes from SAS, only one haplotype, HG03779.1, represents 12–15 site differences from 15 haplotypes (Supplementary Fig. 7A; Supplementary Table 5). These 16 haplotypes were selected. However, a member of the 15 haplotypes, HG00132.1, is found widely in AFR (Supplementary Fig. 7A; Supplementary Table 1), and estimated ages of variants unique to HG03779.1 were much older than the time of the archaic introgression [35, 37, 38] (data not shown). Thus, it is concluded that there is no TGT haplotype derived from archaic ancestry.

Among the TCT haplotypes from SAS, 12–15 site differences from other haplotypes are found multiply in two haplotypes, HG02127.1 and HG03729.1 (Supplementary Table 6), and 12 haplotypes showed 12–15 site differences from both HG02127.1 and HG03729.1. Thus, we selected these 14 haplotypes. In the tree (Supplementary Fig. 7B), the 12 haplotypes form a single cluster and are distantly related to HG02127.1 and HG03729.1, which are deeply divergent from_one other. Seven of the 12 haplotypes are found in AFR (Supplementary Fig. 7B; Supplementary Table 2). In addition, HG02127.1 and HG03729.1 showed that the estimated ages of unique variants were much older than the time of archaic introgression [35, 37, 38] (data not shown). Thus, the selected haplotypes were not derived from the archaic ancestry.

The CGC haplotypes from SAS did not show any 12–15 site differences in the pairwise comparison (Supplementary Table 7). According to our previous work, the CGC-type emerged in the AMH lineage [9]. This is supported by the tree topology in which the CGC-type sequence belonging to the most prevalent haplotype is closely related to the CGT1 sequences because the archaic origin of the CGT2 sequences suggests that the CGT1 and CGT2 lineages correspond to the AMH and AH lineage, respectively (Figs. 3 and 5). The most prevalent CGC haplotype is widely distributed in AFR sub-populations, and Mbuti Pygmy, a deeply divergent population in the AMH lineage, has a CGC haplotype [9]. These findings support the AMH origin of the CGC-type. Taking these results together, it is concluded that the absence of haplotypes showing the AMH–AH site difference in the CGC haplotypes from SAS results from the AMH origin of the CGC-type.

**Fig. 5.**
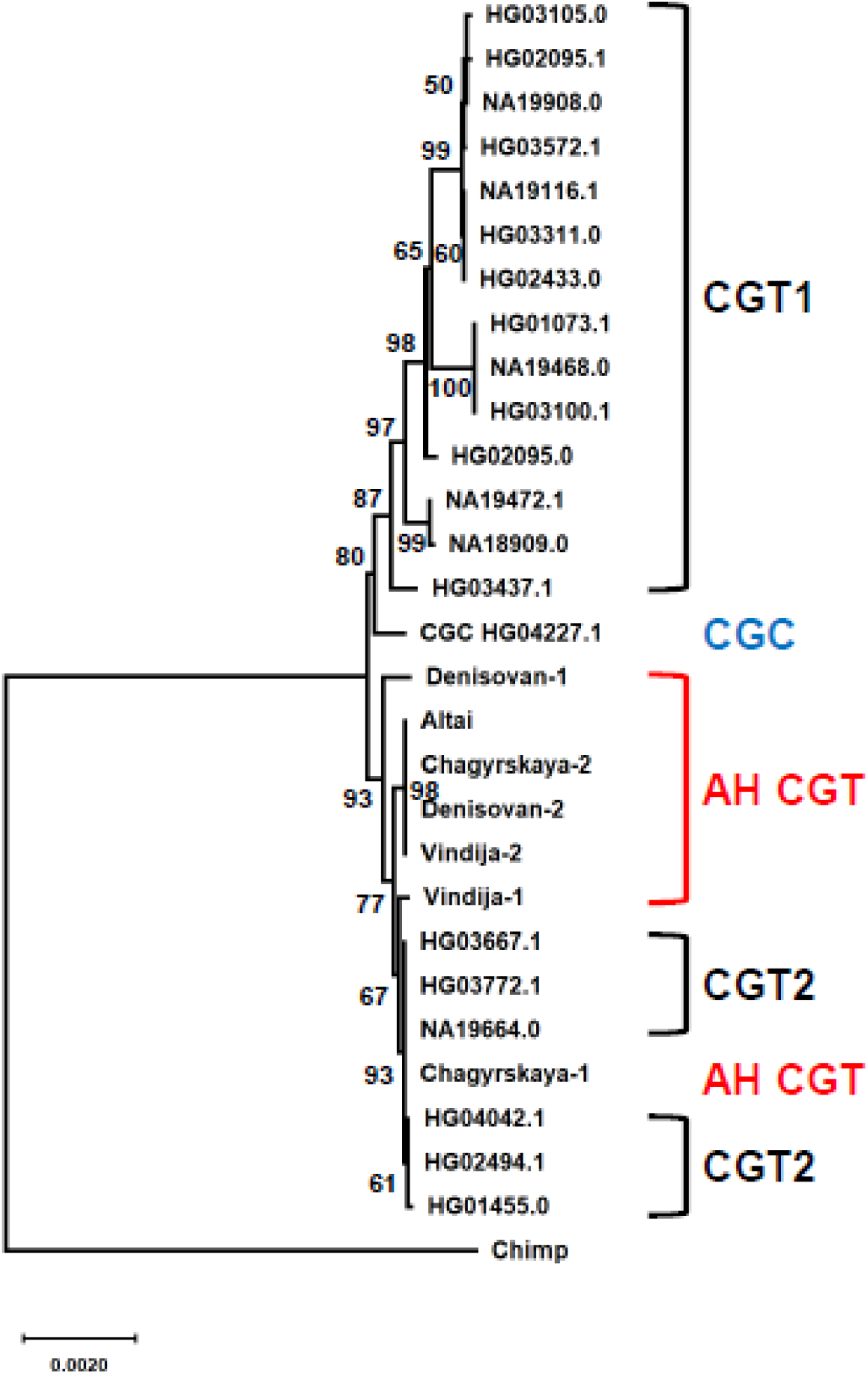
Tree of the CGT-type sequences with a CGC-type sequence belonging to the most prevalent CGC haplotype. The phylogenetic trees were constructed by the Neighbor-Joining method with Jukes–Cantor correction. Bootstrap values of more than 50% from 1,000 replications are shown on the tree branches.

Since the frequency of the CGT2 sequences is very low in AMHs (0.6% in total), the detection of only CGT-type introgression indicates that the impact of the archaic introgression on the promoter-type composition in AMHs is very small. Considering that the chance of introgression of promoter types would have been decided by their frequencies in AH populations, it is concluded that the frequency of the CGT-type was very high in AH populations, consistent with the inferred CGT-type frequency (>71.7%) in AH populations. The promoter-type composition is markedly different between AMHs and AHs (Supplementary Fig. 8).

### Distribution of promoter activities

We previously measured the promoter activity of each promoter type, and found that the CGC-type shows significantly lower activity than the others [9]. This comparison also indicates that promoter activity is significantly different among promoter types (P < 0.005; [23]), which indicates that each promoter type has unique promoter activity. Since the composition of promoter types is diversified among meta-populations, this raised the possibility that the promoter activity differs at the population level. To examine this possibility, we established a concept representing promoter activity at the population level (population promoter activity: PPA) as a population phenotype by using relative promoter activities measured experimentally [9]. As shown in Fig. 6A, EAS shows significantly lower PPA than the others (P < 0.01; Kolmogorov–Smirnov test). This is consistent with the highest frequency of the CGC-type in EAS ([9]; Supplementary Fig. 2D). Outside of Africa, a significant difference was also found among other meta-populations (P < 0.01; Fig. 6A). Furthermore, every non-AFR (EUR, SAS, EAS, and AMR) shows significantly lower PPA than AFR (P < 0.01; Fig. 6A). This is compatible with the highest frequency of the TCT-type showing the second lowest promoter activity in every non-AFR meta-population.

**Fig. 6.**
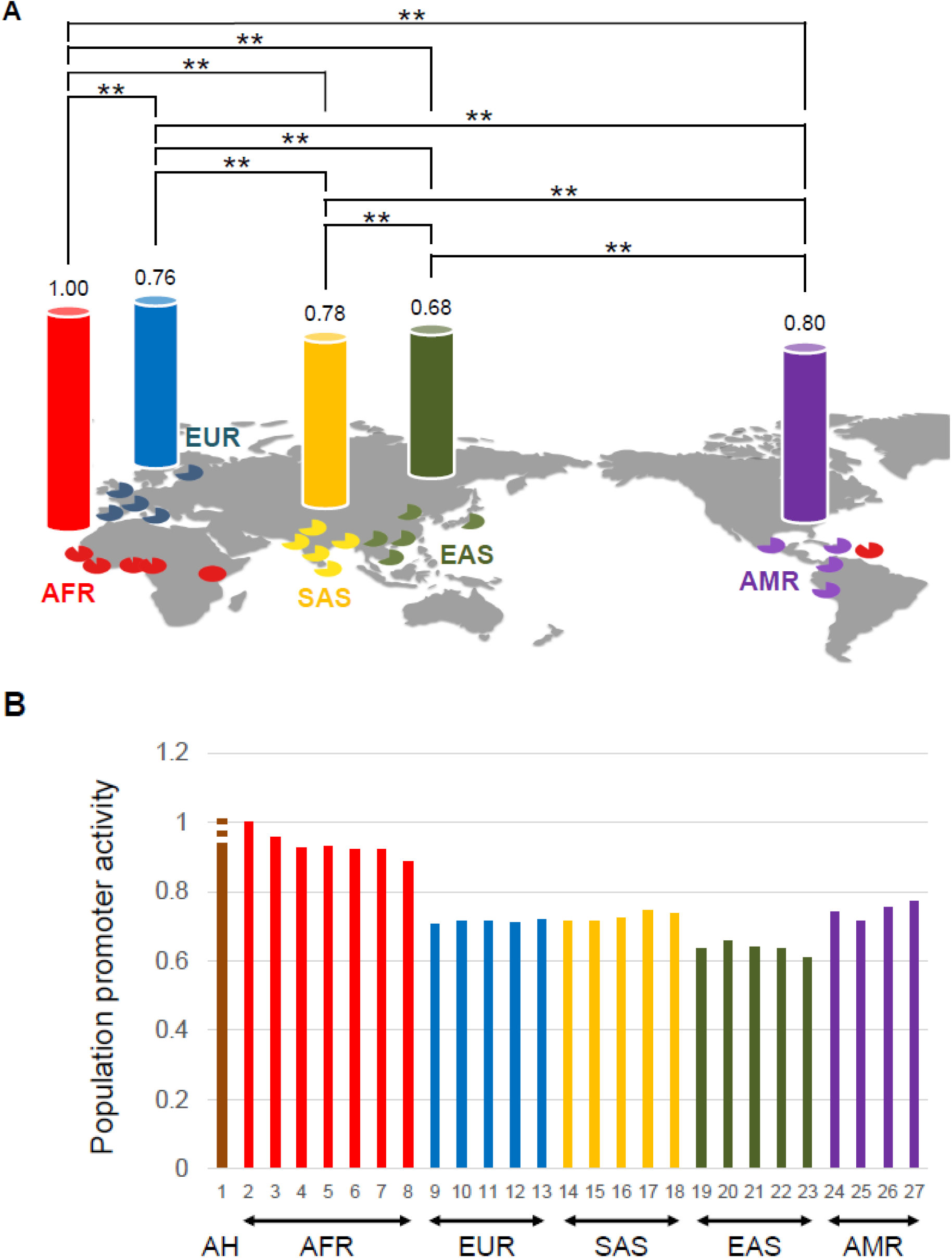
Distribution of PPAs. (A) PPA value of each meta-population is represented by relative activity as compared with AFR. Pie charts represent relative activities as compared with the highest activity (LWK) among subpopulations. **P<0.01. (B) Relative difference of PPA as compared with LWK. For AH, the inferred lowest PPA is represented by a filled line, and the top of the additional dashed line shows the highest inferred PPA value. 1, AH; 2, LWK; 3, ESN; 4, YRI; 5, ASW; 6, GWD; 7, MSL; 8, ACB; 9, TSI; 10, CEU; 11, GBR; 12, IBS; 13, FIN; 14, PJL; 15, GIH; 16, ITU; 17, BEB; 18, STU; 19, CDX; 20, KHV; 21, CHS; 22, CHB, 23, JPT; 24, MXL; 25, CLM; 26, PUR; 27, PEL.

LWK shows the highest PPA (Fig. 6B). This is compatible with the highest frequency (58.1%) of the TGT-type, which shows the highest promoter activity, in LWK. As mentioned previously, LWK has retained the ancestral promoter-type composition more than other subpopulations. Thus, the PPA has become lower even within Africa. Taking these findings together with the lower PPA in non-AFR than in AFR, we can see the trend toward lowered PPA in the spread of AMHs (Fig. 6B).

### PPA of AHs

As mentioned above, AH populations had the CGT-, TGT-, and TCT-types. Their CGT-type frequency was inferred to be over 71.7%, and the frequency of the TGT-type was assumed to be higher than that of the TCT-type. Based on these findings, the promoter-type combination of individuals was generated by random mating with no difference in the promoter-type frequency between males and females, and used for PPA calculation (Supplementary Table 8). All calculated PPAs were slightly higher than that of AFR (1.01- to 1.08-fold difference; Supplementary Table 8). Furthermore, even if we looked closely at the PPAs of AFR subpopulations, the inferred lowest PPA of AHs is higher than those of AFR subpopulations, with the exceptions of LWK and ESN, which show the top two PPAs (Fig. 6B; Supplementary Table 8).

Although AHs lived outside of Africa, their PPA was inferred to be higher than those of non-AFR subpopulations (see Fig. 6B). If the PPA of LWK is assumed to be close to that of the ancestor of AMHs and AHs, it is considered that the out-of-Africa migration did not significantly change the PPA at the origin of AHs. It is therefore likely that the trend toward lower PPA is unique to AMHs.

## Discussion

The numbers of heterozygous sites in Denisovan, Vindija, and Chagyrskaya sequences (12, 8, and 6 sites; Supplementary Fig. 1) were greater than those (~2 sites) detected in the whole-genome analysis [25]. As mentioned above, recombination with non-CGT-type was detected in Denisovan-1 and Vindija-1 (Fig. 2; Supplementary Fig. 5). This is consistent with the higher recombination rate (3.3 cM/Mb; [9]) in the 18-kb region as compared with the standard value [32]. In addition, the introgression from Neanderthals influenced Denisovan heterozygosity in the 18-kb region, consistent with the largest number of heterozygous sites in Denisovan sequences. A high recombination rate and introgression may be involved in the unexpectedly large number of heterozygous sites.

We did not detect any non-CGT haplotype derived from archaic ancestry. In cases where a promoter type that emerged uniquely in the AH lineage was introgressed, our method is useless because the AMH–AH site difference is used to identify candidates. In addition, if only sequences that underwent recombination with different promoter types in the AH lineage were introgressed, our method cannot detect haplotypes derived from such archaic ancestry as candidates because such haplotypes do not show the AMH–AH site difference with any haplotype. These are limitations of our method. As for the former, we did not encounter such a case in this study. However, even if it occurred, the distribution inside of Africa may be helpful to detect it because a wide distribution of haplotypes from the archaic ancestry cannot be expected in Africans. As for the latter, the use of unrecombined region would be important to avoid encountering such a case. However, the use of many haplotypes makes the length of an unrecombined region short, resulting in a low resolution. In this study, we used the 18-kb region containing no recombination hotspots. We therefore think that the resolution of our method was well maintained by the minimizing the effect of recombination in this study.

We found a difference in PPA among meta-populations (Fig. 6A). Since each promoter type has unique promoter activity [9, 23], this difference should depend on the composition of promoter types. The CGC-type has the lowest promoter activity, and its frequency is highest in EAS [9] (Table 1). It is therefore considered that the lowest PPA in EAS results from the increase in the frequency of the CGC-type under selection. This indicates that the selective increase of one functional genotype can influence a biological trend at the population level (population phenotype), even if the selected allele is still segregating at intermediate frequency.

The CGC-type frequency has been influenced by the recent positive selection in SAS, EAS, and AMR, but is very low in EUR and AFR. This suggests that the frequency of the CGC-type was very low in the ancestral population of non-AFR. However, the TCT-type shows the highest frequency in every non-AFR meta-population (Table 1). Furthermore, we found that the TCT-type was highly prevalent in the ancestral population of non-AFR (~90%). Since the TCT-type shows the lowest promoter activity with the exception of the CGC-type [9], it is reasonable that the lowered PPA was already established in the ancestral population of non-AFR, which is compatible with the lowered PPA shared by every non-AFR population.

Based on the TCT haplotype composition (Table 2; Supplementary Table 2), it is likely that the ancestral population of non-AFR (high frequency of HG00097.1 and low frequency of HG01556.0) emerged directly from the ancestor represented by LWK (high frequencies of both HG00097.1 and HG01556.0), accompanying the reduction of HG01556.0 frequency. While the TCT-type frequency is lowest in LWK (35.9%), it was inferred to be very high in the ancestral population of non-AFR (~90%). Therefore, the increase of the TCT-type would have occurred in the ancestral population of non-AFR, suggesting positive selection on the TCT-type in the non-AFR ancestor. The similar *π* values of the TCT-type between AFR and EUR indicate the concentration of the TCT-type in the non-AFR ancestor, and the out-of-Africa migration would have acted as a filter for this concentration.

Since schizophrenia develops through the interaction between genetic risk factors and environmental risk factors, an increase of the non-risk type of brain-related genes can be regarded as a result of adaptation to psychosocial stress [9, 11–15]. In terms of stress resistance, risk and non-risk types therefore confer sensitivity and tolerance to psychosocial stress and are regarded as sensitive and tolerant types, respectively. In addition to the CGC-type showing the lowest promoter activity, the TCT-type showing the second lowest promoter activity has been identified as a non-risk type in populations in which the frequency of the CGC-type is very low [8]. It is therefore considered that the lowering of PPA resulting from the increase of the frequency of tolerant types, namely, the TCT- and CGC-types, occurred due to selection as an adaptation to emerging stressful social environments. The lower PPA in non-AFR than in AFR (see Fig. 6) suggests that migration was a factor raising psychosocial stress, consistent with psychosocial stress in migration (immigration) having been identified as an environmental risk factor [13, 15, 39–41]. In the case of the spread of AMHs, it is considered that such stress was caused by the close encounters with earlier migrants including AHs. Therefore, it is concluded that the facilitation of social interaction (cooperation) by overcoming psychosocial stress has been advantageous in the spread of AMHs (Supplementary Fig. 9).

Societies and social networks act as collective brains in which individuals are regarded as neurons and drive cumulative cultural evolution [2, 3]. It is likely that overcoming psychosocial stress in close encounters with others makes collective brains bigger by increasing the degree of social interaction and the population size, and accelerates cumulative cultural evolution. Therefore, the trend toward lower PPA in the *ST8SIA2* gene would be an example of genetic tuning needed for the evolution of collective brains.

Since the PPA of AHs was inferred to be high (Fig. 6B), AHs might have been under stress when encountering AMHs. Because of the near monomorphism of the sensitive type (over 71.7% of the CGT-type) in AHs (Supplementary Fig. 8), it is assumed that AHs could not achieve a rapid change in their high PPA. Considering that AHs became extinct after the encounter with AMHs, their stable high PPA might have been disadvantageous for surviving the encounter with AMHs. Furthermore, the stable high PPA in AHs, resulting from near monomorphism of the sensitive type, also raises the possibility that AHs had stable small collective brains because they could not undergo increases of the degree of social interaction and the population size. This is consistent with the smaller population size in AHs than in AMHs [25] and supports Henrich’s hypothesis that AMHs have bigger collective brains than AHs [2].

Each meta-population represents a unique PPA (Fig. 6A). Diamond (1997) argued that the rate of spread of food production and technological innovations was higher in the Eurasian continent than in the African and American continents, resulting from their geographical and biogeographical differences (i.e., east–west major axis of the Eurasian continent, and north–south major axes of the African and American continents) [42]. Considering that the rate of spread depends on the degree of social interactions through migration, this is consistent with the finding that AFR and AMR show the two highest PPAs (Fig. 6A). The difference in AMR from other non-AFR meta-populations is smaller than that in AFR from AMR (Fig. 6A). However, this smaller difference might result from the short history of human settlement of the Americas (i.e., a recent reduction of psychosocial stress). Within the Eurasian continent, Diamond argued that geographical connectedness is high, moderate, and low in East Asia (China), Europe, and the Indian subcontinent, respectively [42]. Since the degree of social interactions through migration is proportional to that of geographical connectedness, this is compatible with the PPAs of EAS, EUR, and SAS (see Fig. 6A). Taking these findings together, the trend toward lower PPA might roughly reflect the local differences of the degree of social interaction resulting from the geographical and biogeographical differences.

Although the CGT-type is a sensitive type, archaic introgression of the CGT-type was accepted in the ancestral population of SAS. In addition, a local increase of the other sensitive type, the TGT-type, is found in SAS. The increase of these sensitive types in SAS ancestors might have been accepted because of low psychosocial stress resulting from low geographical connectedness in the Indian subcontinent [42].

The *ST8SIA2* gene was identified as a schizophrenia-related gene because it is functionally involved in sensitivity to psychosocial stress, an environmental risk factor of schizophrenia. In the adaptive increase of its tolerant types, the onset of schizophrenia is not necessarily required as a selective target (see Supplementary Fig. 10). Therefore, the trend toward lower PPA should not be hastily interpreted as a result of negative selection against the schizophrenia phenotype (i.e., positive and negative symptoms) or schizophrenia onset. Schizophrenia might be a modern phenotype induced by accidental exposure to genetically intolerable levels of environmental risk factors. Archaic admixture influences disease risk in AMHs [30, 43]. The introgression of the CGT-type from AHs is an example of an introgressed allele negatively affecting present-day humans in terms of the risk of schizophrenia.

One may argue that the trend toward lower PPA results in a lowering of the prevalence of schizophrenia from AFR to non-AFR. However, this is not guaranteed because of the variety of environmental risk factors. Although it is considered that ST8SIA2 played a central role in the adaptation to psychosocial stress associated with the spread of AMHs, its current role in the onset of schizophrenia should be limited because psychosocial stress caused by migration is only one of the environmental risk factors.

In conclusion, we detected a difference of the promoter-type composition between AMHs and AHs as well as within AMHs, and found that PPA has changed in the spread of AMHs by the compositional alteration of promoter types. In AMHs, the TCT-type was most prevalent in the ancestral population of non-AFR in addition to the selective increase of the CGC-type in Asia. In contrast, the frequency of the CGT-type was inferred to be very high in AHs. Promoter-type compositions were influenced independently in AMHs and AHs by interspecies gene flows: one into AMHs from AHs, and the other into Denisovans from Neanderthals. The difference of the promoter-type composition confers a unique PPA to each AMH population. PPA has become lower inside of Africa via the increase of the TCT-type. Furthermore, every non-AFR meta-population shows significantly lower PPA than AFR, which resulted from the high prevalence of the TCT-type in the ancestral population of non-AFR. In addition, EAS shows the lowest PPA by the selective increase of the CGC-type. These findings indicate that the increase of the TCT- and CGC-types, which show the two lowest promoter activities, has induced the trend toward lower PPA in the spread of AMHs. The inference of higher PPA in AHs than in AMHs suggests that this trend toward lower PPA is unique to AMHs. Since the TCT- and CGC-types can be regarded as tolerant types to psychosocial stress associated with the migration, it is concluded that the trend toward lower PPA results from the adaptation to the increased psychosocial stress associated with the spread of AMHs. The trend toward lower PPA is also considered as a form of genetic tuning for the evolution of collective brains in AMHs.

## Acknowledgements

This research was supported by the Japanese Society for Promotion of Science (JP16K07535 and JP19K06866 to T.H.). We thank Edanz (https://en-author-services.edanz.com/ac) for editing a draft of this manuscript.

