## Supplementary Figure 5 for "Lower promoter activity of the *ST8SIA2* gene has been favored in evolving human collective brains"

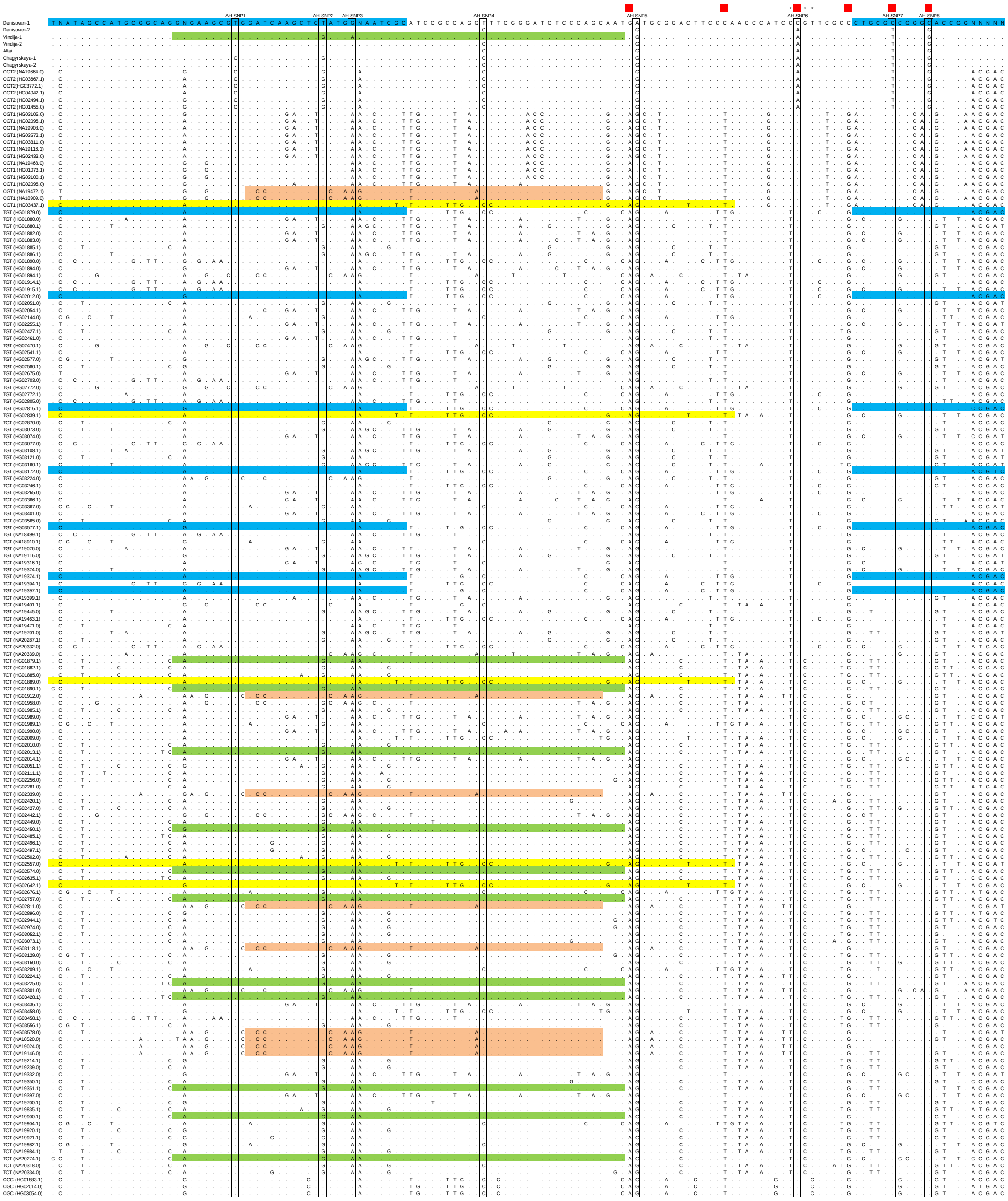


Supplementary Fig. 5. Sequence alignment of AH and AFR haplotypes. Except for rs144258052 (0.34% in total; AH-SNP3), SNPs with minor allele frequency of 0.5% or over in the total AMH population are shown. Open boxes represent AH-SNP sites. As for the non-CGT-types, all AFR haplotypes are represented. The three promoter SNPs are represented by asterisks. Longest identical tracts between AH sequences and non-CGT haplotypes are highlighted in blue (Denisovan-1) and green (Vindija-1), and those between CGT1 haplotypes and non-CGT haplotypes are highlighted in orange and yellow. The positions of six SNPs used for the estimation of variant age are indicated in red (see Supplementary Fig. 1). AH-SNP1 was not used for variant age estimation because the C allele is shared with a TCT haplotype found only in EUR (HG00145.0; two sequences; Supplementary Table 2). It is considered that HG00145.0 containing the C allele at AH-SNP1 is a recombinant with a CGT2 haplotype (HG03667.1) found only in EUR because of sequence identity in the region surrounding AH-SNP1 between them (data not shown).
