## Supplementary Table 1 for "Lower promoter activity of the *ST8SIA2* gene has been favored in evolving human collective brains"

Supplementary Table 1. Distribution of the TGT haplotypes*

| | AFR | | | | | | | EUR | | | | | SAS | | | | | EAS | | | | | AMR | | | | |
|---|---|---|---|---|---|---|---|---|---|---|---|---|---|---|---|---|---|---|---|---|---|---|---|---|---|---|---|
| TGT Haplotype | ACB | ASW | ESN | GWD | LWK | MSL | YRI | CEU | FIN | GBR | IBS | TSI | BEB | GIH | ITU | PJL | STU | CHB | CHS | CDX | JPT | KHV | CLM | MXL | PEL | PUR | Total |
| HG00132.1 | 1 | 5 | | 2 | | 2 | 2 | 7 | 9 | 4 | 3 | 3 | 17 | 14 | 18 | 12 | 17 | 11 | 18 | 15 | 20 | 15 | 10 | 14 | 36 | 9 | 264 |
| HG01271.0 | 12 | 4 | 13 | 11 | 17 | 7 | 12 | | | | 1 | 1 | | | | | | | | | | | 3 | | | | 81 |
| HG00728.0 | 8 | 4 | 9 | 13 | 12 | 8 | 8 | | | | 1 | | | | | | | | 1 | | | | 1 | | | 1 | 66 |
| HG00553.0 | 10 | 8 | 7 | 6 | 15 | 8 | 5 | | | | | | | | | | | | | | | | | 1 | | 3 | 63 |
| HG01886.1 | 6 | 7 | 6 | 4 | 5 | 11 | 16 | | | | | | | | | | | | | | | | | 1 | | | 56 |
| HG01241.0 | 1 | 3 | 7 | 9 | 7 | 10 | 11 | | | | | 1 | | | | | | | | | | | | 2 | | 1 | 52 |
| HG01110.1 | 7 | 2 | 5 | 7 | 7 | 12 | 5 | | | | | | | | | | | | | | | | | | | 2 | 47 |
| HG01082.1 | 6 | 3 | 4 | 6 | 4 | 3 | 9 | | | | | | | | | | | | | | | | | | | 2 | 37 |
| HG01171.1 | 3 | 1 | 2 | 11 | 11 | 2 | 4 | | | | | | | | | | | | | | | | | | | 1 | 35 |
| HG00409.1 | | | | | | | | | | | | | 2 | | | | | 2 | 3 | 3 | 6 | 8 | | 1 | 6 | 2 | 33 |
| HG01095.0 | 4 | 3 | 5 | 4 | 8 | | 4 | | | | | | | | | | | | | | | | 1 | | | 2 | 31 |
| HG01494.0 | 3 | 1 | 3 | 4 | 1 | 3 | 5 | | | | | | | | | | | | | | | | 1 | | | | 21 |
| HG00707.1 | | | | | | | | | | | | | | 1 | 1 | | | 1 | 1 | 1 | 3 | 1 | 2 | 3 | 5 | | 19 |
| HG00590.1 | | | | | | | | | | | | | | | | | 2 | 4 | 1 | 3 | 2 | 4 | 1 | | | | 17 |
| HG02051.0 | 4 | 2 | 1 | 4 | 1 | 1 | 1 | | | | | | | | | | | | | | | | | 1 | | | 15 |
| HG01250.1 | 2 | 1 | 2 | 1 | 4 | | 3 | | | | | | | | | | | | | | | | 1 | | | | 14 |
| HG00422.0 | | | | | | | | | | | | | | | | | | 1 | 2 | 4 | | 2 | 1 | | 2 | 1 | 13 |
| HG01894.0 | 4 | 1 | 3 | 1 | | | 3 | | | | | | | | | | | | | | | | | | | | 12 |
| HG01437.1 | 1 | | 2 | 3 | 2 | 1 | 1 | | | | | | | | | | | | | | | | 1 | | | | 11 |
| HG00125.0 | | | | | | | | | | 1 | | | 3 | 1 | 2 | 1 | 1 | | | | | | | | | | 9 |
| HG01894.1 | 3 | | | 1 | | 4 | | | | | | | | | | | | | | | | | | | | | 8 |
| HG01440.0 | | | 2 | 1 | 1 | 1 | | 1 | | | | | | | | | | | | | | | 1 | | | | 7 |
| HG00637.1 | | | 2 | | | 1 | 1 | | | | | | | | | | | | | | | | | | | 1 | 5 |
| HG01077.0 | | | | 2 | | 2 | | | | | | | | | | | | | | | | | | | | 1 | 5 |
| HG03074.0 | | 2 | 1 | | 1 | 1 | | | | | | | | | | | | | | | | | | | | | 5 |
| NA19026.0 | | | | | 5 | | | | | | | | | | | | | | | | | | | | | | 5 |
| NA19374.1 | | | | | 5 | | | | | | | | | | | | | | | | | | | | | | 5 |
| HG02577.0 | 1 | 1 | | | | 1 | 1 | | | | | | | | | | | | | | | | | | | | 4 |
| HG02772.0 | | | | 2 | | 2 | | | | | | | | | | | | | | | | | | | | | 4 |
| HG01890.0 | 1 | | 1 | | | | 1 | | | | | | | | | | | | | | | | | | | | 3 |
| HG02789.0 | | | | | | | | | | | | | 2 | | | 1 | | | | | | | | | | | 3 |
| HG03121.0 | | | 2 | | | 1 | | | | | | | | | | | | | | | | | | | | | 3 |
| NA19401.1 | | 1 | | | 2 | | | | | | | | | | | | | | | | | | | | | | 3 |
| HG00236.1 | | | | | | | | | | 1 | | | | | | | | | | | | | | | 1 | | 2 |
| HG01067.1 | | | | | | | | | | | | | | | | | | | | | | | | | | 2 | 2 |
| HG02054.1 | 1 | | | | | 1 | | | | | | | | | | | | | | | | | | | | | 2 |
| HG02675.0 | | | | 1 | | | 1 | | | | | | | | | | | | | | | | | | | | 2 |
| HG02772.1 | | | | 2 | | | | | | | | | | | | | | | | | | | | | | | 2 |
| HG03265.0 | | | 2 | | | | | | | | | | | | | | | | | | | | | | | | 2 |
| HG03615.0 | | | | | | | | | | | | | 1 | | | | 1 | | | | | | | | | | 2 |
| NA19399.1 | | | | | 2 | | | | | | | | | | | | | | | | | | | | | | 2 |
| NA20867.0 | | | | | | | | | | | | | | 2 | | | | | | | | | | | | | 2 |
| HG01092.1 | | | | | | | | | | | | | | | | | | | | | | | | | | 1 | 1 |
| HG01170.0 | | | | | | | | | | | | | | | | | | | | | | | | | | 1 | 1 |
| HG01871.0 | | | | | | | | | | | | | | | | | | | | | | 1 | | | | | 1 |
| HG02427.1 | 1 | | | | | | | | | | | | | | | | | | | | | | | | | | 1 |
| HG02494.0 | | | | | | | | | | | | | | | | 1 | | | | | | | | | | | 1 |
| HG02580.1 | 1 | | | | | | | | | | | | | | | | | | | | | | | | | | 1 |
| HG02660.0 | | | | | | | | | | | | | | | | 1 | | | | | | | | | | | 1 |
| HG02805.0 | | | | 1 | | | | | | | | | | | | | | | | | | | | | | | 1 |
| HG02816.1 | | | | 1 | | | | | | | | | | | | | | | | | | | | | | | 1 |
| HG02839.1 | | | | 1 | | | | | | | | | | | | | | | | | | | | | | | 1 |
| HG02870.0 | | | | 1 | | | | | | | | | | | | | | | | | | | | | | | 1 |
| HG03073.0 | | | | | | 1 | | | | | | | | | | | | | | | | | | | | | 1 |
| HG03160.1 | | | 1 | | | | | | | | | | | | | | | | | | | | | | | | 1 |
| HG03172.0 | | | 1 | | | | | | | | | | | | | | | | | | | | | | | | 1 |
| HG03224.0 | | | | | | 1 | | | | | | | | | | | | | | | | | | | | | 1 |
| HG03246.0 | | | | 1 | | | | | | | | | | | | | | | | | | | | | | | 1 |
| HG03366.1 | | | 1 | | | | | | | | | | | | | | | | | | | | | | | | 1 |
| HG03367.0 | | | 1 | | | | | | | | | | | | | | | | | | | | | | | | 1 |
| HG03401.0 | | | | | | 1 | | | | | | | | | | | | | | | | | | | | | 1 |
| HG03577.1 | | | | | | 1 | | | | | | | | | | | | | | | | | | | | | 1 |
| HG03765.1 | | | | | | | | | | | | | | | | 1 | | | | | | | | | | | 1 |
| HG03779.1 | | | | | | | | | | | | | | | 1 | | | | | | | | | | | | 1 |
| HG03782.1 | | | | | | | | | | | | | | | 1 | | | | | | | | | | | | 1 |
| HG03809.0 | | | | | | | | | | | | | 1 | | | | | | | | | | | | | | 1 |
| HG04003.1 | | | | | | | | | | | | | | | | | 1 | | | | | | | | | | 1 |
| HG04020.0 | | | | | | | | | | | | | | | 1 | | | | | | | | | | | | 1 |
| NA18499.1 | | | | | | | 1 | | | | | | | | | | | | | | | | | | | | 1 |
| NA18910.1 | | | | | | | 1 | | | | | | | | | | | | | | | | | | | | 1 |
| NA19116.0 | | | | | | | 1 | | | | | | | | | | | | | | | | | | | | 1 |
| NA19316.1 | | | | | 1 | | | | | | | | | | | | | | | | | | | | | | 1 |
| NA19324.0 | | | | | 1 | | | | | | | | | | | | | | | | | | | | | | 1 |
| NA19445.0 | | | | | 1 | | | | | | | | | | | | | | | | | | | | | | 1 |
| NA19463.1 | | | | | 1 | | | | | | | | | | | | | | | | | | | | | | 1 |
| NA19471.0 | | | | | 1 | | | | | | | | | | | | | | | | | | | | | | 1 |
| NA19701.0 | | 1 | | | | | | | | | | | | | | | | | | | | | | | | | 1 |
| NA20287.1 | | 1 | | | | | | | | | | | | | | | | | | | | | | | | | 1 |
| NA20339.0 | | 1 | | | | | | | | | | | | | | | | | | | | | | | | | 1 |
| NA20884.0 | | | | | | | | | | | | | | 1 | | | | | | | | | | | | | 1 |
| NA20901.1 | | | | | | | | | | | | | | 1 | | | | | | | | | | | | | 1 |
| Total | 80 | 52 | 83 | 100 | 115 | 86 | 96 | 8 | 9 | 6 | 5 | 5 | 26 | 20 | 24 | 17 | 22 | 19 | 26 | 26 | 31 | 31 | 23 | 23 | 50 | 30 | 1013 |
