## Supplementary Table 2 for "Lower promoter activity of the *ST8SIA2* gene has been favored in evolving human collective brains"

Supplementary Table 2. Distribution of the TCT haplotypes*

| | AFR | | | | | | | EUR | | | | | SAS | | | | | EAS | | | | | AMR | | | | |
|---|---|---|---|---|---|---|---|---|---|---|---|---|---|---|---|---|---|---|---|---|---|---|---|---|---|---|---|
| TCT Haplotype | ACB | ASW | ESN | GWD | LWK | MSL | YRI | CEU | FIN | GBR | IBS | TSI | BEB | GIH | ITU | PJL | STU | CHB | CHS | CDX | JPT | KHV | CLM | MXL | PEL | PUR | Total |
| HG00097.1 | 13 | 15 | 7 | 9 | 13 | 6 | 7 | 76 | 75 | 72 | 94 | 100 | 30 | 43 | 36 | 49 | 30 | 66 | 75 | 45 | 48 | 45 | 52 | 36 | 34 | 70 | 1146 |
| HG00105.1 | 1 | 1 | | 1 | 3 | | | 34 | 21 | 27 | 36 | 33 | 23 | 33 | 27 | 16 | 43 | 8 | 10 | 15 | 10 | 24 | 31 | 7 | 6 | 22 | 432 |
| HG00130.0 | 4 | 2 | | | | | | 11 | 5 | 8 | 6 | 6 | 21 | 16 | 10 | 12 | 17 | 10 | 16 | 15 | 3 | 10 | 1 | 3 | | 3 | 179 |
| HG00096.0 | 4 | 1 | 1 | 2 | 2 | 1 | 2 | 3 | 2 | 5 | 12 | 10 | 7 | 9 | 7 | 11 | 8 | 14 | 7 | 6 | 9 | 10 | 2 | 13 | 9 | 9 | 166 |
| HG01556.0 | 14 | 5 | 20 | 23 | 16 | 14 | 15 | | | | | | | | | | | | | | | | 1 | | 3 | | 111 |
| HG00131.0 | 2 | 2 | | 1 | 1 | | | 10 | 6 | 6 | 7 | 9 | 1 | 5 | 3 | 6 | 1 | | | | | | 11 | 1 | | 8 | 80 |
| HG00129.1 | 2 | 1 | | | | | | 6 | 14 | 2 | 4 | 1 | 4 | 5 | 5 | 9 | 10 | 3 | | | | 1 | 3 | 2 | 1 | 3 | 76 |
| HG01254.1 | 12 | 11 | 20 | 9 | 4 | 3 | 13 | | | | | | | | | | | | | | | | 1 | | | | 73 |
| HG01073.0 | 4 | 3 | 13 | 20 | 4 | 6 | 13 | | | | | | | | | | | | | | | | 1 | | 1 | 2 | 67 |
| HG00109.1 | | | | | | | | 8 | 28 | 7 | 7 | 6 | | 1 | | | 1 | | | | | | 2 | 3 | | 1 | 64 |
| HG01149.0 | 10 | 2 | 15 | 6 | 5 | 5 | 15 | | | | | | | | | | | | | | | | 3 | | | | 61 |
| HG01589.0 | | | | | | | | | | | | | 12 | 11 | 12 | 14 | 12 | | | | | | | | | | 61 |
| HG00732.0 | 3 | 3 | 10 | 14 | 3 | 12 | 7 | | | | | | | | | | | | | | | | 4 | 1 | 1 | 1 | 59 |
| HG00149.0 | 1 | 1 | | | | | | 6 | 9 | 3 | 3 | 3 | 7 | 4 | 5 | | 8 | | | | | 1 | 2 | 2 | | 1 | 56 |
| HG00100.0 | 1 | 1 | | | | | | 7 | 1 | 10 | 4 | 1 | 2 | 5 | 3 | 8 | 6 | | | | | | 2 | | | 4 | 55 |
| HG01047.1 | 12 | 2 | 3 | 9 | 3 | 5 | 11 | | | | | | | | | | | | | | | | | | | 4 | 49 |
| HG01188.0 | | | | | | | | | | | | | | | | | | 3 | | | 1 | | 5 | 8 | 23 | 1 | 41 |
| HG02127.1 | | | | | | | | | | | | | 5 | 8 | 15 | 4 | 6 | | | | | 2 | | | | | 40 |
| HG00117.1 | | | | | | | | 1 | 2 | 4 | 3 | 7 | | 1 | 5 | 1 | 1 | 2 | | 2 | | | 2 | 1 | | 3 | 35 |
| HG00114.0 | | | | | | | | | | 4 | 1 | 7 | 5 | 1 | 3 | 4 | 3 | 1 | | | | | | | 2 | | 31 |
| HG00100.1 | | | | | 1 | | | 3 | 1 | 3 | 2 | 4 | | | 1 | 1 | 1 | 3 | 2 | 2 | 1 | | 1 | | 1 | 1 | 28 |
| HG00118.0 | | | | | | 1 | | 2 | 2 | 2 | 1 | | | | | | | | | | | | 1 | 2 | | 14 | 25 |
| HG00096.1 | 1 | 1 | | | | | | 2 | 1 | 2 | 3 | 1 | 1 | | | 1 | 1 | | | 1 | 3 | | | 3 | 2 | 1 | 24 |
| HG03634.1 | | | | | | | | 1 | | | | 1 | 1 | 3 | 6 | 2 | 4 | | | | | | | 2 | | | 20 |
| HG01912.0 | 3 | | 9 | | 2 | 1 | 4 | | | | | | | | | | | | | | | | | | | | 19 |
| HG00099.1 | | | | | | | | 2 | 2 | 4 | 2 | 2 | | | | 4 | | | | | | | | | | 1 | 17 |
| HG00110.1 | 2 | 1 | | | | | | 2 | | 2 | 4 | 2 | | 1 | | | | 1 | 1 | | 1 | | | | | | 17 |
| HG02485.1 | 2 | 2 | 2 | 5 | 2 | 1 | 1 | | | | | | | | | | | | | | | | | | | | 15 |
| HG00140.1 | | | | | | | | 3 | 3 | 1 | 2 | 2 | | | 1 | | | | | | | | | | | 2 | 14 |
| HG00099.0 | | | | | | | | | | 2 | | | 2 | 4 | 1 | 3 | | | | | | | | | | | 12 |
| HG00123.1 | | | | | | | | | | 2 | | 1 | | 1 | 1 | 1 | 4 | | | | 1 | | | | | 1 | 12 |
| HG00102.0 | | | | | | | | 1 | | 2 | 3 | | | | | | | | | 1 | | 1 | | 2 | | 1 | 11 |
| HG00536.1 | | | | | | | | | | | | | | | | | | 6 | 2 | 1 | 1 | | | | | | 10 |
| HG01311.1 | 1 | | 1 | | 2 | 1 | 3 | | | | | | | | | | | | | | | | | | | 1 | 9 |
| HG00339.0 | | | | | | | | 1 | 2 | | | 1 | | | | | | | | | | | 1 | 1 | | 2 | 8 |
| HG02642.1 | | | | 3 | | 4 | 1 | | | | | | | | | | | | | | | | | | | | 8 |
| HG03686.1 | | | | | | | | | | | | | | 2 | 4 | | 2 | | | | | | | | | | 8 |
| HG01354.1 | | | | | | | | 3 | | | 1 | 2 | | | | | | | | | | | 1 | | | | 7 |
| HG02058.1 | | | | | 1 | | | 1 | | | | | | | | | | 1 | | | 1 | 3 | | | | | 7 |
| HG03836.0 | | | | | | | | | | | | | 1 | 1 | 4 | | 1 | | | | | | | | | | 7 |
| HG01991.1 | | | | | | | | 1 | | | | 1 | | 1 | | | 1 | 1 | | | | | | | 1 | | 6 |
| HG02687.0 | | | | | | | | | | | | | | | 1 | 1 | 4 | | | | | | | | | | 6 |
| HG00182.0 | | | | | | | | | 3 | | | | | | | 1 | 1 | | | | | | | | | | 5 |
| HG02051.1 | 4 | | | 1 | | | | | | | | | | | | | | | | | | | | | | | 5 |
| HG02442.1 | 2 | | | | 1 | | 2 | | | | | | | | | | | | | | | | | | | | 5 |
| HG00334.1 | 1 | | | | | | | | 2 | | | | | | | | | | | | | | 1 | | | | 4 |
| HG01284.0 | 1 | | 1 | | 1 | | | | | | | | | | | | | | | | | | 1 | | | | 4 |
| HG01389.1 | | | | | | | | | | | 2 | | | | | | | | | | | | 1 | | 1 | | 4 |
| HG02734.1 | | | | | | | | | | | | | 1 | 2 | | 1 | | | | | | | | | | | 4 |
| HG00106.0 | | 1 | | | | | | | | 1 | | 1 | | | | | | | | | | | | | | | 3 |
| HG00171.1 | | | | | | | | | 1 | 1 | | | | | | | | | | | | | 1 | | | | 3 |
| HG00259.0 | | | | | | | | | 1 | 1 | 1 | | | | | | | | | | | | | | | | 3 |
| HG00266.1 | | | | | | | | | 1 | | | | | | | | | | | | | | | | | 2 | 3 |
| HG00640.0 | | | | | 2 | | | | | | | | | | | | | | | | | | | | | 1 | 3 |
| HG01121.0 | | | | | | | | | | | | | | | | | 1 | | | | 1 | | 1 | | | | 3 |
| HG01357.1 | | | | | | | | | | | 1 | | | | 1 | | | | | | | | 1 | | | | 3 |
| HG01438.0 | | | | | 2 | | | | | | | | | | | | | | | | | | 1 | | | | 3 |
| HG01485.0 | | | | | | | | 1 | | | | | | | | | | | | | 1 | | 1 | | | | 3 |
| HG01871.1 | | | | | | | | | | | | | | | | | | | | | | 3 | | | | | 3 |
| HG02130.1 | | | | | | | | | | | | | | | | | | 1 | | | | 2 | | | | | 3 |
| HG02600.0 | | | | | | | | | | | | | | | | 1 | 2 | | | | | | | | | | 3 |
| HG02793.1 | | | | | | | | | | | | 1 | 1 | | | 1 | | | | | | | | | | | 3 |
| HG03129.0 | | | 2 | | | | 1 | | | | | | | | | | | | | | | | | | | | 3 |
| HG03301.0 | | | 1 | | 2 | | | | | | | | | | | | | | | | | | | | | | 3 |
| NA18945.1 | | | | | | | | | | | | | | | | | | | | | 3 | | | | | | 3 |
| HG00133.1 | | 1 | | | | | | | | 1 | | | | | | | | | | | | | | | | | 2 |
| HG00145.0 | | | | | | | | | 1 | 1 | | | | | | | | | | | | | | | | | 2 |
| HG00306.1 | | | | | | | | 1 | 1 | | | | | | | | | | | | | | | | | | 2 |
| HG00553.1 | | | | | | | | | | | | | | | | | | | | | | | | | | 2 | 2 |
| HG01518.0 | | | | | | | | | | | 1 | | | | | | | | | | | | | | 1 | | 2 |
| HG01631.1 | | | | | | | | | | | 2 | | | | | | | | | | | | | | | | 2 |
| HG02339.0 | 1 | | | 1 | | | | | | | | | | | | | | | | | | | | | | | 2 |
| HG02574.0 | | 1 | | 1 | | | | | | | | | | | | | | | | | | | | | | | 2 |
| HG02654.0 | | | | | | | | | | | | | 1 | | | 1 | | | | | | | | | | | 2 |
| HG02896.0 | | | | 1 | | | 1 | | | | | | | | | | | | | | | | | | | | 2 |
| HG02974.0 | | | 2 | | | | | | | | | | | | | | | | | | | | | | | | 2 |
| HG03015.1 | | | | | | | | | | | | | | | | 1 | | 1 | | | | | | | | | 2 |
| HG03052.1 | | | | | | 2 | | | | | | | | | | | | | | | | | | | | | 2 |
| HG03428.1 | | | | | | 1 | | | | | | | | | | 1 | | | | | | | | | | | 2 |
| HG03458.1 | | | | | | 2 | | | | | | | | | | | | | | | | | | | | | 2 |
| HG03717.1 | | | | | | | | | | | | | | | 2 | | | | | | | | | | | | 2 |
| HG03730.0 | | | | | | | | | | | | | | | 1 | | 1 | | | | | | | | | | 2 |
| HG03742.1 | | | | | | | | | | | | | 1 | | 1 | | | | | | | | | | | | 2 |
| NA18520.0 | | | | | | | 2 | | | | | | | | | | | | | | | | | | | | 2 |
| NA20503.1 | | | | | | | | | | | | 1 | | 1 | | | | | | | | | | | | | 2 |
| HG00154.1 | | | | | | | | | | 1 | | | | | | | | | | | | | | | | | 1 |
| HG00171.0 | | | | | | | | | 1 | | | | | | | | | | | | | | | | | | 1 |
| HG00178.1 | | | | | | | | | 1 | | | | | | | | | | | | | | | | | | 1 |
| HG00238.0 | | | | | | | | | | 1 | | | | | | | | | | | | | | | | | 1 |
| HG00251.1 | | | | | | | | | | 1 | | | | | | | | | | | | | | | | | 1 |
| HG00284.1 | | | | | | | | | 1 | | | | | | | | | | | | | | | | | | 1 |
| HG00327.1 | | | | | | | | | 1 | | | | | | | | | | | | | | | | | | 1 |
| HG00457.0 | | | | | | | | | | | | | | | | | | | 1 | | | | | | | | 1 |
| HG00743.1 | | | | | | | | | | | | | | | | | | | | | | | | | | 1 | 1 |
| HG01077.1 | | | | | | | | | | | | | | | | | | | | | | | | | | 1 | 1 |
| HG01108.0 | | | | | | | | | | | | | | | | | | | | | | | | | | 1 | 1 |
| HG01190.0 | | | | | | | | | | | | | | | | | | | | | | | | | | 1 | 1 |
| HG01251.1 | | | | | | | | | | | | | | | | | | | | | | | 1 | | | | 1 |
| HG01413.0 | | | | | | | | | | | | | | | | | | | | | | | | | | 1 | 1 |
| HG01519.1 | | | | | | | | | | | 1 | | | | | | | | | | | | | | | | 1 |
| HG01628.0 | | | | | | | | | | | 1 | | | | | | | | | | | | | | | | 1 |
| HG01631.0 | | | | | | | | | | | 1 | | | | | | | | | | | | | | | | 1 |
| HG01866.1 | | | | | | | | | | | | | | | | | | | | | | 1 | | | | | 1 |
| HG01977.0 | | | | | | | | | | | | | | | | | | | | | | | | | 1 | | 1 |
| HG02178.1 | | | | | | | | | | | | | | | | | | | | 1 | | | | | | | 1 |
| HG02232.1 | | | | | | | | | | | 1 | | | | | | | | | | | | | | | | 1 |
| HG02236.0 | | | | | | | | | | | 1 | | | | | | | | | | | | | | | | 1 |
| HG02277.1 | | | | | | | | | | | | | | | | | | | | | | | | | 1 | | 1 |
| HG02427.0 | 1 | | | | | | | | | | | | | | | | | | | | | | | | | | 1 |
| HG02502.0 | 1 | | | | | | | | | | | | | | | | | | | | | | | | | | 1 |
| HG02635.1 | | | | 1 | | | | | | | | | | | | | | | | | | | | | | | 1 |
| HG02757.0 | | | | 1 | | | | | | | | | | | | | | | | | | | | | | | 1 |
| HG02811.0 | | | | 1 | | | | | | | | | | | | | | | | | | | | | | | 1 |
| HG02944.1 | | | 1 | | | | | | | | | | | | | | | | | | | | | | | | 1 |
| HG03006.1 | | | | | | | | | | | | | 1 | | | | | | | | | | | | | | 1 |
| HG03018.1 | | | | | | | | | | | | | | | | 1 | | | | | | | | | | | 1 |
| HG03160.0 | | | 1 | | | | | | | | | | | | | | | | | | | | | | | | 1 |
| HG03209.1 | | | | | | 1 | | | | | | | | | | | | | | | | | | | | | 1 |
| HG03224.1 | | | | | | 1 | | | | | | | | | | | | | | | | | | | | | 1 |
| HG03228.1 | | | | | | | | | | | | | | | | 1 | | | | | | | | | | | 1 |
| HG03229.1 | | | | | | | | | | | | | | | | 1 | | | | | | | | | | | 1 |
| HG03436.1 | | | | | | 1 | | | | | | | | | | | | | | | | | | | | | 1 |
| HG03556.1 | | | | | | 1 | | | | | | | | | | | | | | | | | | | | | 1 |
| HG03578.0 | | | | | | 1 | | | | | | | | | | | | | | | | | | | | | 1 |
| HG03681.0 | | | | | | | | | | | | | | | | | 1 | | | | | | | | | | 1 |
| HG03697.1 | | | | | | | | | | | | | | | | | 1 | | | | | | | | | | 1 |
| HG03706.0 | | | | | | | | | | | | | | | | 1 | | | | | | | | | | | 1 |
| HG03713.0 | | | | | | | | | | | | | | | 1 | | | | | | | | | | | | 1 |
| HG03722.1 | | | | | | | | | | | | | | | 1 | | | | | | | | | | | | 1 |
| HG03729.1 | | | | | | | | | | | | | | | 1 | | | | | | | | | | | | 1 |
| HG03752.1 | | | | | | | | | | | | | | | | | 1 | | | | | | | | | | 1 |
| HG03765.0 | | | | | | | | | | | | | | | | 1 | | | | | | | | | | | 1 |
| HG03920.0 | | | | | | | | | | | | | 1 | | | | | | | | | | | | | | 1 |
| HG03965.0 | | | | | | | | | | | | | | | 1 | | | | | | | | | | | | 1 |
| HG04001.0 | | | | | | | | | | | | | | | 1 | | | | | | | | | | | | 1 |
| HG04134.0 | | | | | | | | | | | | | 1 | | | | | | | | | | | | | | 1 |
| HG04155.1 | | | | | | | | | | | | | 1 | | | | | | | | | | | | | | 1 |
| HG04194.1 | | | | | | | | | | | | | 1 | | | | | | | | | | | | | | 1 |
| NA07048.0 | | | | | | | | 1 | | | | | | | | | | | | | | | | | | | 1 |
| NA07051.0 | | | | | | | | 1 | | | | | | | | | | | | | | | | | | | 1 |
| NA11832.0 | | | | | | | | 1 | | | | | | | | | | | | | | | | | | | 1 |
| NA11992.1 | | | | | | | | 1 | | | | | | | | | | | | | | | | | | | 1 |
| NA18549.1 | | | | | | | | | | | | | | | | | | 1 | | | | | | | | | 1 |
| NA18605.1 | | | | | | | | | | | | | | | | | | 1 | | | | | | | | | 1 |
| NA18631.1 | | | | | | | | | | | | | | | | | | 1 | | | | | | | | | 1 |
| NA18757.0 | | | | | | | | | | | | | | | | | | 1 | | | | | | | | | 1 |
| NA19024.0 | | | | | 1 | | | | | | | | | | | | | | | | | | | | | | 1 |
| NA19146.0 | | | | | | | 1 | | | | | | | | | | | | | | | | | | | | 1 |
| NA19700.1 | | 1 | | | | | | | | | | | | | | | | | | | | | | | | | 1 |
| NA19771.1 | | | | | | | | | | | | | | | | | | | | | | | | 1 | | | 1 |
| NA19904.1 | | 1 | | | | | | | | | | | | | | | | | | | | | | | | | 1 |
| NA19920.1 | | 1 | | | | | | | | | | | | | | | | | | | | | | | | | 1 |
| NA19982.1 | | 1 | | | | | | | | | | | | | | | | | | | | | | | | | 1 |
| NA20274.1 | | 1 | | | | | | | | | | | | | | | | | | | | | | | | | 1 |
| NA20318.0 | | 1 | | | | | | | | | | | | | | | | | | | | | | | | | 1 |
| NA20334.0 | | 1 | | | | | | | | | | | | | | | | | | | | | | | | | 1 |
| NA20503.0 | | | | | | | | | | | | 1 | | | | | | | | | | | | | | | 1 |
| NA20542.0 | | | | | | | | | | | | 1 | | | | | | | | | | | | | | | 1 |
| NA20783.1 | | | | | | | | | | | | 1 | | | | | | | | | | | | | | | 1 |
| NA20786.0 | | | | | | | | | | | | 1 | | | | | | | | | | | | | | | 1 |
| NA20798.1 | | | | | | | | | | | | 1 | | | | | | | | | | | | | | | 1 |
| NA20887.0 | | | | | | | | | | | | | | 1 | | | | | | | | | | | | | 1 |
| NA21129.1 | | | | | | | | | | | | | | 1 | | | | | | | | | | | | | 1 |
| Total | 103 | 64 | 109 | 109 | 71 | 70 | 99 | 190 | 188 | 176 | 207 | 208 | 131 | 160 | 160 | 159 | 172 | 125 | 114 | 89 | 84 | 103 | 136 | 88 | 88 | 166 | 3369 |
