## Supplementary Table 3 for "Lower promoter activity of the *ST8SIA2* gene has been favored in evolving human collective brains"

Supplementary Table 3. Distribution of the CGT haplotypes

| Lineage | Haplotype | AFR | | | | | | | EUR | | | | | SAS | | | | | EAS | | | | | AMR | | | | Total |
|---|---|---|---|---|---|---|---|---|---|---|---|---|---|---|---|---|---|---|---|---|---|---|---|---|---|---|---|---|
| | | ACB | ASW | ESN | GWD | LWK | MSL | YRI | CEU | FIN | GBR | IBS | TSI | BEB | GIH | ITU | PJL | STU | CHB | CHS | CDX | JPT | KHV | CLM | MXL | PEL | PUR | |
| CGT1 | HG03105.0 | | | 1 | | | | | | | | | | | | | | | | | | | | | | | | 1 |
| | HG02095.1 | 1 | | | | | | | | | | | | | | | | | | | | | | | | | | 1 |
| | NA19908.0 | 1 | 1 | | | | 1 | 1 | | | | | | | | | | | | | | | | | | | | 4 |
| | HG03572.1 | | | | | | 1 | | | | | | | | | | | | | | | | | | | | | 1 |
| | HG03311.0 | 1 | 1 | 1 | | | 1 | 4 | | | | | | | | | | | | | | | | | | | | 8 |
| | NA19116.1 | | | | | | | 1 | | | | | | | | | | | | | | | | | | | | 1 |
| | HG02433.0 | 1 | | | 3 | | | 4 | | | | | | | | | | | | | | | | | | | | 8 |
| | NA19468.0 | | | | | 1 | | | | | | | | | | | | | | | | | | | | | | 1 |
| | HG01073.1 | | | | 4 | 6 | 1 | 1 | | | | | | | | | | | | | | | | | | | 1 | 13 |
| | HG03100.1 | | | 1 | 1 | | | 1 | | | | | | | | | | | | | | | | | | | | 3 |
| | HG02095.0 | 1 | | | | | | | | | | | | | | | | | | | | | | | | | | 1 |
| | NA19472.1 | | | | | 1 | | | | | | | | | | | | | | | | | | | | | | 1 |
| | NA18909.0 | | | | | | | 1 | | | | | | | | | | | | | | | | | | | | 1 |
| | HG03437.1 | | | | | | 1 | | | | | | | | | | | | | | | | | | | | | 1 |
| CGT2 | NA19664.0 | | | | | | | | | | | | | | | | | | | | | | | 1 | 1 | | | 2 |
| | HG03667.1 | 2 | | | | | | | | | | 1 | | 2 | 2 | | 1 | | 2 | 1 | 2 | 2 | 1 | | | | | 16 |
| | HG03772.1 | | | | | | | | | | | | | | | 1 | | | | | | | | | | | | 1 |
| | HG04042.1 | | | | | | | | | | | | | | 3 | | | 1 | | | | | | 2 | | | 1 | 7 |
| | HG02494.1 | | | | | | | | | | | | | | | | 1 | | | | | | | | | | 1 | 2 |
| | HG01455.0 | | | | | | | | | | | | | | | | | | | | | | | 1 | | | | 1 |
| Total | | 7 | 2 | 3 | 8 | 8 | 5 | 13 | | | | 1 | | 2 | 5 | 1 | 2 | 1 | 2 | 1 | 2 | 2 | 1 | 4 | 1 | | 3 | 74 |
