## Supplementary Table 4 for "Lower promoter activity of the *ST8SIA2* gene has been favored in evolving human collective brains"

Supplementary Table 4. Matrix of site differences among the CGT haplotypes

| | HG03105.0 | HG02095.1 | NA19908.0 | HG03572.1 | HG03311.0 | NA19116.1 | HG02433.0 | NA19468.0 | HG01073.1 | HG03100.1 | HG02095.0 | NA19472.1 | NA18909.0 | HG03437.1 | NA19664.0 | HG03667.1 | HG03772.1 | HG04042.1 | HG02494.1 | HG01455.0 |
|---|---|---|---|---|---|---|---|---|---|---|---|---|---|---|---|---|---|---|---|---|
| HG03105.0 | | | | | | | | | | | | | | | | | | | | |
| HG02095.1 | 1 | | | | | | | | | | | | | | | | | | | |
| NA19908.0 | 1 | 0 | | | | | | | | | | | | | | | | | | |
| HG03572.1 | 1 | 0 | 0 | | | | | | | | | | | | | | | | | |
| HG03311.0 | 1 | 0 | 0 | 0 | | | | | | | | | | | | | | | | |
| NA19116.1 | 1 | 0 | 0 | 0 | 0 | | | | | | | | | | | | | | | |
| HG02433.0 | 1 | 0 | 0 | 0 | 0 | 0 | | | | | | | | | | | | | | |
| NA19468.0 | 4 | 5 | 5 | 5 | 5 | 5 | 5 | | | | | | | | | | | | | |
| HG01073.11 | 4 | 5 | 5 | 5 | 5 | 5 | 5 | 0 | | | | | | | | | | | | |
| HG03100. | 4 | 5 | 5 | 5 | 5 | 5 | 5 | 0 | 0 | | | | | | | | | | | |
| HG02095.0 | 3 | 4 | 4 | 4 | 4 | 4 | 4 | 3 | 3 | 3 | | | | | | | | | | |
| NA19472.1 | 15 | 16 | 16 | 16 | 16 | 16 | 16 | 11 | 11 | 11 | 14 | | | | | | | | | |
| NA18909.0 | 15 | 16 | 16 | 16 | 16 | 16 | 16 | 11 | 11 | 11 | 14 | 0 | | | | | | | | |
| HG03437.1 | 15 | 14 | 14 | 14 | 14 | 14 | 14 | 13 | 13 | 13 | 14 | 16 | 16 | | | | | | | |
| NA19664.0 | 14 | 15 | 15 | 15 | 15 | 15 | 15 | 12 | 12 | 12 | 13 | 13 | 13 | 11 | | | | | | |
| HG03667.1 | 15 | 14 | 14 | 14 | 14 | 14 | 14 | 13 | 13 | 13 | 14 | 14 | 14 | 10 | 1 | | | | | |
| HG03772.1 | 15 | 14 | 14 | 14 | 14 | 14 | 14 | 13 | 13 | 13 | 14 | 14 | 14 | 10 | 1 | 0 | | | | |
| HG04042.1 | 15 | 14 | 14 | 14 | 14 | 14 | 14 | 13 | 13 | 13 | 14 | 14 | 14 | 10 | 1 | 0 | 0 | | | |
| HG02494.1 | 14 | 15 | 15 | 15 | 15 | 15 | 15 | 12 | 12 | 12 | 13 | 13 | 13 | 11 | 0 | 1 | 1 | 1 | | |
| HG01455.0 | 14 | 15 | 15 | 15 | 15 | 15 | 15 | 12 | 12 | 12 | 13 | 13 | 13 | 11 | 0 | 1 | 1 | 1 | 0 | |
