## Supplementary Table 5 for "Lower promoter activity of the *ST8SIA2* gene has been favored in evolving human collective brains"

Supplementary Table 5. Matrix of site differences among the TGT haplotypes from SAS

| | HG00125.0 | HG00132.1 | HG00409.1 | HG00590.1 | HG00707.1 | HG02494.0 | HG02660.0 | HG02789.0 | HG03615.0 | HG03765.1 | HG03779.1 | HG03782.1 | HG03809.0 | HG04003.1 | HG04020.0 | NA20867.0 | NA20884.0 | NA20901.1 |
|---|---|---|---|---|---|---|---|---|---|---|---|---|---|---|---|---|---|---|
| HG00125.0 | | | | | | | | | | | | | | | | | | |
| HG00132.1 | 1 | | | | | | | | | | | | | | | | | |
| HG00409.1 | 2 | 1 | | | | | | | | | | | | | | | | |
| HG00590.1 | 2 | 1 | 2 | | | | | | | | | | | | | | | |
| HG00707.1 | 3 | 2 | 3 | 1 | | | | | | | | | | | | | | |
| HG02494.0 | 2 | 1 | 2 | 2 | 3 | | | | | | | | | | | | | |
| HG02660.0 | 8 | 7 | 8 | 8 | 9 | 8 | | | | | | | | | | | | |
| HG02789.0 | 2 | 3 | 4 | 2 | 3 | 4 | 10 | | | | | | | | | | | |
| HG03615.0 | 2 | 3 | 4 | 2 | 1 | 4 | 10 | 2 | | | | | | | | | | |
| HG03765.1 | 2 | 1 | 2 | 2 | 3 | 2 | 8 | 4 | 4 | | | | | | | | | |
| HG03779.1 | 12 | 13 | 14 | 14 | 15 | 14 | 20 | 14 | 14 | 14 | | | | | | | | |
| HG03782.1 | 1 | 2 | 3 | 3 | 4 | 1 | 9 | 3 | 3 | 3 | 13 | | | | | | | |
| HG03809.0 | 22 | 21 | 22 | 22 | 21 | 22 | 26 | 22 | 22 | 22 | 18 | 23 | | | | | | |
| HG04003.1 | 2 | 1 | 2 | 2 | 3 | 2 | 8 | 4 | 4 | 2 | 14 | 3 | 22 | | | | | |
| HG04020.0 | 2 | 1 | 2 | 2 | 3 | 2 | 6 | 4 | 4 | 2 | 14 | 3 | 20 | 2 | | | | |
| NA20867.0 | 2 | 1 | 2 | 2 | 3 | 2 | 8 | 4 | 4 | 2 | 14 | 3 | 22 | 2 | 2 | | | |
| NA20884.0 | 3 | 2 | 3 | 3 | 4 | 3 | 9 | 5 | 5 | 3 | 15 | 4 | 23 | 3 | 3 | 3 | | |
| NA20901.1 | 2 | 1 | 2 | 2 | 3 | 2 | 8 | 4 | 4 | 2 | 14 | 3 | 22 | 2 | 2 | 2 | 3 | |
