## Supplementary Table 6 for "Lower promoter activity of the *ST8SIA2* gene has been favored in evolving human collective brains"

Supplementary Table 6. Matrix of site differences among the TCT haplotypes from SAS

| | HG00096.0 | HG00096.1 | HG00097.1 | HG00099.0 | HG00099.1 | HG00100.0 | HG00100.1 | HG00105.1 | HG00109.1 | HG00110.1 | HG00114.0 | HG00117.1 | HG00123.1 | HG00129.1 | HG00130.0 | HG00131.0 | HG00140.1 | HG00149.0 | HG00182.0 | HG01121.0 | HG01357.1 | HG01589.0 | HG01991.1 | HG02127.1 | HG02600.0 | HG02654.0 | HG02687.0 | HG02734.1 | HG02793.1 | HG03006.1 | HG03015.1 | HG03018.1 | HG03228.1 | HG03229.1 | HG03428.1 | HG03634.1 | HG03681.0 | HG03686.1 | HG03697.1 | HG03706.0 | HG03713.0 | HG03717.1 | HG03722.1 | HG03729.1 | HG03730.0 | HG03742.1 | HG03752.1 | HG03765.0 | HG03836.0 | HG03920.0 | HG03965.0 | HG04001.0 | HG04134.0 | HG04155.1 | HG04194.1 | NA20503.1 | NA20887.0 | NA21129.1 |
|---|---|---|---|---|---|---|---|---|---|---|---|---|---|---|---|---|---|---|---|---|---|---|---|---|---|---|---|---|---|---|---|---|---|---|---|---|---|---|---|---|---|---|---|---|---|---|---|---|---|---|---|---|---|---|---|---|---|---|
| HG00096.0 |
| HG00096.1 | 30 |
| HG00097.1 | 29 | 1 |
| HG00099.0 | 31 | 3 | 2 |
| HG00099.1 | 28 | 2 | 1 | 3 |
| HG00100.0 | 30 | 2 | 1 | 3 | 2 |
| HG00100.1 | 2 | 28 | 27 | 29 | 28 | 28 |
| HG00105.1 | 30 | 2 | 1 | 1 | 2 | 2 | 28 |
| HG00109.1 | 31 | 3 | 2 | 2 | 3 | 3 | 29 | 3 |
| HG00110.1 | 1 | 29 | 28 | 30 | 27 | 29 | 1 | 29 | 30 |
| HG00114.0 | 1 | 29 | 28 | 30 | 29 | 29 | 1 | 29 | 30 | 2 |
| HG00117.1 | 29 | 3 | 2 | 2 | 1 | 3 | 29 | 1 | 4 | 28 | 30 |
| HG00123.1 | 1 | 29 | 30 | 32 | 29 | 31 | 3 | 31 | 32 | 2 | 2 | 30 |
| HG00129.1 | 5 | 27 | 26 | 28 | 27 | 27 | 3 | 27 | 28 | 4 | 4 | 28 | 6 |
| HG00130.0 | 30 | 2 | 1 | 1 | 2 | 2 | 28 | 2 | 1 | 29 | 29 | 3 | 31 | 27 |
| HG00131.0 | 31 | 3 | 2 | 4 | 3 | 3 | 29 | 3 | 4 | 30 | 30 | 4 | 32 | 28 | 3 |
| HG00140.1 | 32 | 4 | 3 | 3 | 4 | 4 | 30 | 2 | 5 | 31 | 31 | 3 | 33 | 29 | 4 | 5 |
| HG00149.0 | 30 | 2 | 1 | 3 | 2 | 2 | 28 | 2 | 3 | 29 | 29 | 3 | 31 | 27 | 2 | 3 | 4 |
| HG00182.0 | 4 | 26 | 25 | 27 | 26 | 26 | 2 | 26 | 27 | 3 | 3 | 27 | 5 | 1 | 26 | 27 | 28 | 26 |
| HG01121.0 | 27 | 3 | 2 | 4 | 1 | 3 | 29 | 3 | 4 | 28 | 28 | 2 | 28 | 28 | 3 | 4 | 5 | 3 | 27 |
| HG01357.1 | 30 | 2 | 1 | 3 | 2 | 2 | 28 | 2 | 3 | 29 | 29 | 3 | 31 | 27 | 2 | 1 | 4 | 2 | 26 | 3 |
| HG01589.0 | 31 | 3 | 2 | 2 | 3 | 3 | 29 | 3 | 2 | 30 | 30 | 4 | 32 | 28 | 1 | 4 | 5 | 3 | 27 | 4 | 3 |
| HG01991.1 | 28 | 2 | 1 | 3 | 2 | 2 | 28 | 2 | 3 | 29 | 27 | 3 | 29 | 27 | 2 | 3 | 4 | 2 | 26 | 1 | 2 | 3 |
| HG02127.1 | 33 | 13 | 14 | 16 | 15 | 15 | 31 | 15 | 16 | 32 | 32 | 16 | 32 | 30 | 15 | 16 | 17 | 15 | 29 | 16 | 15 | 16 | 15 |
| HG02600.0 | 29 | 3 | 2 | 2 | 3 | 3 | 29 | 3 | 2 | 30 | 28 | 4 | 30 | 28 | 1 | 4 | 5 | 3 | 27 | 2 | 3 | 2 | 1 | 16 |
| HG02654.0 | 32 | 4 | 3 | 1 | 4 | 4 | 30 | 2 | 3 | 31 | 31 | 3 | 33 | 29 | 2 | 5 | 4 | 4 | 28 | 5 | 4 | 3 | 4 | 15 | 3 |
| HG02687.0 | 30 | 4 | 5 | 7 | 6 | 6 | 28 | 6 | 7 | 29 | 29 | 7 | 29 | 27 | 6 | 7 | 8 | 6 | 26 | 7 | 6 | 7 | 6 | 9 | 7 | 8 |
| HG02734.1 | 19 | 27 | 28 | 30 | 27 | 29 | 19 | 29 | 30 | 18 | 20 | 28 | 18 | 20 | 29 | 30 | 31 | 29 | 19 | 28 | 29 | 30 | 29 | 16 | 30 | 31 | 23 |
| HG02793.1 | 31 | 3 | 2 | 2 | 3 | 3 | 29 | 1 | 4 | 30 | 30 | 2 | 32 | 28 | 3 | 4 | 3 | 3 | 27 | 4 | 3 | 4 | 3 | 14 | 4 | 3 | 7 | 28 |
| HG03006.1 | 32 | 2 | 3 | 3 | 4 | 4 | 30 | 4 | 3 | 31 | 31 | 5 | 31 | 29 | 2 | 5 | 6 | 4 | 28 | 5 | 4 | 1 | 4 | 15 | 3 | 4 | 6 | 29 | 5 |
| HG03015.1 | 31 | 3 | 2 | 2 | 3 | 1 | 29 | 3 | 2 | 30 | 30 | 4 | 32 | 28 | 1 | 4 | 5 | 3 | 27 | 4 | 3 | 2 | 3 | 16 | 2 | 3 | 7 | 30 | 4 | 3 |
| HG03018.1 | 29 | 5 | 4 | 6 | 5 | 5 | 27 | 5 | 6 | 28 | 28 | 6 | 30 | 26 | 5 | 6 | 7 | 5 | 25 | 6 | 5 | 6 | 5 | 14 | 6 | 7 | 5 | 28 | 6 | 7 | 6 |
| HG03228.1 | 2 | 30 | 31 | 31 | 30 | 32 | 4 | 32 | 31 | 3 | 3 | 31 | 1 | 7 | 30 | 33 | 34 | 32 | 6 | 29 | 32 | 31 | 30 | 33 | 29 | 32 | 30 | 19 | 33 | 30 | 31 | 31 |
| HG03229.1 | 6 | 26 | 25 | 27 | 26 | 26 | 4 | 26 | 27 | 5 | 5 | 27 | 7 | 1 | 26 | 27 | 28 | 26 | 2 | 27 | 26 | 27 | 26 | 29 | 27 | 28 | 26 | 21 | 27 | 28 | 27 | 25 | 8 |
| HG03428.1 | 30 | 2 | 1 | 3 | 2 | 2 | 28 | 2 | 3 | 29 | 29 | 3 | 31 | 27 | 2 | 3 | 4 | 2 | 26 | 3 | 2 | 3 | 2 | 15 | 3 | 4 | 6 | 29 | 3 | 4 | 3 | 5 | 32 | 26 |
| HG03634.1 | 30 | 4 | 3 | 3 | 2 | 4 | 30 | 2 | 5 | 29 | 31 | 1 | 31 | 29 | 4 | 5 | 4 | 4 | 28 | 3 | 4 | 5 | 4 | 15 | 5 | 4 | 8 | 27 | 1 | 6 | 5 | 7 | 32 | 28 | 4 |
| HG03681.0 | 30 | 4 | 3 | 1 | 4 | 4 | 30 | 2 | 3 | 31 | 29 | 3 | 31 | 29 | 2 | 5 | 4 | 4 | 28 | 3 | 4 | 3 | 2 | 17 | 1 | 2 | 8 | 31 | 3 | 4 | 3 | 7 | 30 | 28 | 4 | 4 |
| HG03686.1 | 30 | 2 | 1 | 3 | 2 | 2 | 28 | 2 | 3 | 29 | 29 | 3 | 31 | 27 | 2 | 3 | 4 | 2 | 26 | 3 | 2 | 1 | 2 | 15 | 3 | 4 | 6 | 29 | 3 | 2 | 3 | 5 | 32 | 26 | 2 | 4 | 4 |
| HG03697.1 | 30 | 4 | 3 | 3 | 4 | 4 | 28 | 4 | 3 | 29 | 29 | 5 | 31 | 27 | 2 | 5 | 6 | 4 | 26 | 5 | 4 | 1 | 4 | 15 | 3 | 4 | 8 | 29 | 5 | 2 | 3 | 7 | 30 | 26 | 4 | 6 | 4 | 2 |
| HG03706.0 | 31 | 3 | 2 | 2 | 3 | 3 | 29 | 3 | 2 | 30 | 30 | 4 | 32 | 28 | 1 | 4 | 3 | 3 | 27 | 4 | 3 | 2 | 3 | 16 | 2 | 3 | 7 | 30 | 4 | 3 | 2 | 6 | 31 | 27 | 3 | 5 | 3 | 3 | 3 |
| HG03713.0 | 24 | 8 | 7 | 9 | 8 | 6 | 22 | 8 | 9 | 23 | 23 | 9 | 25 | 21 | 8 | 9 | 10 | 8 | 20 | 9 | 8 | 9 | 8 | 15 | 9 | 10 | 8 | 29 | 9 | 10 | 7 | 7 | 26 | 20 | 8 | 10 | 10 | 8 | 10 | 9 |
| HG03717.1 | 11 | 21 | 20 | 20 | 21 | 21 | 9 | 21 | 20 | 10 | 10 | 22 | 12 | 8 | 19 | 22 | 23 | 21 | 7 | 22 | 21 | 20 | 21 | 30 | 20 | 21 | 25 | 20 | 22 | 21 | 20 | 24 | 11 | 9 | 21 | 23 | 21 | 21 | 19 | 20 | 27 |
| HG03722.1 | 10 | 20 | 19 | 21 | 20 | 20 | 8 | 20 | 21 | 9 | 9 | 21 | 11 | 7 | 20 | 21 | 22 | 20 | 6 | 21 | 20 | 21 | 20 | 29 | 21 | 22 | 24 | 19 | 21 | 22 | 21 | 23 | 12 | 8 | 20 | 22 | 22 | 20 | 20 | 21 | 26 | 1 |
| HG03729.1 | 15 | 15 | 14 | 16 | 15 | 15 | 13 | 15 | 16 | 14 | 14 | 16 | 16 | 12 | 15 | 16 | 17 | 15 | 11 | 16 | 15 | 16 | 15 | 22 | 16 | 17 | 15 | 22 | 16 | 17 | 16 | 14 | 17 | 13 | 15 | 17 | 17 | 15 | 17 | 16 | 9 | 18 | 17 |
| HG03730.0 | 2 | 30 | 29 | 29 | 30 | 30 | 2 | 30 | 29 | 3 | 1 | 31 | 3 | 5 | 28 | 31 | 32 | 30 | 4 | 29 | 30 | 29 | 28 | 33 | 27 | 30 | 30 | 21 | 31 | 30 | 29 | 29 | 2 | 6 | 30 | 32 | 28 | 30 | 28 | 29 | 24 | 9 | 10 | 15 |
| HG03742.1 | 32 | 4 | 3 | 1 | 4 | 4 | 30 | 2 | 3 | 31 | 31 | 3 | 33 | 29 | 2 | 5 | 4 | 4 | 28 | 5 | 4 | 3 | 4 | 15 | 3 | 2 | 8 | 29 | 1 | 4 | 3 | 7 | 32 | 28 | 4 | 2 | 2 | 4 | 4 | 3 | 10 | 21 | 22 | 17 | 30 |
| HG03752.1 | 29 | 3 | 2 | 4 | 3 | 3 | 29 | 3 | 4 | 30 | 28 | 4 | 30 | 28 | 3 | 4 | 5 | 3 | 27 | 2 | 3 | 2 | 1 | 16 | 2 | 5 | 7 | 30 | 4 | 3 | 4 | 6 | 31 | 27 | 3 | 5 | 3 | 1 | 3 | 4 | 9 | 22 | 21 | 16 | 29 | 5 |
| HG03765.0 | 31 | 5 | 4 | 2 | 5 | 5 | 29 | 3 | 4 | 30 | 30 | 4 | 32 | 28 | 3 | 6 | 5 | 5 | 27 | 6 | 5 | 4 | 5 | 14 | 4 | 3 | 9 | 28 | 2 | 5 | 4 | 8 | 31 | 27 | 5 | 3 | 3 | 5 | 3 | 4 | 11 | 20 | 21 | 18 | 29 | 1 | 6 |
| HG03836.0 | 31 | 3 | 2 | 2 | 3 | 3 | 29 | 3 | 2 | 30 | 30 | 4 | 32 | 28 | 1 | 4 | 5 | 1 | 27 | 4 | 3 | 2 | 3 | 16 | 2 | 3 | 7 | 30 | 4 | 3 | 2 | 6 | 31 | 27 | 3 | 5 | 3 | 3 | 3 | 2 | 9 | 20 | 21 | 16 | 29 | 3 | 4 | 4 |
| HG03920.0 | 27 | 5 | 4 | 4 | 3 | 5 | 27 | 5 | 4 | 26 | 28 | 4 | 28 | 26 | 3 | 6 | 7 | 5 | 25 | 4 | 5 | 4 | 5 | 18 | 4 | 5 | 9 | 26 | 6 | 5 | 4 | 8 | 27 | 25 | 5 | 5 | 5 | 5 | 5 | 4 | 11 | 18 | 19 | 14 | 27 | 5 | 6 | 6 | 4 |
| HG03965.0 | 4 | 28 | 29 | 31 | 30 | 30 | 2 | 30 | 31 | 3 | 3 | 31 | 3 | 5 | 30 | 31 | 30 | 30 | 4 | 31 | 30 | 31 | 30 | 31 | 31 | 32 | 28 | 19 | 31 | 30 | 31 | 29 | 4 | 6 | 30 | 32 | 32 | 30 | 30 | 31 | 24 | 11 | 10 | 15 | 4 | 32 | 31 | 31 | 31 | 29 |
| HG04001.0 | 29 | 3 | 2 | 4 | 3 | 3 | 27 | 3 | 4 | 28 | 28 | 4 | 30 | 26 | 3 | 4 | 5 | 1 | 25 | 4 | 3 | 4 | 3 | 16 | 4 | 5 | 7 | 28 | 4 | 5 | 4 | 6 | 31 | 25 | 3 | 5 | 5 | 3 | 5 | 4 | 9 | 20 | 19 | 14 | 29 | 5 | 4 | 6 | 2 | 4 | 29 |
| HG04134.0 | 2 | 30 | 29 | 29 | 28 | 30 | 2 | 30 | 29 | 1 | 3 | 29 | 3 | 5 | 28 | 31 | 32 | 30 | 4 | 29 | 30 | 29 | 30 | 33 | 29 | 30 | 30 | 19 | 31 | 30 | 29 | 29 | 2 | 6 | 30 | 30 | 30 | 30 | 28 | 29 | 24 | 9 | 10 | 15 | 2 | 30 | 31 | 29 | 29 | 25 | 4 | 29 |
| HG04155.1 | 30 | 4 | 3 | 3 | 4 | 4 | 28 | 2 | 5 | 29 | 29 | 3 | 31 | 27 | 4 | 5 | 4 | 4 | 26 | 5 | 4 | 3 | 4 | 17 | 5 | 4 | 8 | 29 | 3 | 4 | 5 | 7 | 32 | 26 | 4 | 4 | 4 | 2 | 4 | 5 | 10 | 21 | 20 | 15 | 30 | 4 | 3 | 5 | 5 | 5 | 30 | 5 | 30 |
| HG04194.1 | 18 | 28 | 29 | 31 | 28 | 30 | 20 | 30 | 31 | 19 | 19 | 29 | 17 | 21 | 30 | 31 | 32 | 30 | 20 | 27 | 30 | 31 | 28 | 17 | 29 | 32 | 24 | 1 | 29 | 30 | 31 | 29 | 18 | 22 | 30 | 28 | 30 | 30 | 30 | 31 | 30 | 21 | 20 | 23 | 20 | 30 | 29 | 29 | 31 | 27 | 20 | 29 | 20 | 30 |
| NA20503.1 | 32 | 8 | 7 | 9 | 8 | 8 | 30 | 8 | 9 | 31 | 31 | 9 | 33 | 29 | 8 | 9 | 10 | 8 | 28 | 9 | 8 | 9 | 8 | 17 | 9 | 10 | 8 | 31 | 9 | 10 | 9 | 3 | 34 | 28 | 8 | 10 | 10 | 8 | 10 | 9 | 10 | 27 | 26 | 17 | 32 | 10 | 9 | 11 | 9 | 11 | 32 | 9 | 32 | 10 | 32 |
| NA20887.0 | 8 | 22 | 21 | 23 | 22 | 22 | 6 | 22 | 23 | 7 | 7 | 23 | 9 | 5 | 22 | 23 | 24 | 22 | 4 | 23 | 22 | 23 | 22 | 25 | 23 | 24 | 22 | 23 | 23 | 24 | 23 | 21 | 10 | 6 | 22 | 24 | 24 | 22 | 22 | 23 | 16 | 11 | 10 | 7 | 8 | 24 | 23 | 23 | 23 | 21 | 8 | 21 | 8 | 22 | 24 | 24 |
| NA21129.1 | 33 | 9 | 8 | 10 | 9 | 9 | 31 | 9 | 10 | 32 | 32 | 10 | 34 | 30 | 9 | 10 | 11 | 9 | 29 | 10 | 9 | 8 | 9 | 18 | 10 | 11 | 9 | 32 | 10 | 9 | 10 | 4 | 35 | 29 | 9 | 11 | 11 | 7 | 9 | 10 | 11 | 28 | 27 | 18 | 33 | 11 | 8 | 12 | 10 | 12 | 33 | 10 | 33 | 9 | 33 | 1 | 25 |
