## Supplementary Table 7 for "Lower promoter activity of the *ST8SIA2* gene has been favored in evolving human collective brains"

Supplementary Table 7. Matrix of site differences among the CGC haplotypes from SAS

| | HG03817.1 | HG00356.0 | HG02069.0 | HG03809.1 | HG03870.1 | HG02490.0 | HG03868.0 |
|---|---|---|---|---|---|---|---|
| HG03817.1 | | | | | | | |
| HG00356.0 | 1 | | | | | | |
| HG02069.0 | 3 | 2 | | | | | |
| HG03809.1 | 23 | 22 | 24 | | | | |
| HG03870.1 | 1 | 0 | 2 | 22 | | | |
| HG02490.0 | 1 | 0 | 2 | 22 | 0 | | |
| HG03868.0 | 1 | 0 | 2 | 22 | 0 | 0 | |
