## Supplementary Table 8 for "Lower promoter activity of the *ST8SIA2* gene has been favored in evolving human collective brains"

Supplementary Table 8. Inference of promoter type composition and PPA of AH population.

| Promoter type frequency | | | Genotype frequency | | | | | | PPA | Difference of PPA relative to AFR | Difference of PPA relative to LWK |
|---|---|---|---|---|---|---|---|---|---|---|---|
| CGT | TGT | TCT | CGT/CGT | CGT/TGT | CGT/TCT | TGT/TGT | TCT/TCT | TGT/TCT | | | |
| 0.717 | 0.142 | 0.141 | 0.514 | 0.204 | 0.202 | 0.020 | 0.020 | 0.040 | 2.891 | 1.010 | 0.944 |
| 0.717 | 0.152 | 0.131 | 0.514 | 0.218 | 0.188 | 0.023 | 0.017 | 0.040 | 2.906 | 1.016 | 0.949 |
| 0.717 | 0.162 | 0.121 | 0.514 | 0.232 | 0.174 | 0.026 | 0.015 | 0.039 | 2.921 | 1.021 | 0.954 |
| 0.717 | 0.172 | 0.111 | 0.514 | 0.247 | 0.159 | 0.030 | 0.012 | 0.038 | 2.936 | 1.026 | 0.959 |
| 0.717 | 0.182 | 0.101 | 0.514 | 0.261 | 0.145 | 0.033 | 0.010 | 0.037 | 2.951 | 1.031 | 0.963 |
| 0.717 | 0.192 | 0.091 | 0.514 | 0.275 | 0.130 | 0.037 | 0.008 | 0.035 | 2.966 | 1.037 | 0.968 |
| 0.717 | 0.202 | 0.081 | 0.514 | 0.290 | 0.116 | 0.041 | 0.007 | 0.033 | 2.981 | 1.042 | 0.973 |
| 0.717 | 0.212 | 0.071 | 0.514 | 0.304 | 0.102 | 0.045 | 0.005 | 0.030 | 2.996 | 1.047 | 0.978 |
| 0.717 | 0.222 | 0.061 | 0.514 | 0.318 | 0.087 | 0.049 | 0.004 | 0.027 | 3.011 | 1.052 | 0.983 |
| 0.717 | 0.232 | 0.051 | 0.514 | 0.333 | 0.073 | 0.054 | 0.003 | 0.024 | 3.026 | 1.058 | 0.988 |
| 0.717 | 0.242 | 0.041 | 0.514 | 0.347 | 0.059 | 0.059 | 0.002 | 0.020 | 3.041 | 1.063 | 0.993 |
| 0.717 | 0.252 | 0.031 | 0.514 | 0.361 | 0.044 | 0.064 | 0.001 | 0.016 | 3.056 | 1.068 | 0.998 |
| 0.717 | 0.262 | 0.021 | 0.514 | 0.376 | 0.030 | 0.069 | 0.000 | 0.011 | 3.071 | 1.073 | 1.003 |
| 0.717 | 0.272 | 0.011 | 0.514 | 0.390 | 0.016 | 0.074 | 0.000 | 0.006 | 3.086 | 1.079 | 1.008 |
| 0.72 | 0.15 | 0.13 | 0.518 | 0.216 | 0.187 | 0.023 | 0.017 | 0.039 | 2.905 | 1.015 | 0.948 |
| 0.72 | 0.16 | 0.12 | 0.518 | 0.230 | 0.173 | 0.026 | 0.014 | 0.038 | 2.920 | 1.021 | 0.953 |
| 0.72 | 0.17 | 0.11 | 0.518 | 0.245 | 0.158 | 0.029 | 0.012 | 0.037 | 2.935 | 1.026 | 0.958 |
| 0.72 | 0.18 | 0.1 | 0.518 | 0.259 | 0.144 | 0.032 | 0.010 | 0.036 | 2.950 | 1.031 | 0.963 |
| 0.72 | 0.19 | 0.09 | 0.518 | 0.274 | 0.130 | 0.036 | 0.008 | 0.034 | 2.965 | 1.036 | 0.968 |
| 0.72 | 0.2 | 0.08 | 0.518 | 0.288 | 0.115 | 0.040 | 0.006 | 0.032 | 2.980 | 1.042 | 0.973 |
| 0.72 | 0.21 | 0.07 | 0.518 | 0.302 | 0.101 | 0.044 | 0.005 | 0.029 | 2.995 | 1.047 | 0.978 |
| 0.72 | 0.22 | 0.06 | 0.518 | 0.317 | 0.086 | 0.048 | 0.004 | 0.026 | 3.010 | 1.052 | 0.983 |
| 0.72 | 0.23 | 0.05 | 0.518 | 0.331 | 0.072 | 0.053 | 0.003 | 0.023 | 3.025 | 1.057 | 0.988 |
| 0.72 | 0.24 | 0.04 | 0.518 | 0.346 | 0.058 | 0.058 | 0.002 | 0.019 | 3.041 | 1.063 | 0.993 |
| 0.72 | 0.25 | 0.03 | 0.518 | 0.360 | 0.043 | 0.063 | 0.001 | 0.015 | 3.056 | 1.068 | 0.998 |
| 0.72 | 0.26 | 0.02 | 0.518 | 0.374 | 0.029 | 0.068 | 0.000 | 0.010 | 3.071 | 1.073 | 1.002 |
| 0.72 | 0.27 | 0.01 | 0.518 | 0.389 | 0.014 | 0.073 | 0.000 | 0.005 | 3.086 | 1.078 | 1.007 |
| 0.73 | 0.14 | 0.13 | 0.533 | 0.204 | 0.190 | 0.020 | 0.017 | 0.036 | 2.898 | 1.013 | 0.946 |
| 0.73 | 0.15 | 0.12 | 0.533 | 0.219 | 0.175 | 0.023 | 0.014 | 0.036 | 2.913 | 1.018 | 0.951 |
| 0.73 | 0.16 | 0.11 | 0.533 | 0.234 | 0.161 | 0.026 | 0.012 | 0.035 | 2.928 | 1.023 | 0.956 |
| 0.73 | 0.17 | 0.1 | 0.533 | 0.248 | 0.146 | 0.029 | 0.010 | 0.034 | 2.943 | 1.028 | 0.961 |
| 0.73 | 0.18 | 0.09 | 0.533 | 0.263 | 0.131 | 0.032 | 0.008 | 0.032 | 2.958 | 1.034 | 0.966 |
| 0.73 | 0.19 | 0.08 | 0.533 | 0.277 | 0.117 | 0.036 | 0.006 | 0.030 | 2.973 | 1.039 | 0.971 |
| 0.73 | 0.2 | 0.07 | 0.533 | 0.292 | 0.102 | 0.040 | 0.005 | 0.028 | 2.988 | 1.044 | 0.975 |
| 0.73 | 0.21 | 0.06 | 0.533 | 0.307 | 0.088 | 0.044 | 0.004 | 0.025 | 3.003 | 1.049 | 0.980 |
| 0.73 | 0.22 | 0.05 | 0.533 | 0.321 | 0.073 | 0.048 | 0.003 | 0.022 | 3.018 | 1.055 | 0.985 |
| 0.73 | 0.23 | 0.04 | 0.533 | 0.336 | 0.058 | 0.053 | 0.002 | 0.018 | 3.033 | 1.060 | 0.990 |
| 0.73 | 0.24 | 0.03 | 0.533 | 0.350 | 0.044 | 0.058 | 0.001 | 0.014 | 3.048 | 1.065 | 0.995 |
| 0.73 | 0.25 | 0.02 | 0.533 | 0.365 | 0.029 | 0.063 | 0.000 | 0.010 | 3.063 | 1.070 | 1.000 |
| 0.73 | 0.26 | 0.01 | 0.533 | 0.380 | 0.015 | 0.068 | 0.000 | 0.005 | 3.078 | 1.076 | 1.005 |
| 0.74 | 0.14 | 0.12 | 0.548 | 0.207 | 0.178 | 0.020 | 0.014 | 0.034 | 2.905 | 1.015 | 0.948 |
| 0.74 | 0.15 | 0.11 | 0.548 | 0.222 | 0.163 | 0.023 | 0.012 | 0.033 | 2.920 | 1.021 | 0.953 |
| 0.74 | 0.16 | 0.1 | 0.548 | 0.237 | 0.148 | 0.026 | 0.010 | 0.032 | 2.935 | 1.026 | 0.958 |
| 0.74 | 0.17 | 0.09 | 0.548 | 0.252 | 0.133 | 0.029 | 0.008 | 0.031 | 2.950 | 1.031 | 0.963 |
| 0.74 | 0.18 | 0.08 | 0.548 | 0.266 | 0.118 | 0.032 | 0.006 | 0.029 | 2.966 | 1.036 | 0.968 |
| 0.74 | 0.19 | 0.07 | 0.548 | 0.281 | 0.104 | 0.036 | 0.005 | 0.027 | 2.981 | 1.042 | 0.973 |
| 0.74 | 0.2 | 0.06 | 0.548 | 0.296 | 0.089 | 0.040 | 0.004 | 0.024 | 2.996 | 1.047 | 0.978 |
| 0.74 | 0.21 | 0.05 | 0.548 | 0.311 | 0.074 | 0.044 | 0.003 | 0.021 | 3.011 | 1.052 | 0.983 |
| 0.74 | 0.22 | 0.04 | 0.548 | 0.326 | 0.059 | 0.048 | 0.002 | 0.018 | 3.026 | 1.057 | 0.988 |
| 0.74 | 0.23 | 0.03 | 0.548 | 0.340 | 0.044 | 0.053 | 0.001 | 0.014 | 3.041 | 1.063 | 0.993 |
| 0.74 | 0.24 | 0.02 | 0.548 | 0.355 | 0.030 | 0.058 | 0.000 | 0.010 | 3.056 | 1.068 | 0.998 |
| 0.74 | 0.25 | 0.01 | 0.548 | 0.370 | 0.015 | 0.063 | 0.000 | 0.005 | 3.071 | 1.073 | 1.002 |
| 0.75 | 0.13 | 0.12 | 0.563 | 0.195 | 0.180 | 0.017 | 0.014 | 0.031 | 2.898 | 1.013 | 0.946 |
| 0.75 | 0.14 | 0.11 | 0.563 | 0.210 | 0.165 | 0.020 | 0.012 | 0.031 | 2.913 | 1.018 | 0.951 |
| 0.75 | 0.15 | 0.1 | 0.563 | 0.225 | 0.150 | 0.023 | 0.010 | 0.030 | 2.928 | 1.023 | 0.956 |
| 0.75 | 0.16 | 0.09 | 0.563 | 0.240 | 0.135 | 0.026 | 0.008 | 0.029 | 2.943 | 1.028 | 0.961 |
| 0.75 | 0.17 | 0.08 | 0.563 | 0.255 | 0.120 | 0.029 | 0.006 | 0.027 | 2.958 | 1.034 | 0.966 |
| 0.75 | 0.18 | 0.07 | 0.563 | 0.270 | 0.105 | 0.032 | 0.005 | 0.025 | 2.973 | 1.039 | 0.971 |
| 0.75 | 0.19 | 0.06 | 0.563 | 0.285 | 0.090 | 0.036 | 0.004 | 0.023 | 2.988 | 1.044 | 0.976 |
| 0.75 | 0.2 | 0.05 | 0.563 | 0.300 | 0.075 | 0.040 | 0.003 | 0.020 | 3.003 | 1.049 | 0.980 |
| 0.75 | 0.21 | 0.04 | 0.563 | 0.315 | 0.060 | 0.044 | 0.002 | 0.017 | 3.018 | 1.055 | 0.985 |
| 0.75 | 0.22 | 0.03 | 0.563 | 0.330 | 0.045 | 0.048 | 0.001 | 0.013 | 3.033 | 1.060 | 0.990 |
| 0.75 | 0.23 | 0.02 | 0.563 | 0.345 | 0.030 | 0.053 | 0.000 | 0.009 | 3.048 | 1.065 | 0.995 |
| 0.75 | 0.24 | 0.01 | 0.563 | 0.360 | 0.015 | 0.058 | 0.000 | 0.005 | 3.063 | 1.071 | 1.000 |
| 0.76 | 0.13 | 0.11 | 0.578 | 0.198 | 0.167 | 0.017 | 0.012 | 0.029 | 2.906 | 1.015 | 0.949 |
| 0.76 | 0.14 | 0.1 | 0.578 | 0.213 | 0.152 | 0.020 | 0.010 | 0.028 | 2.921 | 1.021 | 0.953 |
| 0.76 | 0.15 | 0.09 | 0.578 | 0.228 | 0.137 | 0.023 | 0.008 | 0.027 | 2.936 | 1.026 | 0.958 |
| 0.76 | 0.16 | 0.08 | 0.578 | 0.243 | 0.122 | 0.026 | 0.006 | 0.026 | 2.951 | 1.031 | 0.963 |
| 0.76 | 0.17 | 0.07 | 0.578 | 0.258 | 0.106 | 0.029 | 0.005 | 0.024 | 2.966 | 1.036 | 0.968 |
| 0.76 | 0.18 | 0.06 | 0.578 | 0.274 | 0.091 | 0.032 | 0.004 | 0.022 | 2.981 | 1.042 | 0.973 |
| 0.76 | 0.19 | 0.05 | 0.578 | 0.289 | 0.076 | 0.036 | 0.003 | 0.019 | 2.996 | 1.047 | 0.978 |
| 0.76 | 0.2 | 0.04 | 0.578 | 0.304 | 0.061 | 0.040 | 0.002 | 0.016 | 3.011 | 1.052 | 0.983 |
| 0.76 | 0.21 | 0.03 | 0.578 | 0.319 | 0.046 | 0.044 | 0.001 | 0.013 | 3.026 | 1.057 | 0.988 |
| 0.76 | 0.22 | 0.02 | 0.578 | 0.334 | 0.030 | 0.048 | 0.000 | 0.009 | 3.041 | 1.063 | 0.993 |
| 0.76 | 0.23 | 0.01 | 0.578 | 0.350 | 0.015 | 0.053 | 0.000 | 0.005 | 3.056 | 1.068 | 0.998 |
| 0.77 | 0.12 | 0.11 | 0.593 | 0.185 | 0.169 | 0.014 | 0.012 | 0.026 | 2.898 | 1.013 | 0.946 |
| 0.77 | 0.13 | 0.1 | 0.593 | 0.200 | 0.154 | 0.017 | 0.010 | 0.026 | 2.913 | 1.018 | 0.951 |
| 0.77 | 0.14 | 0.09 | 0.593 | 0.216 | 0.139 | 0.020 | 0.008 | 0.025 | 2.928 | 1.023 | 0.956 |
| 0.77 | 0.15 | 0.08 | 0.593 | 0.231 | 0.123 | 0.023 | 0.006 | 0.024 | 2.943 | 1.029 | 0.961 |
| 0.77 | 0.16 | 0.07 | 0.593 | 0.246 | 0.108 | 0.026 | 0.005 | 0.022 | 2.958 | 1.034 | 0.966 |
| 0.77 | 0.17 | 0.06 | 0.593 | 0.262 | 0.092 | 0.029 | 0.004 | 0.020 | 2.973 | 1.039 | 0.971 |
| 0.77 | 0.18 | 0.05 | 0.593 | 0.277 | 0.077 | 0.032 | 0.003 | 0.018 | 2.988 | 1.044 | 0.976 |
| 0.77 | 0.19 | 0.04 | 0.593 | 0.293 | 0.062 | 0.036 | 0.002 | 0.015 | 3.003 | 1.050 | 0.980 |
| 0.77 | 0.2 | 0.03 | 0.593 | 0.308 | 0.046 | 0.040 | 0.001 | 0.012 | 3.018 | 1.055 | 0.985 |
| 0.77 | 0.21 | 0.02 | 0.593 | 0.323 | 0.031 | 0.044 | 0.000 | 0.008 | 3.033 | 1.060 | 0.990 |
| 0.77 | 0.22 | 0.01 | 0.593 | 0.339 | 0.015 | 0.048 | 0.000 | 0.004 | 3.048 | 1.065 | 0.995 |
| 0.78 | 0.12 | 0.1 | 0.608 | 0.187 | 0.156 | 0.014 | 0.010 | 0.024 | 2.906 | 1.015 | 0.949 |
| 0.78 | 0.13 | 0.09 | 0.608 | 0.203 | 0.140 | 0.017 | 0.008 | 0.023 | 2.921 | 1.021 | 0.953 |
| 0.78 | 0.14 | 0.08 | 0.608 | 0.218 | 0.125 | 0.020 | 0.006 | 0.022 | 2.936 | 1.026 | 0.958 |
| 0.78 | 0.15 | 0.07 | 0.608 | 0.234 | 0.109 | 0.023 | 0.005 | 0.021 | 2.951 | 1.031 | 0.963 |
| 0.78 | 0.16 | 0.06 | 0.608 | 0.250 | 0.094 | 0.026 | 0.004 | 0.019 | 2.966 | 1.036 | 0.968 |
| 0.78 | 0.17 | 0.05 | 0.608 | 0.265 | 0.078 | 0.029 | 0.003 | 0.017 | 2.981 | 1.042 | 0.973 |
| 0.78 | 0.18 | 0.04 | 0.608 | 0.281 | 0.062 | 0.032 | 0.002 | 0.014 | 2.996 | 1.047 | 0.978 |
| 0.78 | 0.19 | 0.03 | 0.608 | 0.296 | 0.047 | 0.036 | 0.001 | 0.011 | 3.011 | 1.052 | 0.983 |
| 0.78 | 0.2 | 0.02 | 0.608 | 0.312 | 0.031 | 0.040 | 0.000 | 0.008 | 3.026 | 1.057 | 0.988 |
| 0.78 | 0.21 | 0.01 | 0.608 | 0.328 | 0.016 | 0.044 | 0.000 | 0.004 | 3.041 | 1.063 | 0.993 |
| 0.79 | 0.11 | 0.1 | 0.624 | 0.174 | 0.158 | 0.012 | 0.010 | 0.022 | 2.898 | 1.013 | 0.946 |
| 0.79 | 0.12 | 0.09 | 0.624 | 0.190 | 0.142 | 0.014 | 0.008 | 0.022 | 2.913 | 1.018 | 0.951 |
| 0.79 | 0.13 | 0.08 | 0.624 | 0.205 | 0.126 | 0.017 | 0.006 | 0.021 | 2.928 | 1.023 | 0.956 |
| 0.79 | 0.14 | 0.07 | 0.624 | 0.221 | 0.111 | 0.020 | 0.005 | 0.020 | 2.943 | 1.029 | 0.961 |
| 0.79 | 0.15 | 0.06 | 0.624 | 0.237 | 0.095 | 0.023 | 0.004 | 0.018 | 2.958 | 1.034 | 0.966 |
| 0.79 | 0.16 | 0.05 | 0.624 | 0.253 | 0.079 | 0.026 | 0.003 | 0.016 | 2.973 | 1.039 | 0.971 |
| 0.79 | 0.17 | 0.04 | 0.624 | 0.269 | 0.063 | 0.029 | 0.002 | 0.014 | 2.988 | 1.044 | 0.976 |
| 0.79 | 0.18 | 0.03 | 0.624 | 0.284 | 0.047 | 0.032 | 0.001 | 0.011 | 3.003 | 1.050 | 0.981 |
| 0.79 | 0.19 | 0.02 | 0.624 | 0.300 | 0.032 | 0.036 | 0.000 | 0.008 | 3.018 | 1.055 | 0.985 |
| 0.79 | 0.2 | 0.01 | 0.624 | 0.316 | 0.016 | 0.040 | 0.000 | 0.004 | 3.033 | 1.060 | 0.990 |
| 0.8 | 0.11 | 0.09 | 0.640 | 0.176 | 0.144 | 0.012 | 0.008 | 0.020 | 2.906 | 1.015 | 0.949 |
| 0.8 | 0.12 | 0.08 | 0.640 | 0.192 | 0.128 | 0.014 | 0.006 | 0.019 | 2.921 | 1.021 | 0.954 |
| 0.8 | 0.13 | 0.07 | 0.640 | 0.208 | 0.112 | 0.017 | 0.005 | 0.018 | 2.936 | 1.026 | 0.958 |
| 0.8 | 0.14 | 0.06 | 0.640 | 0.224 | 0.096 | 0.020 | 0.004 | 0.017 | 2.951 | 1.031 | 0.963 |
| 0.8 | 0.15 | 0.05 | 0.640 | 0.240 | 0.080 | 0.023 | 0.003 | 0.015 | 2.966 | 1.036 | 0.968 |
| 0.8 | 0.16 | 0.04 | 0.640 | 0.256 | 0.064 | 0.026 | 0.002 | 0.013 | 2.981 | 1.042 | 0.973 |
| 0.8 | 0.17 | 0.03 | 0.640 | 0.272 | 0.048 | 0.029 | 0.001 | 0.010 | 2.996 | 1.047 | 0.978 |
| 0.8 | 0.18 | 0.02 | 0.640 | 0.288 | 0.032 | 0.032 | 0.000 | 0.007 | 3.011 | 1.052 | 0.983 |
| 0.8 | 0.19 | 0.01 | 0.640 | 0.304 | 0.016 | 0.036 | 0.000 | 0.004 | 3.026 | 1.057 | 0.988 |
| 0.81 | 0.1 | 0.09 | 0.656 | 0.162 | 0.146 | 0.010 | 0.008 | 0.018 | 2.898 | 1.013 | 0.946 |
| 0.81 | 0.11 | 0.08 | 0.656 | 0.178 | 0.130 | 0.012 | 0.006 | 0.018 | 2.913 | 1.018 | 0.951 |
| 0.81 | 0.12 | 0.07 | 0.656 | 0.194 | 0.113 | 0.014 | 0.005 | 0.017 | 2.928 | 1.023 | 0.956 |
| 0.81 | 0.13 | 0.06 | 0.656 | 0.211 | 0.097 | 0.017 | 0.004 | 0.016 | 2.943 | 1.029 | 0.961 |
| 0.81 | 0.14 | 0.05 | 0.656 | 0.227 | 0.081 | 0.020 | 0.003 | 0.014 | 2.958 | 1.034 | 0.966 |
| 0.81 | 0.15 | 0.04 | 0.656 | 0.243 | 0.065 | 0.023 | 0.002 | 0.012 | 2.973 | 1.039 | 0.971 |
| 0.81 | 0.16 | 0.03 | 0.656 | 0.259 | 0.049 | 0.026 | 0.001 | 0.010 | 2.989 | 1.044 | 0.976 |
| 0.81 | 0.17 | 0.02 | 0.656 | 0.275 | 0.032 | 0.029 | 0.000 | 0.007 | 3.004 | 1.050 | 0.981 |
| 0.81 | 0.18 | 0.01 | 0.656 | 0.292 | 0.016 | 0.032 | 0.000 | 0.004 | 3.019 | 1.055 | 0.985 |
| 0.82 | 0.1 | 0.08 | 0.672 | 0.164 | 0.131 | 0.010 | 0.006 | 0.016 | 2.906 | 1.016 | 0.949 |
| 0.82 | 0.11 | 0.07 | 0.672 | 0.180 | 0.115 | 0.012 | 0.005 | 0.015 | 2.921 | 1.021 | 0.954 |
| 0.82 | 0.12 | 0.06 | 0.672 | 0.197 | 0.098 | 0.014 | 0.004 | 0.014 | 2.936 | 1.026 | 0.958 |
| 0.82 | 0.13 | 0.05 | 0.672 | 0.213 | 0.082 | 0.017 | 0.003 | 0.013 | 2.951 | 1.031 | 0.963 |
| 0.82 | 0.14 | 0.04 | 0.672 | 0.230 | 0.066 | 0.020 | 0.002 | 0.011 | 2.966 | 1.037 | 0.968 |
| 0.82 | 0.15 | 0.03 | 0.672 | 0.246 | 0.049 | 0.023 | 0.001 | 0.009 | 2.981 | 1.042 | 0.973 |
| 0.82 | 0.16 | 0.02 | 0.672 | 0.262 | 0.033 | 0.026 | 0.000 | 0.006 | 2.996 | 1.047 | 0.978 |
| 0.82 | 0.17 | 0.01 | 0.672 | 0.279 | 0.016 | 0.029 | 0.000 | 0.003 | 3.011 | 1.052 | 0.983 |
| 0.83 | 0.09 | 0.08 | 0.689 | 0.149 | 0.133 | 0.008 | 0.006 | 0.014 | 2.898 | 1.013 | 0.946 |
| 0.83 | 0.1 | 0.07 | 0.689 | 0.166 | 0.116 | 0.010 | 0.005 | 0.014 | 2.914 | 1.018 | 0.951 |
| 0.83 | 0.11 | 0.06 | 0.689 | 0.183 | 0.100 | 0.012 | 0.004 | 0.013 | 2.929 | 1.023 | 0.956 |
| 0.83 | 0.12 | 0.05 | 0.689 | 0.199 | 0.083 | 0.014 | 0.003 | 0.012 | 2.944 | 1.029 | 0.961 |
| 0.83 | 0.13 | 0.04 | 0.689 | 0.216 | 0.066 | 0.017 | 0.002 | 0.010 | 2.959 | 1.034 | 0.966 |
| 0.83 | 0.14 | 0.03 | 0.689 | 0.232 | 0.050 | 0.020 | 0.001 | 0.008 | 2.974 | 1.039 | 0.971 |
| 0.83 | 0.15 | 0.02 | 0.689 | 0.249 | 0.033 | 0.023 | 0.000 | 0.006 | 2.989 | 1.044 | 0.976 |
| 0.83 | 0.16 | 0.01 | 0.689 | 0.266 | 0.017 | 0.026 | 0.000 | 0.003 | 3.004 | 1.050 | 0.981 |
| 0.84 | 0.09 | 0.07 | 0.706 | 0.151 | 0.118 | 0.008 | 0.005 | 0.013 | 2.906 | 1.016 | 0.949 |
| 0.84 | 0.1 | 0.06 | 0.706 | 0.168 | 0.101 | 0.010 | 0.004 | 0.012 | 2.921 | 1.021 | 0.954 |
| 0.84 | 0.11 | 0.05 | 0.706 | 0.185 | 0.084 | 0.012 | 0.003 | 0.011 | 2.936 | 1.026 | 0.959 |
| 0.84 | 0.12 | 0.04 | 0.706 | 0.202 | 0.067 | 0.014 | 0.002 | 0.010 | 2.951 | 1.031 | 0.963 |
| 0.84 | 0.13 | 0.03 | 0.706 | 0.218 | 0.050 | 0.017 | 0.001 | 0.008 | 2.966 | 1.037 | 0.968 |
| 0.84 | 0.14 | 0.02 | 0.706 | 0.235 | 0.034 | 0.020 | 0.000 | 0.006 | 2.981 | 1.042 | 0.973 |
| 0.84 | 0.15 | 0.01 | 0.706 | 0.252 | 0.017 | 0.023 | 0.000 | 0.003 | 2.996 | 1.047 | 0.978 |
| 0.85 | 0.08 | 0.07 | 0.723 | 0.136 | 0.119 | 0.006 | 0.005 | 0.011 | 2.899 | 1.013 | 0.946 |
| 0.85 | 0.09 | 0.06 | 0.723 | 0.153 | 0.102 | 0.008 | 0.004 | 0.011 | 2.914 | 1.018 | 0.951 |
| 0.85 | 0.1 | 0.05 | 0.723 | 0.170 | 0.085 | 0.010 | 0.003 | 0.010 | 2.929 | 1.023 | 0.956 |
| 0.85 | 0.11 | 0.04 | 0.723 | 0.187 | 0.068 | 0.012 | 0.002 | 0.009 | 2.944 | 1.029 | 0.961 |
| 0.85 | 0.12 | 0.03 | 0.723 | 0.204 | 0.051 | 0.014 | 0.001 | 0.007 | 2.959 | 1.034 | 0.966 |
| 0.85 | 0.13 | 0.02 | 0.723 | 0.221 | 0.034 | 0.017 | 0.000 | 0.005 | 2.974 | 1.039 | 0.971 |
| 0.85 | 0.14 | 0.01 | 0.723 | 0.238 | 0.017 | 0.020 | 0.000 | 0.003 | 2.989 | 1.044 | 0.976 |
| 0.86 | 0.08 | 0.06 | 0.740 | 0.138 | 0.103 | 0.006 | 0.004 | 0.010 | 2.906 | 1.016 | 0.949 |
| 0.86 | 0.09 | 0.05 | 0.740 | 0.155 | 0.086 | 0.008 | 0.003 | 0.009 | 2.921 | 1.021 | 0.954 |
| 0.86 | 0.1 | 0.04 | 0.740 | 0.172 | 0.069 | 0.010 | 0.002 | 0.008 | 2.936 | 1.026 | 0.959 |
| 0.86 | 0.11 | 0.03 | 0.740 | 0.189 | 0.052 | 0.012 | 0.001 | 0.007 | 2.951 | 1.031 | 0.963 |
| 0.86 | 0.12 | 0.02 | 0.740 | 0.206 | 0.034 | 0.014 | 0.000 | 0.005 | 2.966 | 1.037 | 0.968 |
| 0.86 | 0.13 | 0.01 | 0.740 | 0.224 | 0.017 | 0.017 | 0.000 | 0.003 | 2.981 | 1.042 | 0.973 |
| 0.87 | 0.07 | 0.06 | 0.757 | 0.122 | 0.104 | 0.005 | 0.004 | 0.008 | 2.899 | 1.013 | 0.946 |
| 0.87 | 0.08 | 0.05 | 0.757 | 0.139 | 0.087 | 0.006 | 0.003 | 0.008 | 2.914 | 1.018 | 0.951 |
| 0.87 | 0.09 | 0.04 | 0.757 | 0.157 | 0.070 | 0.008 | 0.002 | 0.007 | 2.929 | 1.024 | 0.956 |
| 0.87 | 0.1 | 0.03 | 0.757 | 0.174 | 0.052 | 0.010 | 0.001 | 0.006 | 2.944 | 1.029 | 0.961 |
| 0.87 | 0.11 | 0.02 | 0.757 | 0.191 | 0.035 | 0.012 | 0.000 | 0.004 | 2.959 | 1.034 | 0.966 |
| 0.87 | 0.12 | 0.01 | 0.757 | 0.209 | 0.017 | 0.014 | 0.000 | 0.002 | 2.974 | 1.039 | 0.971 |
| 0.88 | 0.07 | 0.05 | 0.774 | 0.123 | 0.088 | 0.005 | 0.003 | 0.007 | 2.906 | 1.016 | 0.949 |
| 0.88 | 0.08 | 0.04 | 0.774 | 0.141 | 0.070 | 0.006 | 0.002 | 0.006 | 2.921 | 1.021 | 0.954 |
| 0.88 | 0.09 | 0.03 | 0.774 | 0.158 | 0.053 | 0.008 | 0.001 | 0.005 | 2.936 | 1.026 | 0.959 |
| 0.88 | 0.1 | 0.02 | 0.774 | 0.176 | 0.035 | 0.010 | 0.000 | 0.004 | 2.951 | 1.031 | 0.964 |
| 0.88 | 0.11 | 0.01 | 0.774 | 0.194 | 0.018 | 0.012 | 0.000 | 0.002 | 2.966 | 1.037 | 0.968 |
| 0.89 | 0.06 | 0.05 | 0.792 | 0.107 | 0.089 | 0.004 | 0.003 | 0.006 | 2.899 | 1.013 | 0.946 |
| 0.89 | 0.07 | 0.04 | 0.792 | 0.125 | 0.071 | 0.005 | 0.002 | 0.006 | 2.914 | 1.018 | 0.951 |
| 0.89 | 0.08 | 0.03 | 0.792 | 0.142 | 0.053 | 0.006 | 0.001 | 0.005 | 2.929 | 1.024 | 0.956 |
| 0.89 | 0.09 | 0.02 | 0.792 | 0.160 | 0.036 | 0.008 | 0.000 | 0.004 | 2.944 | 1.029 | 0.961 |
| 0.89 | 0.1 | 0.01 | 0.792 | 0.178 | 0.018 | 0.010 | 0.000 | 0.002 | 2.959 | 1.034 | 0.966 |
| 0.9 | 0.06 | 0.04 | 0.810 | 0.108 | 0.072 | 0.004 | 0.002 | 0.005 | 2.906 | 1.016 | 0.949 |
| 0.9 | 0.07 | 0.03 | 0.810 | 0.126 | 0.054 | 0.005 | 0.001 | 0.004 | 2.921 | 1.021 | 0.954 |
| 0.9 | 0.08 | 0.02 | 0.810 | 0.144 | 0.036 | 0.006 | 0.000 | 0.003 | 2.937 | 1.026 | 0.959 |
| 0.9 | 0.09 | 0.01 | 0.810 | 0.162 | 0.018 | 0.008 | 0.000 | 0.002 | 2.952 | 1.031 | 0.964 |
| 0.91 | 0.05 | 0.04 | 0.828 | 0.091 | 0.073 | 0.003 | 0.002 | 0.004 | 2.899 | 1.013 | 0.946 |
| 0.91 | 0.06 | 0.03 | 0.828 | 0.109 | 0.055 | 0.004 | 0.001 | 0.004 | 2.914 | 1.018 | 0.951 |
| 0.91 | 0.07 | 0.02 | 0.828 | 0.127 | 0.036 | 0.005 | 0.000 | 0.003 | 2.929 | 1.024 | 0.956 |
| 0.91 | 0.08 | 0.01 | 0.828 | 0.146 | 0.018 | 0.006 | 0.000 | 0.002 | 2.944 | 1.029 | 0.961 |
| 0.92 | 0.05 | 0.03 | 0.846 | 0.092 | 0.055 | 0.003 | 0.001 | 0.003 | 2.907 | 1.016 | 0.949 |
| 0.92 | 0.06 | 0.02 | 0.846 | 0.110 | 0.037 | 0.004 | 0.000 | 0.002 | 2.922 | 1.021 | 0.954 |
| 0.92 | 0.07 | 0.01 | 0.846 | 0.129 | 0.018 | 0.005 | 0.000 | 0.001 | 2.937 | 1.026 | 0.959 |
| 0.93 | 0.04 | 0.03 | 0.865 | 0.074 | 0.056 | 0.002 | 0.001 | 0.002 | 2.899 | 1.013 | 0.946 |
| 0.93 | 0.05 | 0.02 | 0.865 | 0.093 | 0.037 | 0.003 | 0.000 | 0.002 | 2.914 | 1.018 | 0.951 |
| 0.93 | 0.06 | 0.01 | 0.865 | 0.112 | 0.019 | 0.004 | 0.000 | 0.001 | 2.929 | 1.024 | 0.956 |
| 0.94 | 0.04 | 0.02 | 0.884 | 0.075 | 0.038 | 0.002 | 0.000 | 0.002 | 2.907 | 1.016 | 0.949 |
| 0.94 | 0.05 | 0.01 | 0.884 | 0.094 | 0.019 | 0.003 | 0.000 | 0.001 | 2.922 | 1.021 | 0.954 |
| 0.95 | 0.03 | 0.02 | 0.903 | 0.057 | 0.038 | 0.001 | 0.000 | 0.001 | 2.899 | 1.013 | 0.947 |
| 0.95 | 0.04 | 0.01 | 0.903 | 0.076 | 0.019 | 0.002 | 0.000 | 0.001 | 2.914 | 1.018 | 0.951 |
| 0.96 | 0.03 | 0.01 | 0.922 | 0.058 | 0.019 | 0.001 | 0.000 | 0.001 | 2.907 | 1.016 | 0.949 |
