## Supplementary Figs. for "Lower promoter activity of the *ST8SIA2* gene has been favored in evolving human collective brains"

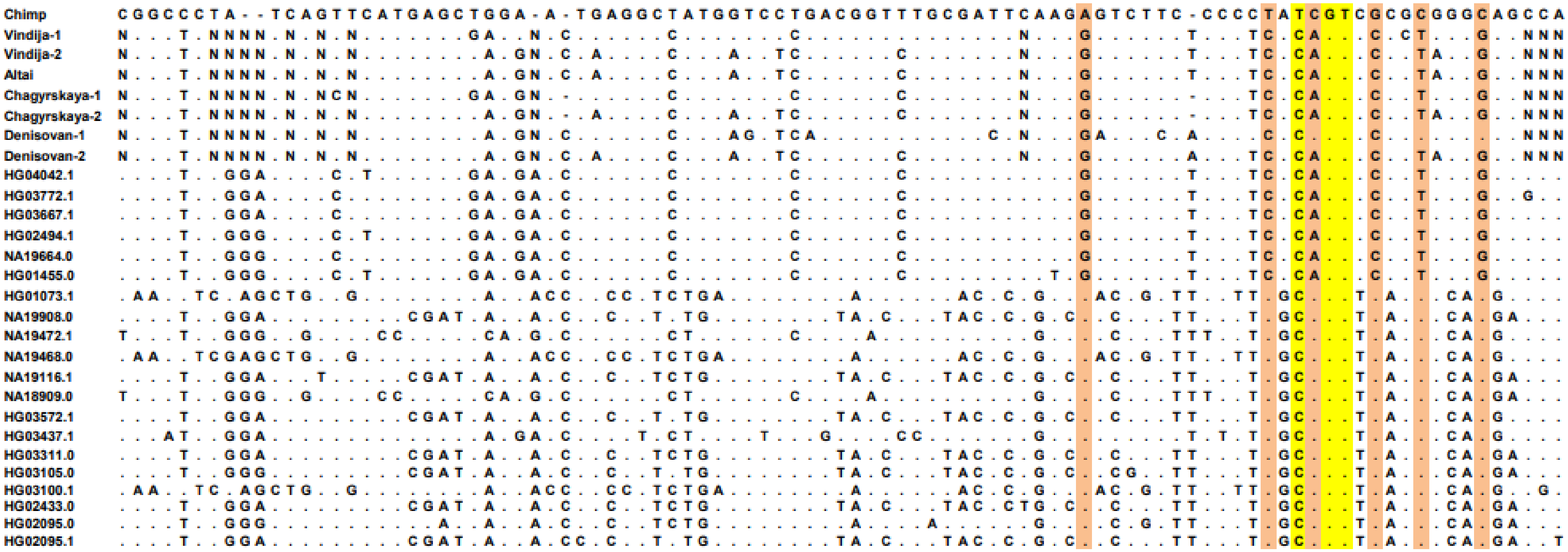


Supplementary Fig. 1. Sequence alignment of haplotypes containing three promoter SNPs of the *ST8SIA2* gene. The three promoter SNPs are highlighted in yellow. The “N”s represent uncertain sites. The name of the AMH haplotype is shown by the ID of the individual who has the sequence (phase 3 of the 1000 Genomes Project database [16]). The six SNPs used for the estimation of variant age are highlighted in orange.

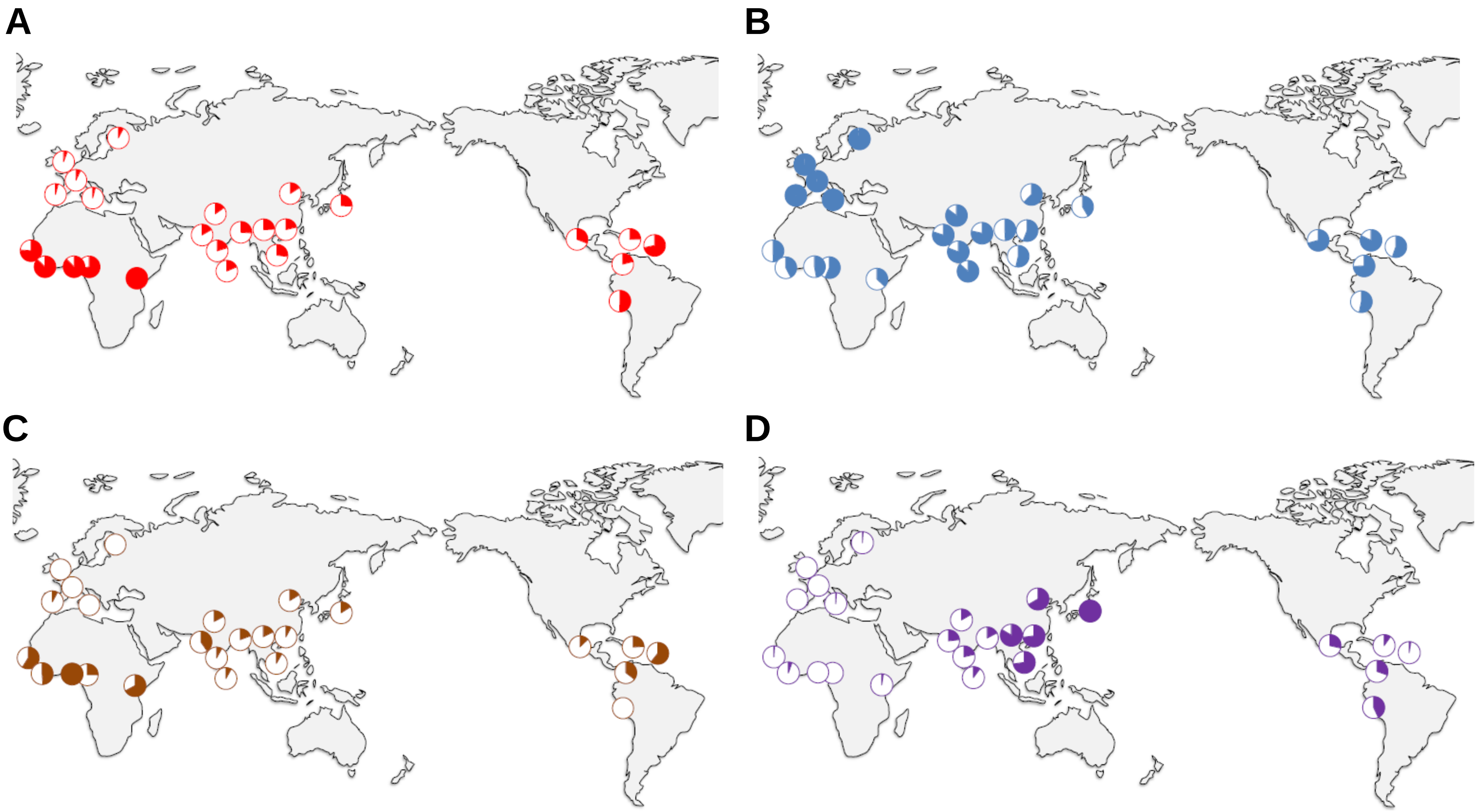


Supplementary Fig. 2. Global distribution of the TGT (A), TCT (B), CGT (C), and CGC (D) sequences. Pie charts represent relative frequencies as compared with the highest frequency among subpopulations [LWK in (A), TSI in (B), YRI in (C), and JPT in (D)]. ASW is not shown because of a lack of information about homelands in Africa.

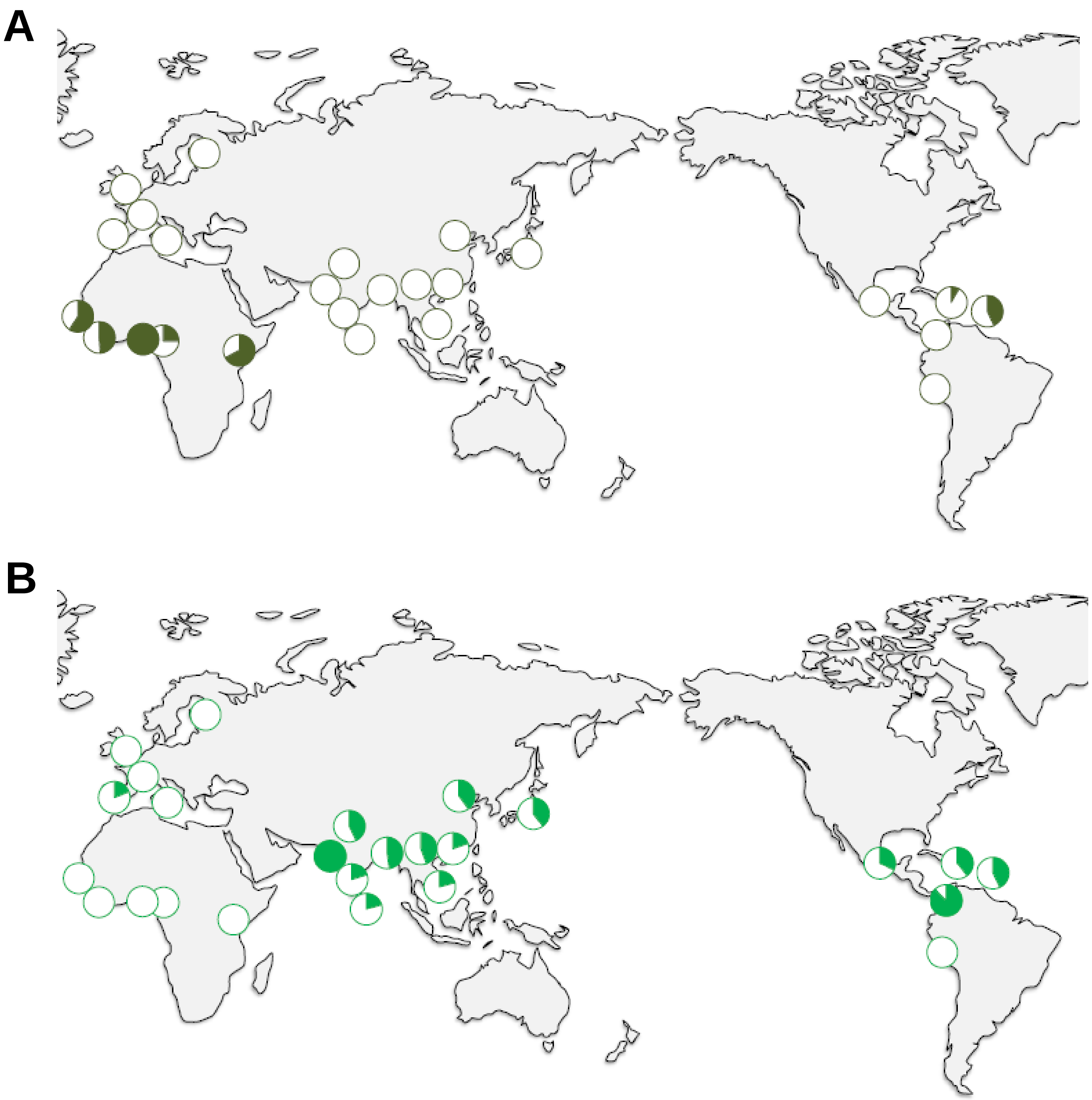


Supplementary Fig. 3. Global distribution of the CGT1 (A) and CGT2 (B) sequences. Pie charts represent relative frequencies as compared with the highest frequency among subpopulations [YRI in (A) and GIH in (B)]. ASW is not shown because of a lack of information about homelands in Africa.

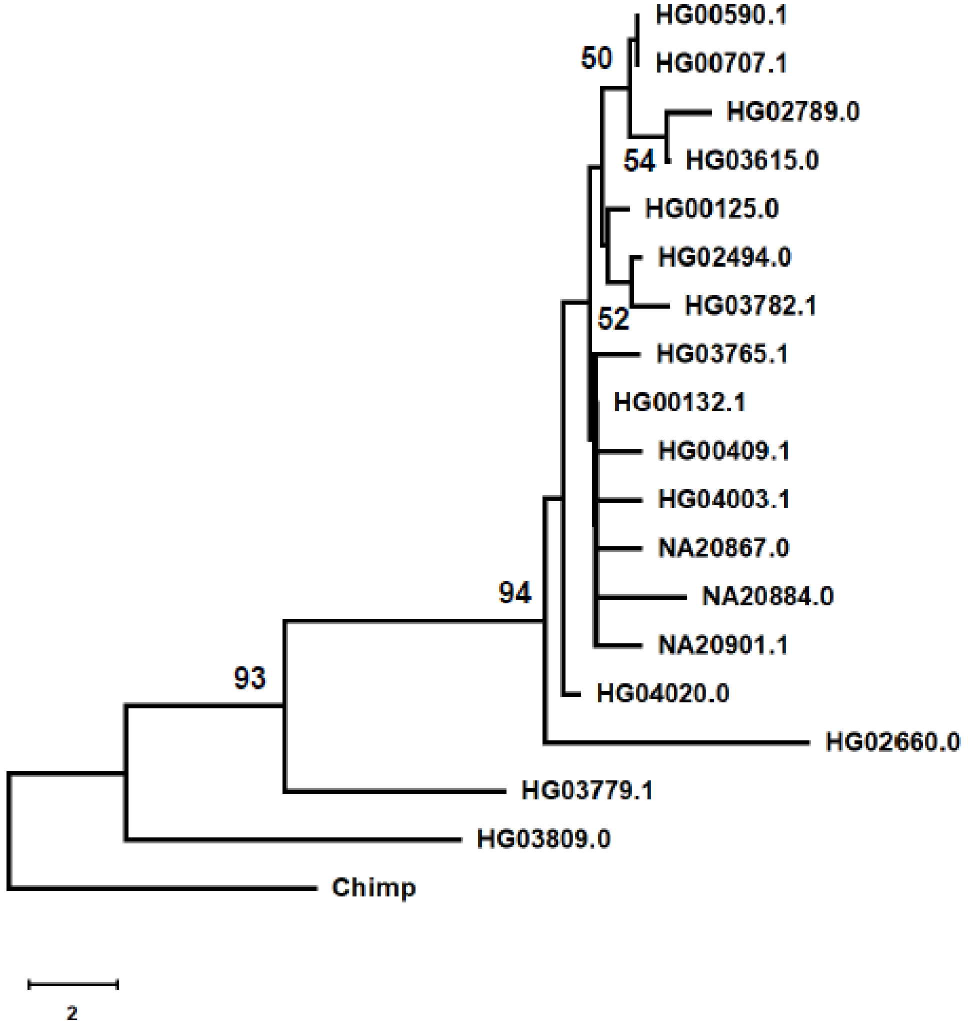


Supplementary Fig. 4. Tree of the TGT haplotypes from SAS. The phylogenetic tree was constructed by the Neighbor-Joining method with the number of differences. Bootstrap values of more than 50% from 1,000 replications are shown on the tree branches.

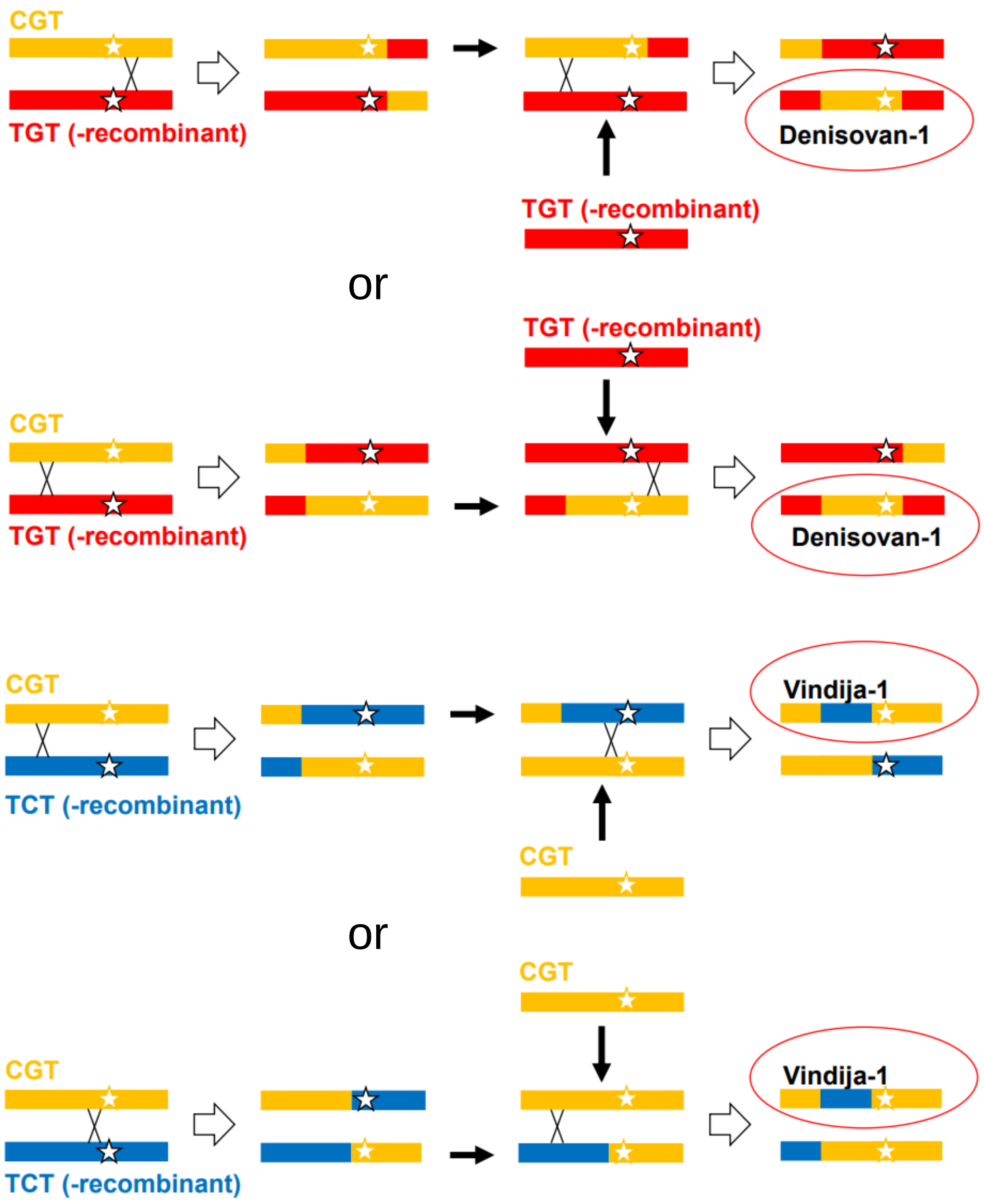


Supplementary Fig. 6. Recombination process producing Denisovan-1 and Vindija-1. Star represents the position of the three promoter SNPs.

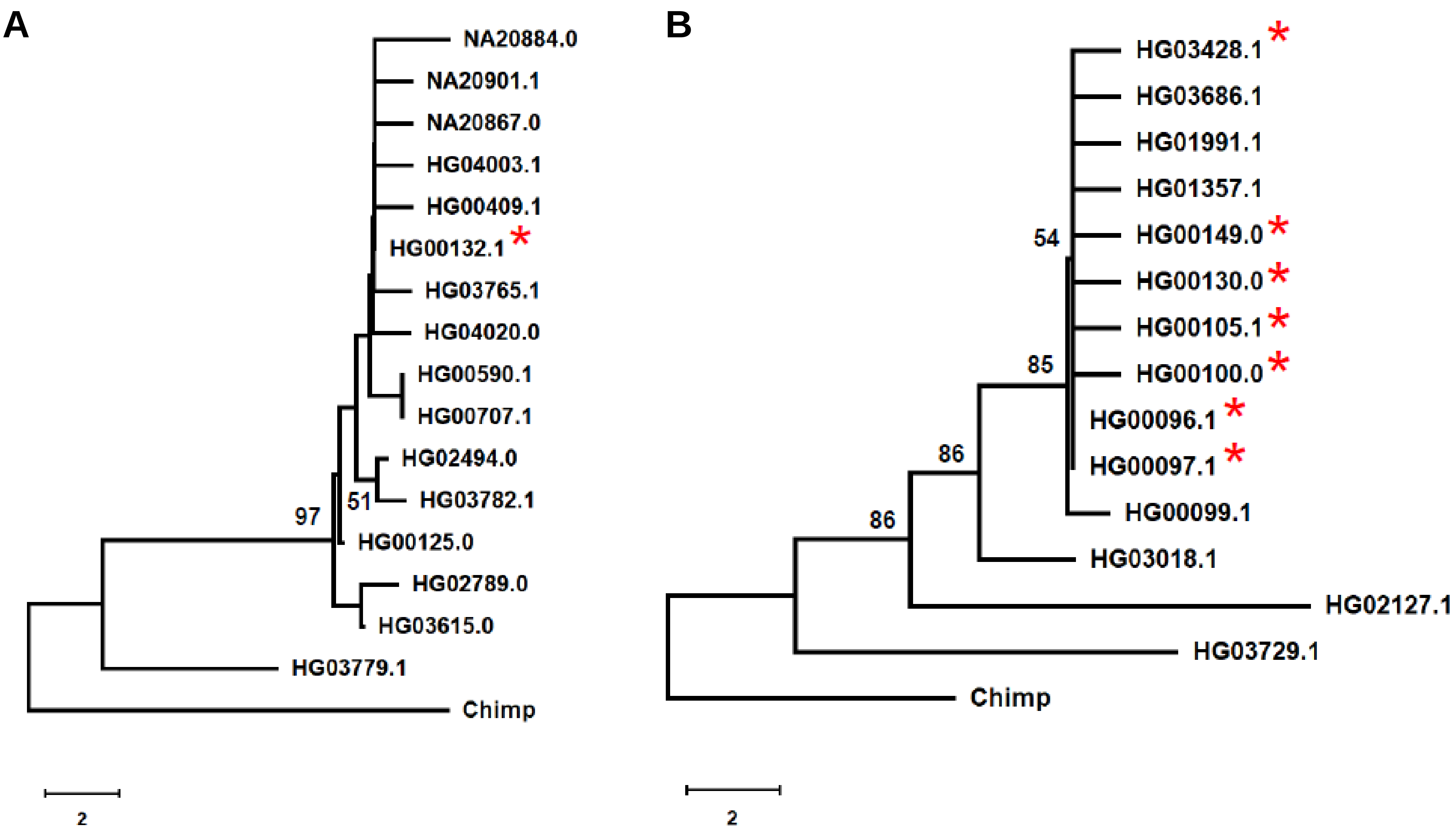


Supplementary Fig. 7. Haplotype trees of promoter types from SAS. (A) Tree of the TGT haplotypes. (B) Tree of the TCT haplotypes. These haplotypes were selected based on the analyses using site differences (see the "Materials and Methods" section). The phylogenetic trees were constructed by the Neighbor-Joining method with the number of differences. Bootstrap values of more than 50% from 1,000 replications are shown on the tree branches. Red star represents the haplotype found in AFR.

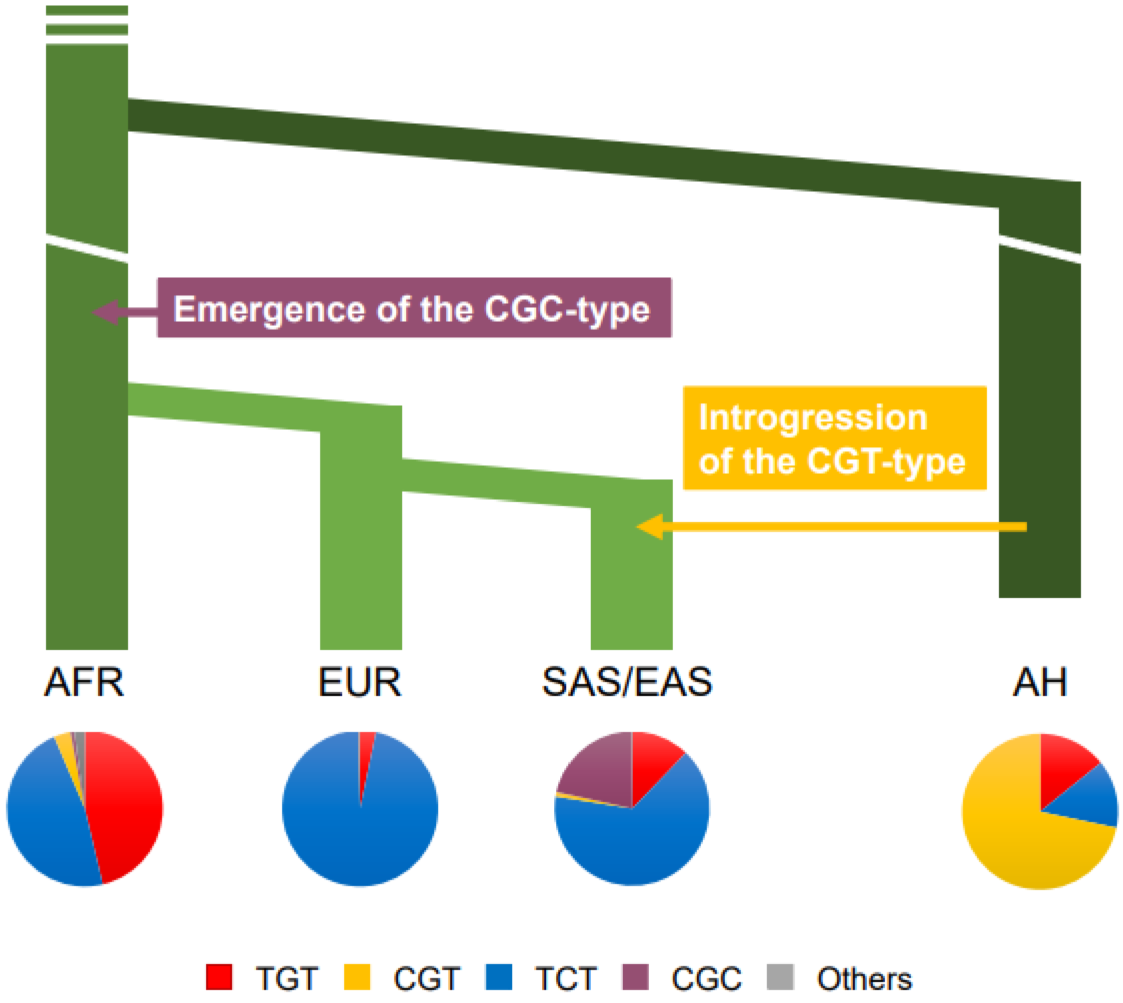


Supplementary Fig. 8. Evolution of promoter types in AMHs and AHs. Pie chart of AHs was drawn by promoter-type composition producing the lowest inferred PPA (see Supplementary Table 8).

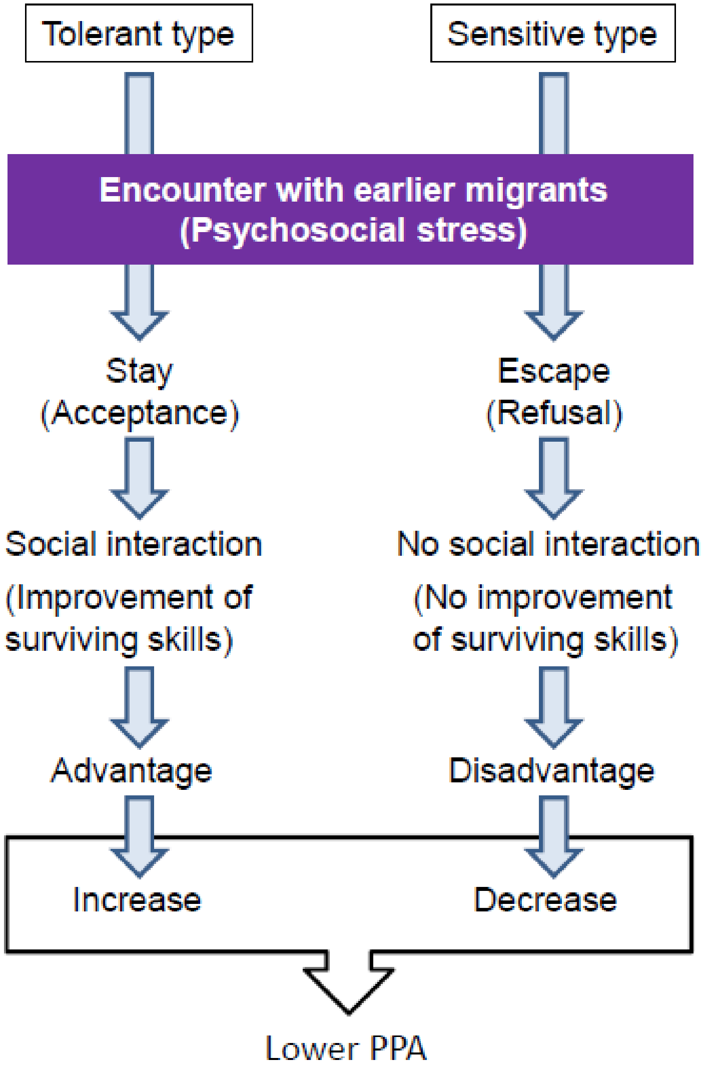


Supplementary Fig. 9. Hypothetical process of adaptive increase of tolerant types in the *ST8SIA2* gene. For details of the functional consequence on social interaction, please see our previous report [9].

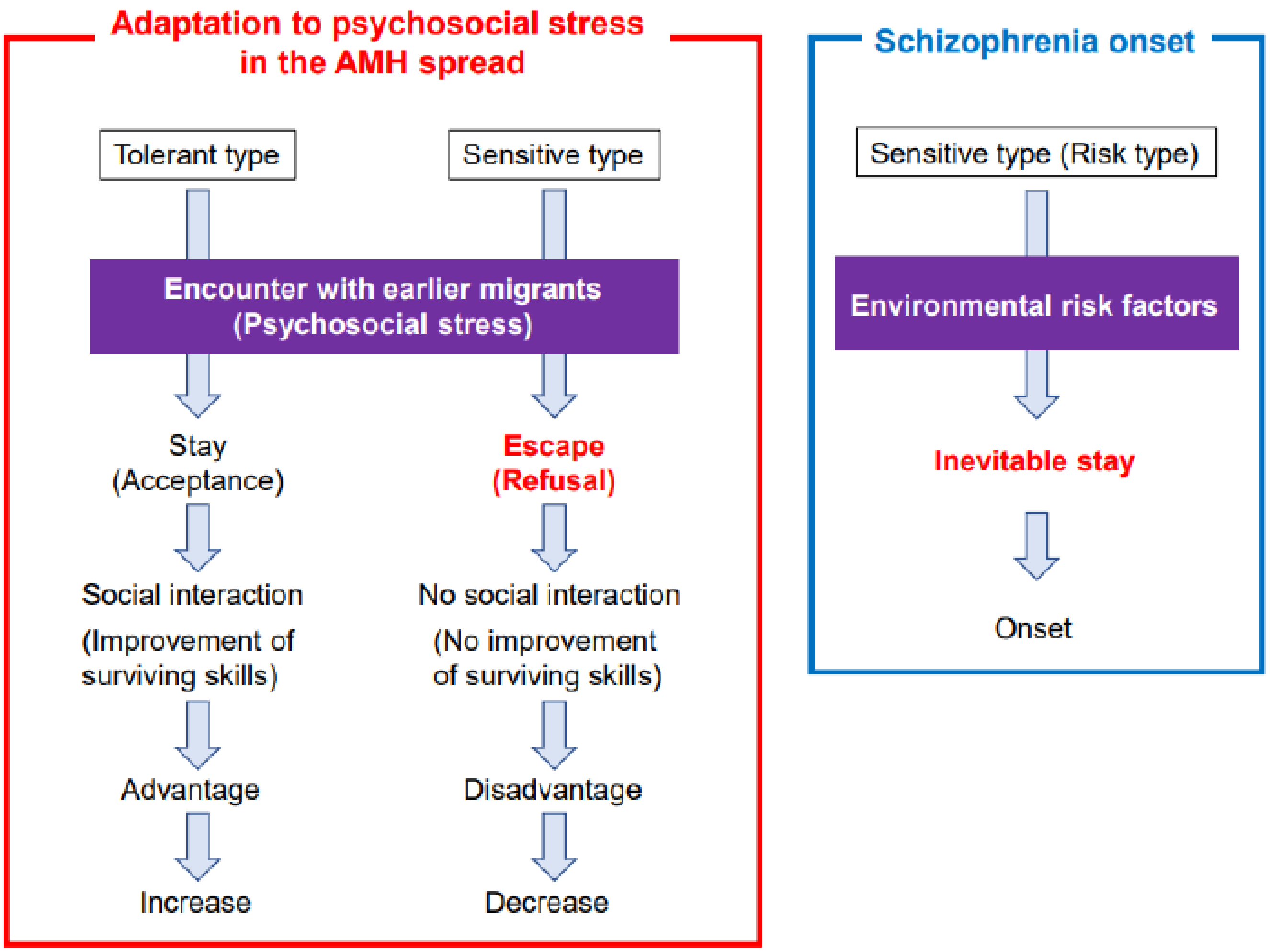


Supplementary Fig. 10. Conditional difference between adaptation to psychosocial stress and the onset of schizophrenia. In the onset of schizophrenia, inevitable stay under the environmental risk factors is essential. However, this inevitability of staying is not necessarily assumed in the adaptation to psychosocial stress.
